# A modular chemigenetic calcium indicator enables in vivo functional imaging with near-infrared light

**DOI:** 10.1101/2023.07.18.549527

**Authors:** Helen Farrants, Yichun Shuai, William C. Lemon, Christian Monroy Hernandez, Shang Yang, Ronak Patel, Guanda Qiao, Michelle S. Frei, Jonathan B. Grimm, Timothy L. Hanson, Filip Tomaska, Glenn C. Turner, Carsen Stringer, Philipp J. Keller, Abraham G. Beyene, Yao Chen, Yajie Liang, Luke D. Lavis, Eric R. Schreiter

## Abstract

Genetically encoded fluorescent calcium indicators have revolutionized neuroscience and other biological fields by allowing cellular-resolution recording of physiology during behavior. However, we currently lack bright, genetically targetable indicators in the near infrared that can be used in animals. Here, we describe WHaloCaMP, a modular chemigenetic calcium indicator built from bright dye-ligands and protein sensor domains that can be genetically targeted to specific cell populations. Fluorescence change in WHaloCaMP results from reversible quenching of the bound dye via a strategically placed tryptophan. WHaloCaMP is compatible with rhodamine dye-ligands that fluoresce from green to near-infrared, including several dye-ligands that efficiently label the central nervous system in animals. When bound to a near-infrared dye-ligand, WHaloCaMP1a is more than twice as bright as jGCaMP8s, and shows a 7× increase in fluorescence intensity and a 2.1 ns increase in fluorescence lifetime upon calcium binding. We use WHaloCaMP1a with near-infrared fluorescence emission to image Ca^2+^ responses in flies and mice, to perform three-color multiplexed functional imaging of hundreds of neurons and astrocytes in zebrafish larvae, and to quantitate calcium concentration using fluorescence lifetime imaging microscopy (FLIM).

## Introduction

Fluorescent indicators allow non-invasive imaging of cellular physiology in living animals, and genetically encoded fluorescent calcium indicators (GECIs) have been especially useful to approximate neuronal activity during behavior^1–3^. The most widely used GECIs are engineered from fluorescent proteins and emit in the green to red region of the visible spectrum. Near-infrared (∼670-900 nm) fluorophores can be imaged deeper in tissue due to reduced scattering and multiplexed with existing green and red probes^4^. However, there are limited functional indicators in the near infrared.

Existing near-infrared fluorescent indicators require either a cofactor or synthetic small molecule, which poses challenges for *in vivo* use^5^. Near-infrared calcium indicators that make use of biliverdin-binding proteins^6, 7^ are limited by the availability of biliverdin, a product of heme catabolism in mammalian cells, and have low quantum yields, making *in vivo* imaging with good signal-to-noise (SNR) challenging. While synthetic small-molecule dyes can be brighter than biliverdin-binding proteins, they must be delivered to the desired tissue. In addition, to achieve genetic targeting the small-molecule dyes must be selectively retained by a specific cell type, for example by using a self-labeling protein^8^. We previously engineered fluorescent indicators^9^ with far-red or near-infrared emission using environmentally sensitive dye-ligands, such as JF_635_-HaloTag ligand^10^, and the self-labeling protein HaloTag protein^9^ in a “chemigenetic” or “hybrid” approach. However, these dye-ligands exhibited limited bioavailability in the central nervous system and the resulting sensors could not be efficiently used for functional imaging in the brains of animals.

Here, we describe new chemigenetic calcium indicators built from bright dye-ligands whose fluorescence output is modulated by photoinduced electron transfer (PET) to a protein-based quencher. The modular approach allowed us to use rhodamine dye-ligands with emission ranging from green to near-infrared, including dye-ligands that efficiently distribute into the central nervous system in animals. We use these sensors to demonstrate imaging of Ca^2+^ responses in flies and mice and perform three color multiplexed functional imaging as well as quantitative Ca^2+^ fluorescence lifetime imaging microscopy (FLIM) in zebrafish larvae.

## Results

### Design of the chemigenetic calcium indicator WHaloCaMP

To construct a bright, near-infrared chemigenetic calcium sensor for use in animals, we focused on JF_669_-HaloTag ligand (Fig. 1a). This dye-ligand has emission just on the edge of the near-infrared and has previously been reported to efficiently label neurons in mouse brains following intravascular injection^11, 12^. We did not concentrate on environmentally sensitive dye-ligands such as JF_635_-HaloTag ligand, which we previously used to engineer the HaloCaMP chemigentic calcium indicators, as we could not detect the JF_635_-HaloTag ligand in the central nervous system of mice following vascular injection. Fluorescence changes in the HaloCaMP sensors relied on modulating the environment of the dyes via protein conformational changes driven by Ca^2+^-binding. The environmental sensitivity of rhodamine dyes is characterized by the equilibrium between the non-fluorescent lactone and the fluorescent zwitterionic forms. In previous work^10, 11, 13^, we found the lactone–zwitterion equilibrium constant (*K*_L–Z_) of a rhodamine dye is a descriptor that largely predicts both the bioavailability in the central nervous system and environmental sensitivity of dye-ligand derivatives (Supplementary Note 1). Unfortunately, the optimal range of *K*_L-Z_ for bioavailability shows little overlap with the optimal range for environmental sensitivity. Thus, the excellent bioavailability of JF_669_-HaloTag ligand is at the expense of environmental sensitivity; this dye ligand does not exhibit large Ca^2+^-dependent fluorescence changes when used with the previously published HaloCaMP sensors (Supplementary Fig. 1).

**Figure 1.**
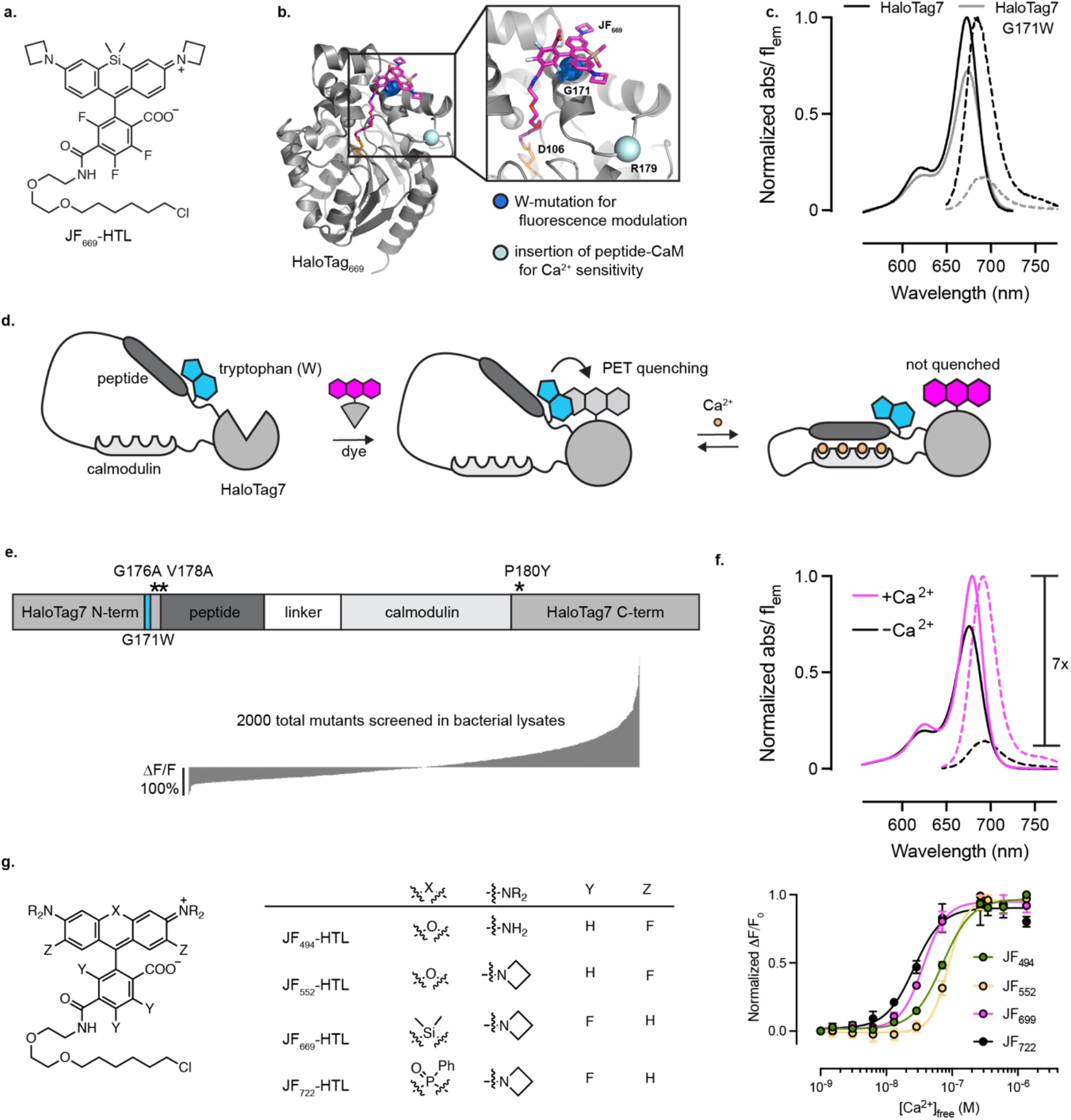
Engineering chemigenetic calcium indicators with tryptophan quenching. **a**, Chemical structure of JF_669_-HaloTag ligand (HTL). **b**, Crystal structure of HaloTag7 bound to JF_669_-HaloTag ligand (HaloTag_669_) (PDBID 8SW8). The position of G171, which was mutated to a tryptophan to quench dye fluorescence emission, and R179, where calcium sensitive protein domains were inserted, are highlighted as spheres. **c**, Normalized absorption (solid lines) and fluorescence emission (dashed lines) spectra of JF_669_-HaloTag ligand bound to HaloTag7 or HaloTag7 G171W mutant. **d**, Schematic representation of WHaloCaMP, showing domain arrangement, covalent binding of the dye-ligand, and the quenching tryptophan. **e**, Primary structure of WHaloCaMP1a (top) and ΔF/F_0_ of variants (bottom) from a bacterial lysate screen to select WHaloCaMP1a. **f**, Normalized absorption (solid lines) and fluorescence emission (dashed lines) spectra of JF_669_-HaloTag ligand bound to purified WHaloCaMP1a in the presence (magenta) and absence (black) of calcium. **g**, Chemical structures of dye-ligands used here with WHaloCaMP1a (left), and normalized Ca^2+^ titrations of WHaloCaMP1a bound to these dye-ligands (right).

Instead of tuning the lactone–zwitterion equilibrium to find a dye with mediocre bioavailbility and environmental sensitivity, we hypothesized that we could generate Ca^2+^ binding-induced fluorescence change via reversible quenching of the dye by tryptophan from the protein through photoinduced electron transfer (PET). Tryptophan mutants of HaloTag have previously been used to quench the fluorescence of rhodamine dyes^14^ and to improve the dynamic range of fluorescent indicators^15^. To guide our engineering efforts, we solved the crystal structure of JF_669_-HaloTag ligand bound to HaloTag7 (Fig. 1b, PDBID 8SW8) and found that G171 was within the required spatial distance of ∼5 Å for fluorescence quenching through PET^16^. Mutating G171 on HaloTag7 to a tryptophan decreased fluorescence emission of bound JF_669_-HaloTag ligand by 85% (Fig. 1c), providing a way to strongly modulate the dye fluorescence emission.

We next sought to couple the fluorescence quenching of HaloTag-bound JF_669_ to Ca^2+^-induced conformational changes in the protein (Fig. 1d, Supplementary Fig. 2). We explored different topologies of peptide-calmodulin (CaM) insertions into HaloTag_G171W_ at positions spatially proximal to the bound dye and screened for insertions that retained a high dye-ligand capture rate close to that of HaloTag7 (above 10^6^ M^-1^ s^-1^)^17^ to ensure good labeling with dye-ligand following bolus loading into complex tissue. Insertion of the myosin light chain kinase (MLCK) CaM-binding peptide followed by CaM at position R179 of HaloTag7_G171W_ resulted in a protein with a fast dye-ligand capture rate and a modest Ca^2+^-dependent change in fluorescence (Supplementary Fig. 3 and 4). We then performed three rounds of directed evolution on this scaffold with single and double site saturation mutagenesis at positions close to the insertion site and the dye-protein interface (Fig. 1e, Supplementary Fig. 5). In total we screened ∼2000 variants in bacterial lysates and validated 10 in primary rat hippocampal neuronal culture. We named the variant with the best performance WHaloCaMP1a: it has the fastest dye-ligand capture rate, largest change in fluorescence intensity on addition of Ca^2+^, reasonable Ca^2+^ kinetics (Supplementary Fig. 6), low one-photon bleaching rate in the absence of Ca^2+^ (Supplementary Fig. 7), and best performance in neuronal culture. We chose the name WHaloCaMP as the one-letter abbreviation for tryptophan is (W), HaloTag is used to localize the dye-ligand to the protein sensor (Halo), and it is a Calcium-Modulated Protein (CaMP).

WHaloCaMP1a_669_ showed a small increase in absorbance (1.4×) upon binding Ca^2+^, but a larger increase in fluorescence quantum yield (6.2×), which produce an overall Ca^2+^-induced fluorescence emission increase of 7× at 690 nm (Fig. 1f and Table 1). WHaloCaMP1a_669_ is more than twice as bright as jGCaMP8s^18^, and 40× brighter than the brightest biliverdin-based near-infrared calcium indicator reported to date, iGECI^7^ (Supplementary Fig. 8), in the Ca^2+^-bound state, and shows favorable maximum fluorescence change, Ca^2+^ affinity, and kinetics of fluorescence change. WHaloCaMP1a exhibits increased fluorescence emission upon the addition of Ca^2+^, making it a “positive-going” indicator, while iGECI and most biliverdin-binding calcium indicators show decreased fluorescence upon addition of Ca^2+^.

**Table I.**
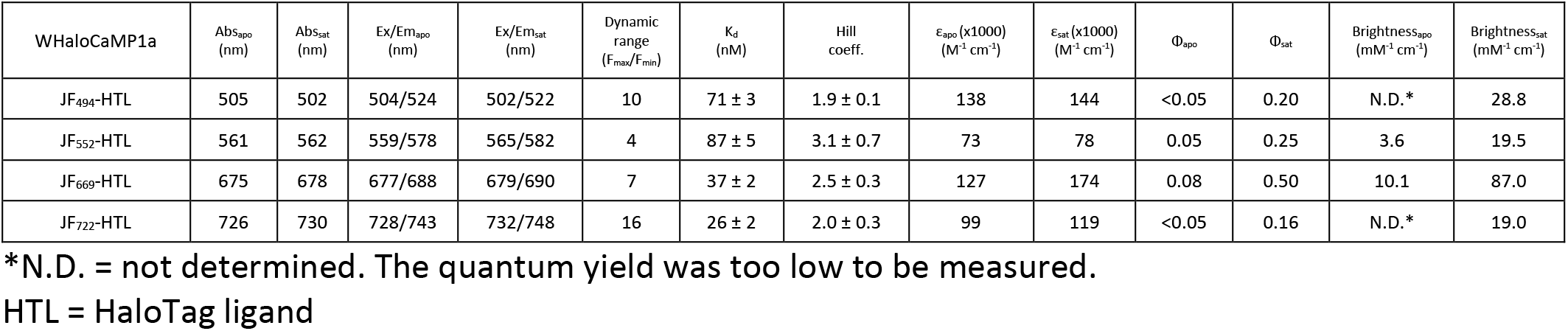
Photophysical properties of WHaloCaMP1a bound to dye-ligands.

WHaloCaMP proved to be amenable to further modification of its properties based on previous GECI engineering efforts. We decreased the Ca^2+^ affinity by single point mutations in the MLCK CaM-binding peptide^19^ (Supplementary Fig. 9) and produced faster response kinetics by substituting the CaM-binding peptide with endothelial nitric oxide synthase peptide (“ENOSP”), previously used in the jGCaMP8 series^18^ (Supplementary Fig. 10). We also explored other topologies of WHaloCaMP. Several protein engineering efforts^20, 21^ have focused on a loop close to the dye binding site at position T155 of HaloTag7. We generated WHaloCaMP1b via insertion of MLCK-calmodulin (CaM) at position T154 and mutation of A151 to tryptophan (Supplementary Fig. 11). We further characterized WHaloCaMP1a in this work since it showed larger fluorescence changes, but we note that the WHaloCaMP1b topology also appears to be a reasonable solution to generating chemigenetic indicators.

A benefit of chemigenetic indicators is that the color of fluorescence emission of the indicator can be altered by changing the small-molecule dye-ligand. In addition to the near-infrared emitting JF_669_-HaloTag ligand, we explored WHaloCaMP1a bound to other rhodamine-based dyes, including the green emitting JF_494_-HaloTag ligand^22^, the orange emitting JF_552_-HaloTag ligand^13^ and the near-infrared emitting JF_722_-HaloTag ligand^11^ (Fig 1g. and Supplementary Fig. 12, 13, 14). Consistent with a tryptophan PET quenching mechanism, fluorescence changes were mainly conferred by changes in quantum yield of each bound dye (Table 1). WHaloCaMP1a displayed a high Ca^2+^ affinity (26 ± 2 nM to 71 ± 3 nM for different dyes, Fig. 1g. and Table 1), similar to that of jGCaMP8s (46 ± 1 nM)^18^. The WHaloCaMP1a sensor bound to different dye-ligand showed Ca^2+^-dependent fluorescence changes from pH 5 to 9. WHaloCaMP1a_669_ was particularly stable, exhibiting little change in Ca^2+^ response in the pH range of 6 to 8 (Supplementary Fig. 15). The relative pH insensitivity indicates that WHaloCaMP1a could be used in a wide range of cellular compartments.

### WHaloCaMP characterization in rodent neuron cultures and acute brain slices

We expressed WHaloCaMP1a in primary rat hippocampal neurons in culture, labeled with dye-ligands, and elicited action potentials (APs) evoked by electrical stimulation with a field electrode (Fig. 2a). WHaloCaMP1a could detect single APs with a ΔF/F_0_ of up to 20% for WHaloCaMP_494_ (Fig. 2b and Supplementary Fig. 16). WHaloCaMP_722_ is the furthest red-shifted chemigenetic fluorescent indicator that can follow single action potential transients in neurons reported to date. WHaloCaMP1a_669_ showed 2.3× larger max ΔF/F_0_ in neurons than the brightest biliverdin-binding calcium indicator, iGECI (max ΔF/F_0_ = 77% for WHaloCaMP1a_669_ vs. -30 % for iGECI^7^, Fig. 2c and Supplementary Fig. 17). In addition, WHaloCaMP1a_669_ shows faster time to peak and half decay time compared to iGECI (time to peak at 1 AP = 176 ms vs 700 ms and weighted average half time decay for 1 AP = 2 s vs 14 s), making it better suited to capture neuronal activity (Supplementary Fig. 18 and 19).

**Figure 2.**
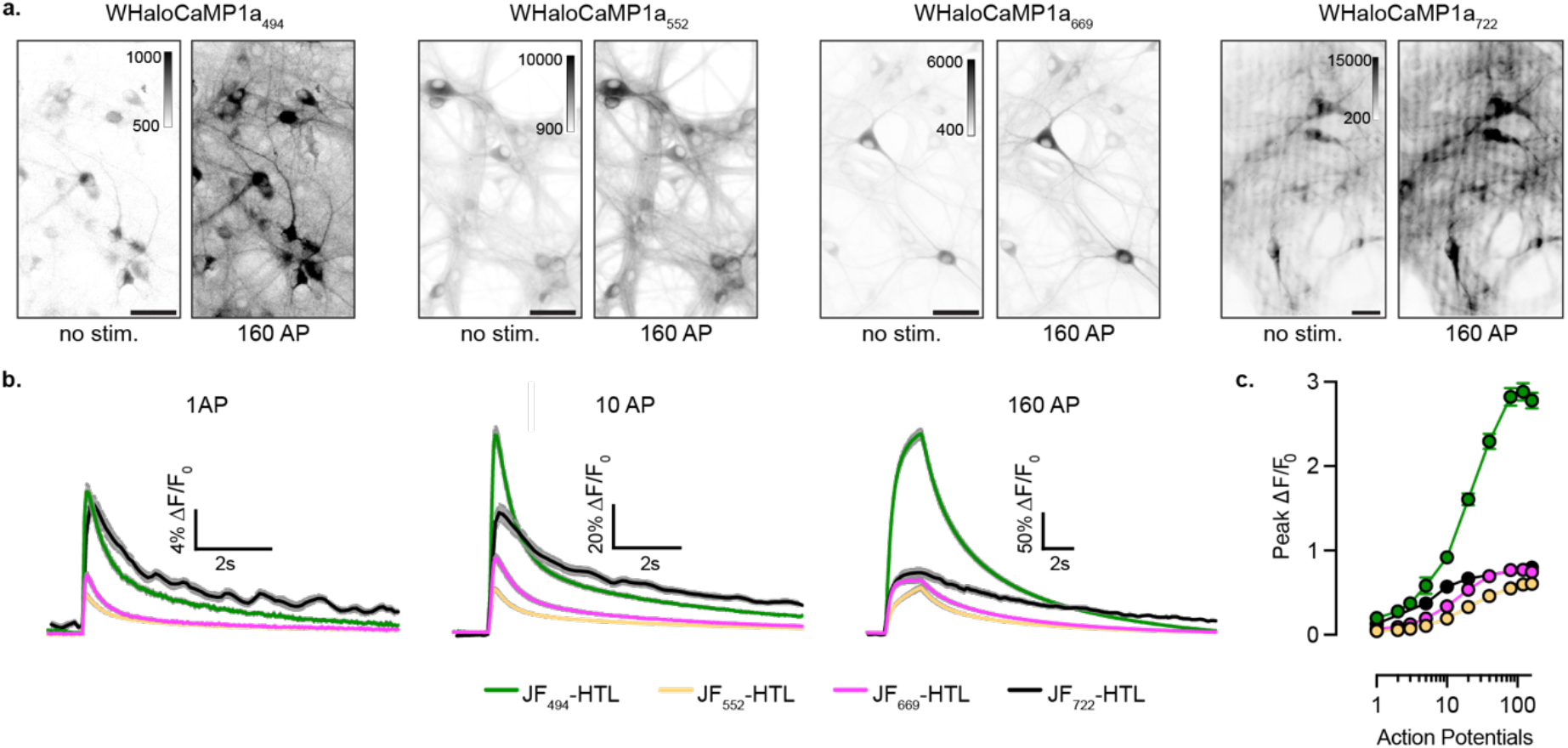
Characterization of WHaloCaMP1a in neuronal cultures. **a**, Representative images of cultured rat hippocampal neurons expressing WHaloCaMP1a labeled with dye-ligands unstimulated or stimulated with 160 induced action potentials (APs) at 80 Hz Scale bars, 50 µm. **b,** ΔF/F_0_ response of WHaloCaMP1a expressed in cultured rat hippocampal neurons and labeled with the indicated dye-ligands to trains of APs at 80 Hz. Solid line (mean) and grey outline (s.e.) for n = 130-165 neurons for JF_494_-HTL, JF_552_-HTL, JF_669_-HTL and n = 20 for JF_722_-HTL. **c,** Peak ΔF/F_0_ as a function of the number of APs at 80 Hz. Data are presented as mean and s.e. for n = 130-165 neurons for JF_494_-HTL, JF_552_-HTL, JF_669_-HTL and n = 20 for JF_722_-HTL.

The excitation spectrum of WHaloCaMP1a_669_ is better separated from the action spectrum of many blue-excited optogenetic tools than red calcium indicators. We found via electrophysiology that the light required for exciting WHaloCaMP1a_669_ did not produce any observable membrane depolarization in neurons expressing the sensitive channelrhodopsin variant CheRiff^23^ (Supplementary Fig. 20). Indeed, even >10× the light power needed for functional imaging with WHaloCaMP1a_669_ did not produce any depolarization. In contrast, the light required for imaging the commonly used red FP-based calcium indicator jRCaMP1a produced substantial subthreshold depolarization of neurons expressing CheRiff.

The near-infrared emission of WHaloCaMP1a_669_ also allows multiplexing with current fluorescent protein-based indicators. We performed simultaneous multiplexed two-photon imaging of fluctuating Ca^2+^ and protein kinase A (PKA) activity in response to muscarinic acetylcholine receptor and adenylate cyclase activation in acute mouse brain slices using WHaloCaMP1a_669_ and a green PKA activity indicator, FLIM-AKAR^24^ (Supplementary Fig. 21). Consistent with previous results with single reporter imaging^25^, we observed concurrent calcium and PKA activity increase in response to mAChR activation, and only increased PKA phosphorylation in response to adenylate cyclase activation. Because calcium increase is upstream of PKA activation, these results demonstrate the possibility to track concurrent dynamics of multiple signals to understand temporal transformations in a signaling transduction cascade.

### WHaloCaMP1a reports neuronal calcium responses in flies and mice

Since previous chemigenetic HaloTag-based sensors, such as HaloCaMPs, were limited to experiments in reduced preparations, we explored whether WHaloCaMP1a could be expressed, labeled with dye-ligands, and used to follow physiological Ca^2+^ responses in living animals. First, we tested whether WHaloCaMP1a could report odor-evoked Ca^2+^ responses in the mushroom body Kenyon cells (KCs) in adult *Drosophila* (Fig. 3a). KCs are third-order neurons in the fly olfactory pathway, and their odor response profile has been extensively characterized^26^. We expressed WHaloCaMP1a in KCs with R13F02-Ga4. We then opened a small window in the head capsule of flies to expose the calyx that houses the dendritic processes of all the KCs (Fig. 3b), and labeled WHaloCaMP1a with JF_669_-HaloTag ligand by application in saline solution. Upon odor presentation, we observed clear WHaloCaMP1a_669_ fluorescence increases in individual trials in response to apple cider vinegar as well as two chemical odors (Fig. 3b). Similar responses were observed when we labeled WHaloCaMP with JF_552_-HaloTag ligand, highlighting the spectral flexibility of the indicator. Trends in odor response were similar to that of the RFP-based calcium indicator jRGECO1a^27^ (Supplementary Fig. 22).

**Figure 3.**
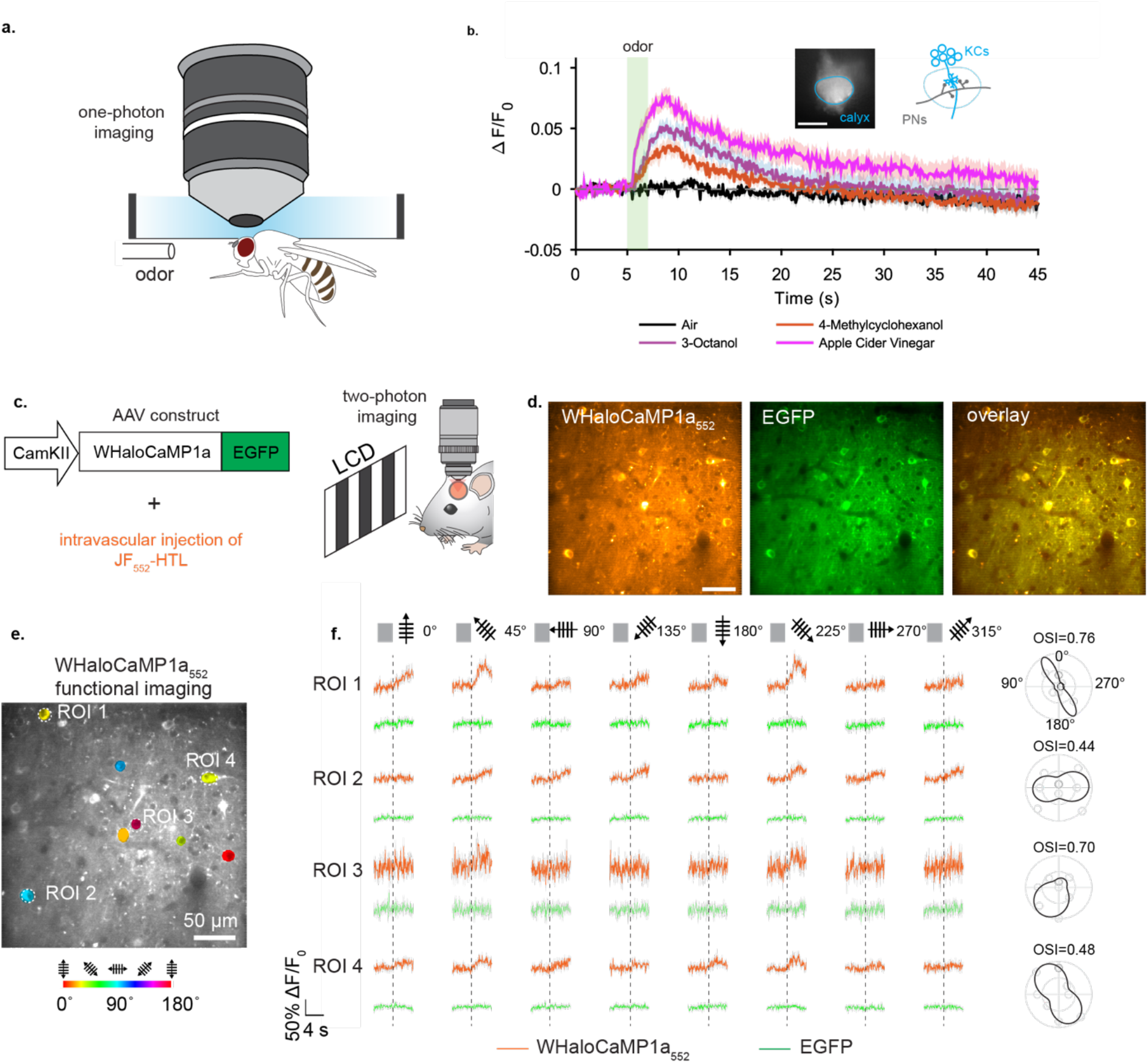
WHaloCaMP1a reports on neuronal activity in flies and mice **a**, One-photon imaging set-up of head-fixed flies expressing WHaloCaMP1a labeled with dye-ligands. **b**, Fluorescence responses from WHaloCaMP1a_669_ in head-fixed flies presented with different odors. WHaloCaMP1a was expressed in mushroom body Kenyon cells (KCs). Images were acquired from the calyx, where KCs receive dendritic inputs from the olfactory projection neurons (PNs) (insets). Green shading indicates odor presentation for 2 s. Data were from 6 flies, and odors were presented three times for each fly. The thick line and shaded areas indicate means and s.e.m. across odor trials. Scale bar, 50 µm. **c**, AAV construct for transducing neurons in mouse V1 and the schematic of the experimental setup for two-photon functional imaging of WHaloCaMP1a in the visual cortex of mice. JF_552_-HTL was intravascularly injected one day before examining orientation selectivity of V1 neurons in the anesthetized mouse exposed to moving grafting visual stimuli of different orientations and directions. **d**, Representative images of a field-of-view in mouse V1 showing neurons expressing WHaloCaMP1a_552_ or EGFP. Scale bar, 50 µm. **e**, Functional imaging of V1 neurons shows the orientations selectivity map. Scale bar, 50 µm. **f**, Functional imaging of Ca^2+^ (WHaloCaMP1a_552_ channel) or control (EGFP channel) traces (ROI1-4) in response to drifting gratings in directions show above the traces. Orientation selectivity index (OSI) of cells is shown in the right panel. Imaging rate was 15 Hz. Representative imaging session from three imaging sessions shown.

We then tested the performance of WHaloCaMP1a in mice. We expressed WHaloCaMP1a with an EGFP expression marker from the Calcium–calmodulin (CaM)-dependent protein kinase II (CaMKII) promoter in mouse primary visual cortex (V1) using adeno-associated virus (AAV) mediated gene delivery (Fig. 3c). Intravenous delivery of JF_552_-HaloTag ligand or JF_669_-HaloTag ligand via retro-orbital injection successfully labeled WHaloCaMP1a expressed in V1 excitatory neurons (Fig. 3d and Supplementary Fig. 23 and 24). We observed clear visually evoked Ca^2+^ transients in layer 2/3 excitatory neurons with WHaloCaMP1a_669_ using 1225 nm two-photon excitation in single trials (Supplementary Fig. 23). Since most standard two-photon microscopes cannot achieve substantial 1225 nm excitation we instead labeled WHaloCaMP1a with JF_552_-HaloTag ligand and imaged at either 850 nm or 1050 nm. Using 850 nm we could co-excite the EGFP expression marker and cleanly separate the fluorescence emission, providing a non-responsive normalization channel during two-photon functional imaging, while 1050 nm selectively excited WHaloCaMP1a_552_ (Supplementary Fig. 24). We performed functional imaging during presentation of drifting grating visual stimuli to the mouse’s contralateral eye and observed responses in individual trials up to 35 % ΔF/F_0_, as well as orientation tuning of the responses that is a hallmark of V1 excitatory neurons (Fig. 3e and 3f and Supplementary Fig. 25).

### Functional imaging in zebrafish larvae with WHaloCaMP1a_669_

We generated transgenic zebrafish expressing WHaloCaMP1a under the pan-neuronal *elavl3* promoter. First, we verified that we could perform whole brain functional neuronal imaging with larvae at 4 to 5 days post fertilization (d.p.f.) with WHaloCaMP1a_669_ using light sheet imaging with excitation at 685 nm at 4 Hz (Supplementary Fig. 26 and Supplementary Movie 1). To benchmark the performance of WHaloCaMP1a_669_ against known red calcium indicators^27^, we crossed the *elavl3*:WHaloCaMP1a zebrafish line with a line expressing pan-neuronal jRGECO1b under the same *elavl3* promoter. We performed dual color light sheet imaging (excitation at 561 nm and 640 nm) from a single plane in the hindbrain in 4 to 5 d.p.f. zebrafish larvae. WHaloCaMP1a_669_ reported all the Ca^2+^ transients observed with jRGECO1b in a region of interest (ROI) in the hindbrain, albeit with lower max ΔF/F_0_ compared to the jRGECO1b signal (66% of the max ΔF/F_0_ signal of jRGECO1b, Supplementary Fig. 27).

### Multiplexed imaging in zebrafish larvae with WHaloCaMP1a_669_

Clean spectral separation of WHaloCaMP1a_669_ from green and red fluorescent proteins enabled us to perform three color multiplexed imaging in zebrafish larvae (Fig. 4a). To simultaneously follow three physiological signals, we crossed the transgenic line expressing pan-neuronal WHaloCaMP1a with a previously established line^28^ expressing the green-emitting glucose sensor iGlucoSnFR throughout the body of the fish using the β-actin promoter and the red-emitting calcium indicator jRGECO1a in skeletal muscle using the α-actinin promoter. After labeling WHaloCaMP1a with JF_669_-HaloTag ligand, we performed light sheet imaging of a single plane in the fish (Fig. 4b) with clear spectral separation of the fluorescence emission of the three sensors (Fig. 4c, Supplementary Fig. 28 and Supplementary Movie 2). Ca^2+^ transients in the zebrafish hindbrain neurons recorded using WHaloCaMP1a_669_ correlated with Ca^2+^ transients in skeletal muscle recorded using jRGECO1a, but neuronal signals in other parts of the brain, such as the optical tectum, did not (Fig. 4d). Glucose changes in muscle observed using iGlucoSnFR were substantially slower than either the neuronal or muscle Ca^2+^ transients as previously reported^28^.

**Figure 4.**
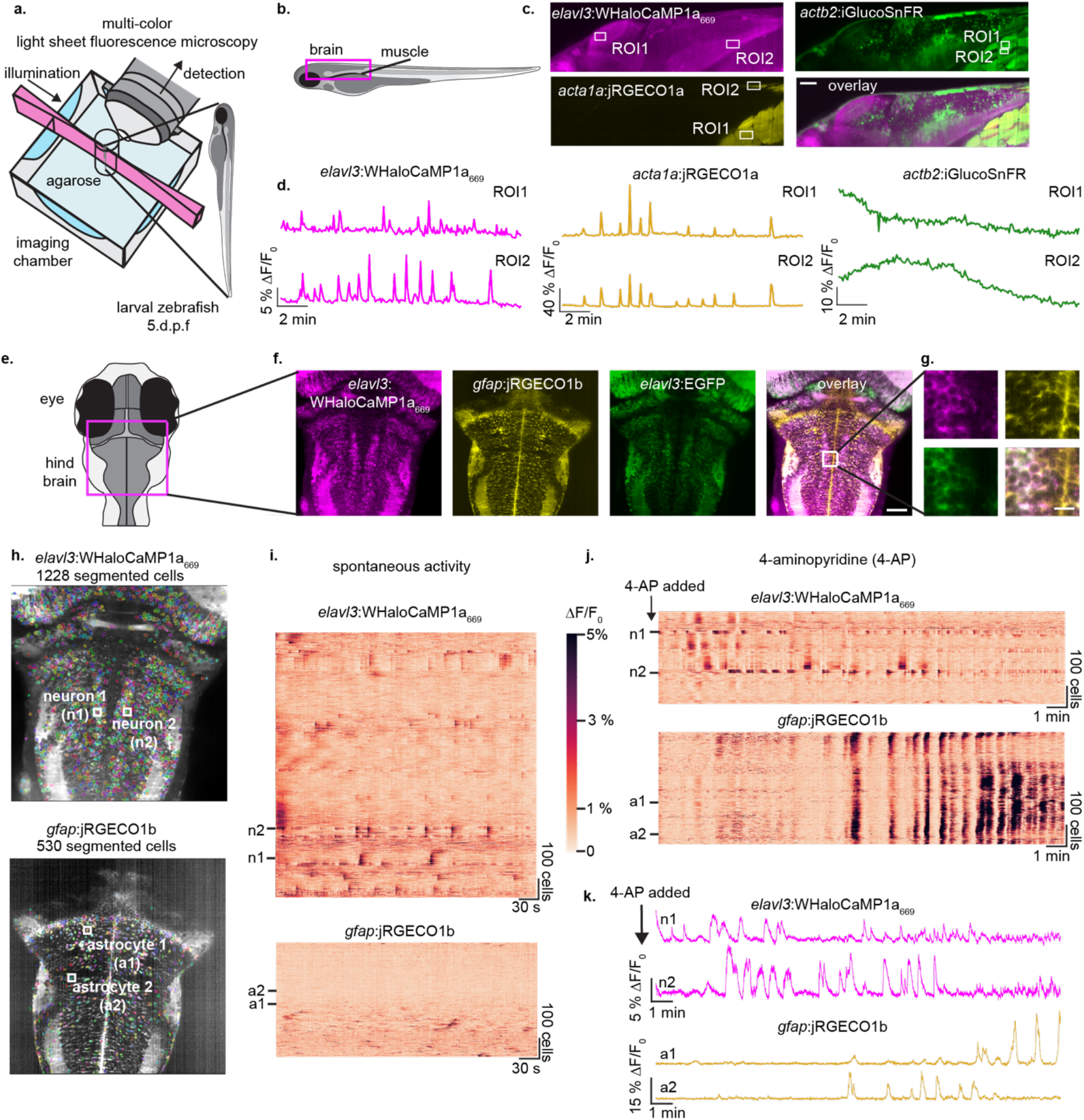
Three color multiplexed functional imaging in zebrafish larvae. **a**, Light sheet imaging set-up for multiplexed imaging. **b**, Schematic of side-view zebrafish larvae highlighting field of view for three color multiplexed functional imaging of glucose and Ca^2+^ in muscles and neurons. **c,** Representative images of WHaloCaMP1a expressed in neurons via the *elavl3* promoter, iGlucoSnFR expressed from the *actb2* ubiquitous promotor and jRGECO1a expressed in muscle via the *acta1a* promotor. Scale bar, 50 µm. **d**, Fluorescence ΔF/F_0_ traces of WHaloCaMP1a_669_, jRGECO1a and iGlucoSnFR in ROIs outlined in **b**. Representative experiment from 3 zebrafish larvae imaged. **e**, Schematic of the zebrafish larvae’s head indicating the field-of-view for light sheet imaging of neuronal and astrocyte activity. **f**, representative images of the expression patterns of WHaloCaMP1a_669_-EGFP expressed under the *elavl3* promoter, jRGECO1b expressed under the *gfap* promotor. Scale bar, 50 µm. **g**, Zoomed in images showing single cell resolution of fluorescent signals in the hind brain. Scale bar, 20 µm. **h**, Images of suite2p and Cellpose segmented cells from simultaneous functional imaging of WHaloCaMP1a_669_ and jRGECO1b. **i**, Rastermaps of activity from 1228 segmented neurons (top) and 530 astrocytes (bottom) during spontaneous brain activity. Two neurons (n1 and n2) indicate the hind brain oscillator. Two astrocytes (a1 and a1) are also indicated. **j**, 4-Aminopyridine (4-AP) was added to the imaging chamber of the zebrafish larva imaged in **i**, and functional imaging was performed. Concatenation of three imaging blocks of 6.2 minutes each. **k**, Fluorescence ΔF/F_0_ traces of n1 and n2 (top) and a1 and a2 (bottom) after addition of 4-AP.

Next, we asked if WHaloCaMP could be used to follow changes in neuronal activity to report changes in physiological states during multiplexed imaging. Activity of neurons and radial astrocytes are both linked to changes in behavior^29^, as reported by dual color functional imaging. We imaged zebrafish larvae expressing WHaloCaMP1a_669_ in neurons, and the red calcium indicator jRGECO1b in radial astrocytes, thus freeing up the green fluorescence emission channel, here using it as a fluorescent marker for neurons (Fig. 4e and f). Using light sheet imaging we could clearly resolve single cells in a plane of the zebrafish hindbrain (Fig. 4g). We then performed functional imaging of paralyzed zebrafish larvae at 4 Hz and used suite2p^30^ together with Cellpose^31^ (Fig. 4h and Supplementary Fig. 29) to segment and follow the activity of over one thousand neurons and hundreds of astrocytes simultaneously during spontaneous activity (Fig. 4i and Supplementary Movie 3). Without any external perturbation, WHaloCaMP1a_669_ clearly showed Ca^2+^ transients in a population of neurons that form part of the hindbrain oscillator^32^ related to motor control during swimming (n1 and n2 in Fig. 4i). Leading up to and during seizure-like states, induced by the potassium channel blocker 4-aminopyridine (4-AP), the pattern of neuronal activity in the hindbrain oscillator was lost (n1 and n2 in Fig. 4j and k, and Supplementary Movie 4), which has previously been described to be due to hyperexcitability^33^ of the neurons. We also observed large Ca^2+^ waves in the astrocyte population in response to 4-AP (a1 and a2 in Fig. 4j and k), as previously described^34^, which we could clearly separate from the neuronal activity, as the emission of jRGECO1b and WHaloCaMP1a_669_ are 100 nm apart. The loss of hindbrain oscillator activity and the large astrocytic Ca^2+^ waves were not observed in fish not treated with 4-AP (Supplementary Fig. 30). WHaloCaMP1a_669_ is well-suited for one-photon multicolor experiments in living animals as it can be excited with common far-red and near-infrared laser lines, and the emission is separated from commonly used fluorescent proteins.

### Fluorescence lifetime imaging microcopy (FLIM) using WHaloCaMP1a

The fluorescence modulation in WHaloCaMP1a on binding to Ca^2+^ is mainly due to changes in quantum yield. We thus explored WHaloCaMP1a as a fluorescence lifetime imaging microscopy (FLIM) probe for Ca^2+^ (Fig. 5a). Fluorescence lifetime is an inherent property of fluorophores, and FLIM allows absolute concentrations to be determined using fluorescent biosensors^35, 36^. However, the technique is limited by the number of photons that can be collected for accurate fitting of fluorescent lifetimes.

**Figure 5.**
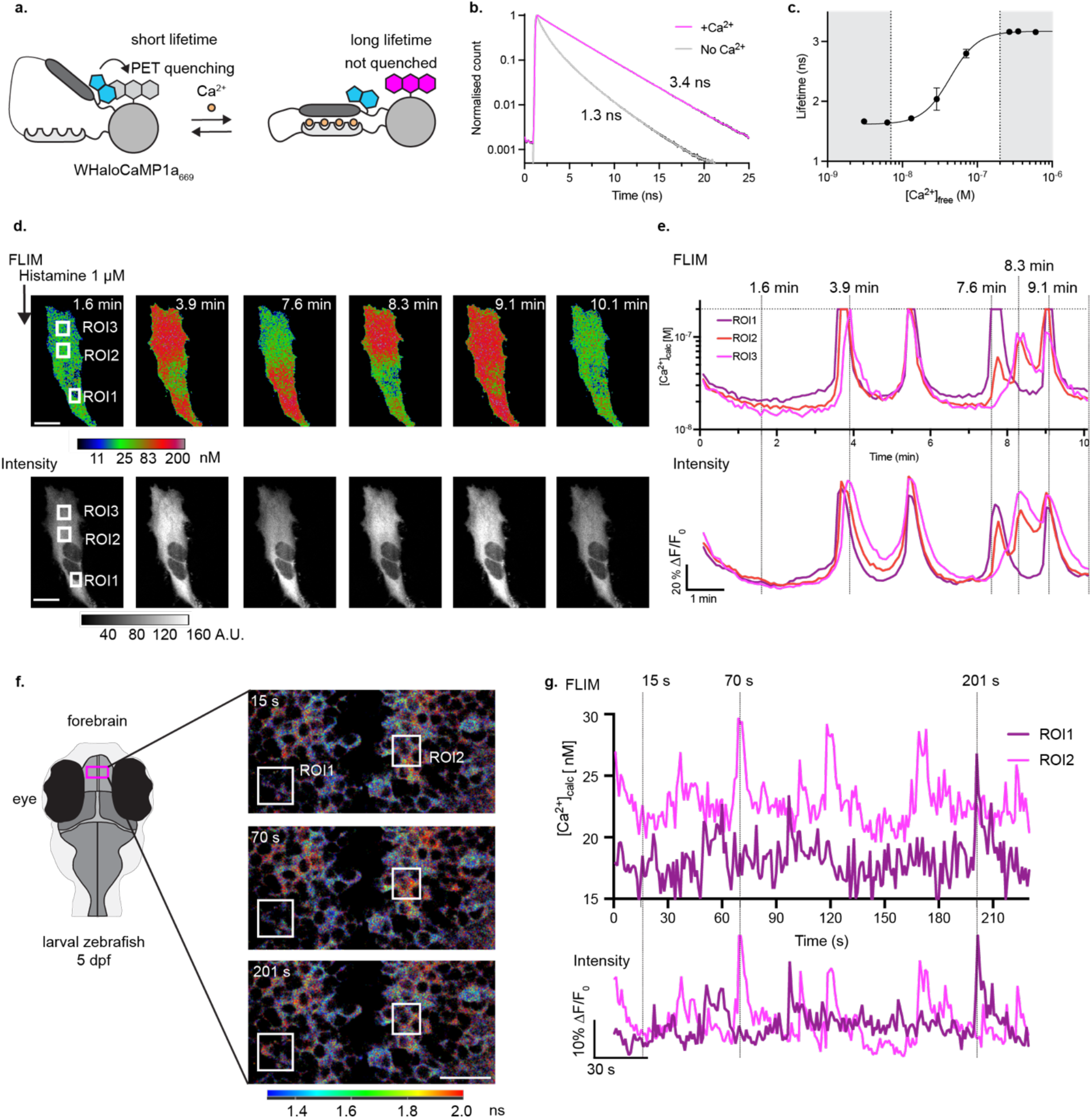
Quantitative Ca^2+^ measurements by florescence lifetime imaging microscopy (FLIM) using WHaloCaMP1a. **a**, Schematic of WHaloCaMP1a bound to a dye-ligand used as a FLIM probe. Tryptophan quenching modulates the fluorescence lifetime. **b**, Normalized fluorescence lifetime of WHaloCaMP1a_669_ in the presence or absence of Ca^2+^, fit to a three-component fluorescence decay. **c**, Calibration curve of averaged fluorescence lifetime of WHaloCaMP1a_669_ vs. [Ca^2+^]. The white box indicates the range where WHaloCaMP1a_669_ can be used to make quantitative measurements of [Ca^2+^]. Performed with purified protein. Mean of three replicates and s.d. plotted. **d**, Pseudocolored concentration (top) and intensity images (bottom) of WHaloCaMP1a_669_ in HeLa cells after histamine addition. Scale bar, 20 µm. Color bar indicates [Ca^2+^], calculated from a calibration curve of fluorescence lifetime. **e**, Quantitative [Ca^2+^] calculated from a FLIM calibration curve (top), and fluorescence traces ΔF/F_0_ calculated from the intensity channel (bottom) in the histamine stimulated HeLa cells in the ROIs highlighted in **d**. Calibrated WHaloCaMP1a_669_ can only be used to measure [Ca^2+^] up to 200 nM, indicated by a dashed horizontal line. Dashed lines vertical indicated time points in the time series at which images in **d** are shown. **f**, FLIM of WHaloCaMP1a_669_ in live zebrafish larvae showing spontaneous neuronal activity in the forebrain. Schematic indicating field of view during imaging (left). Overlaid images of FLIM and intensity using the Leica LASX software, with color bar indicating the fluorescence lifetime. Scale bar, 20 µm. **g**, [Ca^2+^] calculated from a FLIM calibration curve (top), and fluorescence traces ΔF/F_0_ calculated from the intensity channel (bottom) over time for two neurons in the forebrain of zebrafish larvae from the ROIs indicated in **f**. Dashed lines indicated time points of images in **f**. Representative images from three imaging sessions.

WHaloCaMP1a bound to JF_494_, JF_552_, JF_669_ and JF_722_ could all be used as FLIM probes for Ca^2+^ (Supplementary Fig. 31). As WHaloCaMP1a_669_ is brightly fluorescent, we reasoned that it would perform especially well as a FLIM probe in living cells and animals. In purified protein solutions, we observed a 2.1 ns increase in the fluorescence lifetime of WHaloCaMP1a_669_ upon Ca^2+^ binding (τ_apo_ = 1.3 ns vs. τ_sat_ = 3.4 ns) (Fig. 5b). We then generated calibration curves of lifetime vs. [Ca^2+^] both in purified protein and in HeLa cells expressing WHaloCaMP1a_669_ that were permeabilized by digitonin (Fig. 5c and Supplementary Fig. 32). We used these calibration curves to dynamically follow [Ca^2+^] dynamics in HeLa cells stimulated with histamine (Fig. 5d, Supplementary Fig. 33 and Supplementary Movie 5). As the Ca^2+^ affinity of WHaloCaMP1a_669_ is higher than the previously-reported turquoise-emitting FLIM calcium indicator, Tq-Ca-FLITS^36^, we could more accurately determine low [Ca^2+^], down to 7 nM (Fig. 5e). We measured over 100-fold changes in [Ca^2+^] during histamine-induced oscillations in HeLa cells. To determine whether WHaloCaMP allowed *in vivo* FLIM calcium imaging, zebrafish larvae expressing WHaloCaMP1a were labeled with JF_669_-HaloTag ligand and imaged on a commercial confocal FLIM microscope (Fig. 5f and Supplementary Fig. 34). We could follow dynamic changes in absolute intracellular calcium concentration in populations of forebrain neurons at single cell resolution during spontaneous activity at one frame per second (Fig. 5g and Supplementary Movie 6), and calcium events seen by FLIM matched well those observed via fluorescence intensity changes. WHaloCaMP therefore allows the quantitative readout of physiological signaling at cellular resolution across tissue *in vivo*.

## Discussion

Despite their favorable photophysical properties, synthetic dyes have not been fully exploited to build chemigenetic functional indicators with applications in living animals, especially in the near infrared. This has mostly been due to challenges with delivering the dyes in complex tissue and *in vivo*. Here, we started with a near-infrared dye-ligand, JF_669_-HaloTag ligand, that we knew could be delivered to many tissues in mice after vascular injection and built a calcium indicator around it. Since JF_669_-HaloTag ligand did not work with our existing chemigenetic indicators, we utilized a new mechanism to modulate its fluorescence output: PET quenching by a strategically placed tryptophan. This strategy allowed us to record Ca^2+^ transients in the near-infrared channel *in vivo* in several model organisms.

The chemigenetic approach leads to several advantageous properties of WHaloCaMP compared to the biliverdin-binding protein-based indicators. WHaloCaMP1a_669_ is up to 40× brighter than biliverdin-binding indicators. For some applications, the endogenous availability of biliverdin cofactor in cells is advantageous as supplementation of the exogenous cofactor is not required^6, 7^. However, the affinity for biliverdin is often weak in the calcium indicators built from biliverdin-binding proteins, making them dim when used in animals. As opposed to JF_669_-HaloTag ligand, biliverdin does not cross the blood-brain barrier in mice, so additional biliverdin must be generated *in situ* by either delivery of additional biliverdin producing enzymes^37^, or knockouts of biliverdin consuming proteins^38^. A further advantage of WHaloCaMP’s chemigenetic nature over fixed chromophore systems is that the color of the fluorescence emission can be tuned depending on the experimental requirements. Here we show that the same genetically encoded protein sensor, WHaloCaMP1a, can be used with a range of rhodamine dye-ligands with green to near-infrared emission for functional imaging. WHaloCaMP1a thus allows flexibility to match the excitation and emission requirements for specific imaging hardware or for spectrally multiplexed functional imaging.

WHaloCaMP was amenable to known protein engineering strategies to alter Ca^2+^ affinity and binding kinetics, and we envision further protein engineering to optimize maximum fluorescence change. Given the breadth of existing fluorescent protein^39^ and HaloTag/dye-based functional indicators^40^, we anticipate that the tryptophan-quenching chemigenetic indicator approach can be extended to *in vivo* imaging of other physiological signals. Additionally, as we learn more about the bioavailability of dye-ligands in animals, we envision additional chemigenetic approaches where the specificity of genetically encoded tools is combined with the brightness, photostability, and tunability of synthetic dyes.

## Acknowledgements

We thank the Vivarium, Cell Culture, Imaging, Media Prep, Molecular Biology, Virus Production, and the Fly facilities at Janelia Research Campus for assistance. We especially thank D. Walpita, J. Cox, D. Alcor, M. DeSantis, J. Rouchard and S. DiLisio. We thank A. G. Tebo for critical comments on the manuscript and help with collecting X-ray diffraction data, B. Mohar for guidance on dye-ligand choices for use in living animals and S.-H. Sheu for helpful comments on FLIM. We thank the Ahrens Laboratory at Janelia Research Campus for use of animals and useful comments on zebrafish larvae experiments. This work was supported by the Howard Hughes Medical Institute. The contributions by C. M. Hernandez and Y. Chen are supported by the U.S. National Institute of Neurological Disorders and Stroke (R01 NS119821 to Y. C.) and the Mathers Foundation (28323 to Y. C.). M. S. Frei acknowledges funding and resources from the Max Planck Institute for Medical Research, Heidelberg, Germany. Y. Liang was supported by Maryland Stem Cell Research Fund (2022-MSCRFL-5893). The Berkeley Center for Structural Biology is supported in part by the Howard Hughes Medical Institute. The Advanced Light Source is a Department of Energy Office of Science User Facility under contract no. DE-AC02-05CH11231.

## Notes

This article is subject to HHMI’s Open Access to Publications policy. HHMI lab heads have previously granted a nonexclusive CC BY 4.0 license to the public and a sublicensable license to HHMI in their research articles. Pursuant to those licenses, the author-accepted manuscript of this article can be made freely available under a CC BY 4.0 license immediately upon publication.

## Reagent Availability

DNA plasmids encoding WHaloCaMP used in this work are available from Addgene as follows:

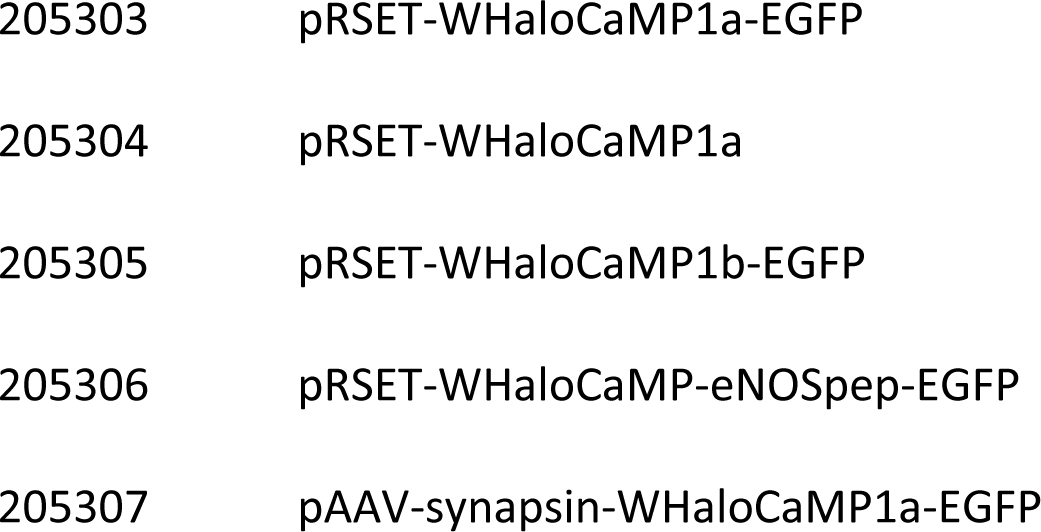
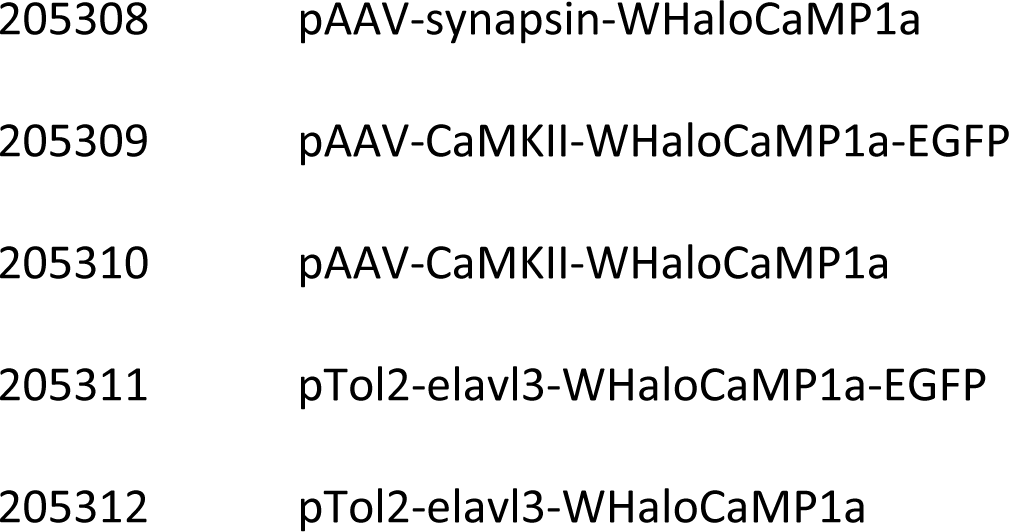

## Contributions

**H. F.** and **E. R. S**. conceived the project and wrote the manuscript with input from all authors. **H. F.** contributed to X-ray crystallography, designed, screened and characterized all WHaloCaMP variants *in vitro* and in neuronal cultures, performed and analyzed functional imaging in zebrafish larvae and in mice, and performed all FLIM experiments with WHaloCaMP. **E. R. S.** contributed to WHaloCaMP design and X-ray crystallography. **Y. S.** performed and analyzed experiments in *Drosophila* and **G. C. T.** oversaw these experiments. **W. C. L.** and **P. J. K.** performed and analyzed data from SiMView light sheet microscopy, and **P. J. K.** oversaw these experiments. **C. M. H.** performed and analyzed experiments in acute brain slices, and **Y. C.** contributed to the analysis and oversaw these experiments. **S. Y.** contributed to optogenetic and electrophysiology experiments. **R. P.** characterized WHaloCaMPs under multiphoton excitation. **G. Q.** performed surgeries for functional imaging of mouse cortex, and **Y. L.** performed and analyzed functional imaging of mouse cortex and oversaw these experiments. **M. S. F.** contributed to WHaloCaMP design. **J. B. G.** and **L. D. L.** synthesized and developed JF-dyes as well as determined their *in vitro* properties and **L. D. L.** oversaw this work. **T. L. H.** and **F. T.** built and developed an assay for functional imaging in mouse cortex. **C. S.** contributed analysis scripts for functional imaging of zebrafish larvae and mice. **A. G. B.** contributed neuronal imaging of WHaloCaMP_722._ **E. R. S.** directed the project.

