## Supplementary Figures for "A modular chemigenetic calcium indicator enables in vivo functional imaging with near-infrared light"

Supplementary figures and tables for:
**A modular chemigenetic calcium indicator enables in vivo functional imaging with near-infrared light**

Helen Farrants, Yichun Shuai, William C. Lemon, Christian Monroy Hernandez, Shang Yang, Ronak Patel, Guanda Qiao, Michelle S. Frei, Jonathan B. Grimm, Timothy L. Hanson, Filip Tomaska, Glenn C. Turner, Carsen Stringer, Philipp J. Keller, Abraham G. Beyene, Yao Chen, Yajie Liang, Luke D. Lavis, *Eric R. Schreiter
*Corresponding author

### Table of Contents

*Supplementary videos* ... 2

*Supplementary Figure 1. Fluorescence emission of $JF_{669}$-HaloTag ligand does not change with addition of $Ca^{2+}$ when used with HaloCaMPs.* ... 3

*Supplementary Figure 2. Engineering strategy, screening and validation approach for WHaloCaMP.* ... 4

*Supplementary Figure 3. Insertion of Calmodulin (CaM) and CaM-binding peptides into HaloTag7 and dye capture rate.* ... 5

*Supplementary Figure 4. Insertion of Calmodulin (CaM) and CaM-binding peptides into HaloTag7 together with rational placement of tryptophan close to the dye binding site.* ... 6

*Supplementary Figure 5. Design of WHaloCaMP1a through three rounds of targeted directed evolution.* ... 7

*Supplementary Figure 6. Kinetics of $Ca^{2+}$ unbinding from WHaloCaMP1a bound to $JF_{669}$-HaloTag ligand.* ... 8

*Supplementary Figure 7. One-photon bleaching during widefield microscopy of $WHaloCaMP1a_{669}$.* ... 9

*Supplementary Fig/Table 8. Comparison of the photophysical properties of $WHaloCaMP1a_{669}$ and biliverdin-binding protein-based calcium indicators.* ... 10

*Supplementary Figure 9. Reducing the $Ca^{2+}$ affinity of WHaloCaMP by rational mutations in the CaM-binding peptide.* ... 11

*Supplementary Figure 10. Peptide swap in WHaloCaMP1a to ENOSP peptide used in the jGCaMP8-series.* ... 12

*Supplementary Figure 11. A second insertion position and tryptophan placement in HaloTag (WHaloCaMP1b).* ... 13

*Supplementary Figure 12. In-gel fluorescence and total protein stain of WHaloCaMP1a labeled with different dyes.* ... 14

*Supplementary Figure 13. Absorption and fluorescence excitation and emission spectra for WHaloCaMP1a* ... 15

*Supplementary Figure 14. Two-photon cross section of WHaloCaMP1a.* ... 16

*Supplementary Figure 15. pH stability of WHaloCaMP1a bound to different dyes.* ... 17

*Supplementary Fig/Table 16. Fluorescence response of WHaloCaMP1a or biliverdin-binding protein-based calcium indicators in a primary neuron culture field stimulation assay with one action potential stimulus.* ... 18

*Supplementary Fig/Table 17. Fluorescence response of WHaloCaMP1a or biliverdin-binding protein-based calcium indicators in a primary neuron culture field stimulation assay with 160 action potentials.* ... 19

*Supplementary Figure 18. Summary of properties of WHaloCaMP1a bound to different dye-ligands in a field stimulation assay in cultured neurons.* ... 20

*Supplementary Fig/Table 19. Decay properties calculated for WHaloCaMP1a bound to different dyes in a field stimulation assay in cultured neurons.* ... 21

*Supplementary Figure 20. $WHaloCaMP1a_{669}$ is spectrally compatible with the channelrhodopsin variant CheRiff.* ... 22

*Supplementary Figure 21. Multiplexed imaging in acute brain slices with $WHaloCaMP1a_{669}$ and FLIM-AKAR.* ... 23

### Supplementary videos

1. Volumetric SiMView light sheet imaging of WHaloCaMP1a$_{669}$ in zebrafish larva
2. Three color functional multiplexed imaging of WHaloCaMP1a$_{669}$, jRGECO1a and iGlucoSnFR in zebrafish larva
3. Dual color imaging of WHaloCaMP1a$_{669}$ in neurons and jRGECO1b in astrocytes in zebrafish larva
4. Dual color imaging of WHaloCaMP1a$_{669}$ in neurons and jRGECO1b in astrocytes during a seizure-like state induced by 4-aminopyridine
5. FLIM of WHaloCaMP1a$_{669}$ expressed in HeLa cells stimulated by histamine
6. FLIM of WHaloCaMP1a$_{669}$ in the forebrain of zebrafish larva

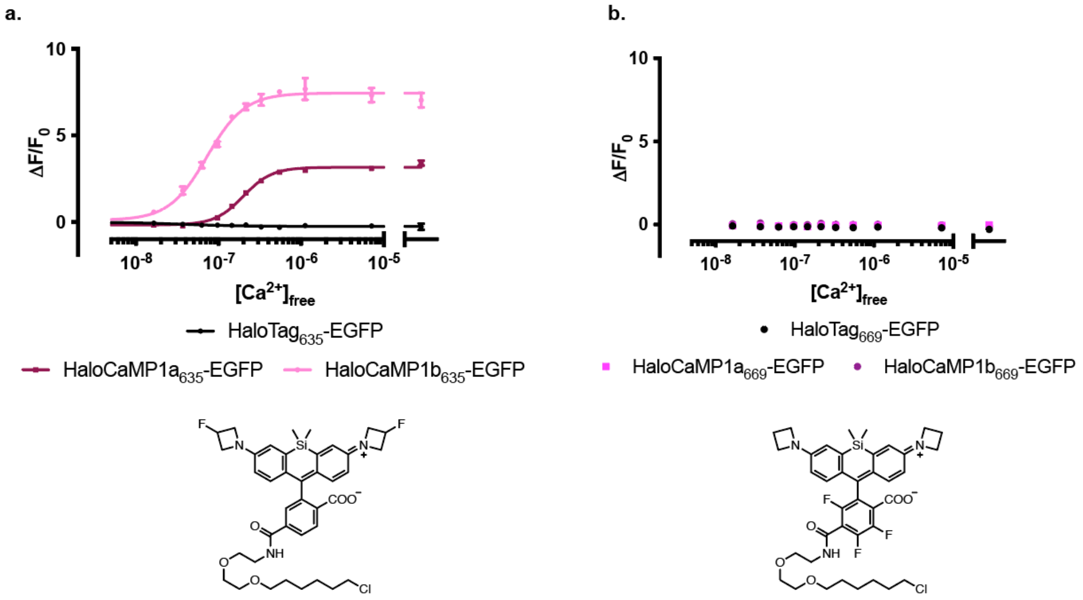

Supplementary Figure 1. Fluorescence emission of $JF_{669}$-HaloTag ligand does not change with addition of $Ca^{2+}$ when used with HaloCaMPs.

**a**, $Ca^{2+}$ titrations with HaloTag7[1]-EGFP, HaloCaMP1a and HaloCaMP1b[2] labeled with $JF_{635}$-HaloTag ligand. **b**, $Ca^{2+}$ titrations of same protein variants as in (**a.**) but labeled with $JF_{669}$-HaloTag ligand.

HaloCaMP1$a_{635}$ and HaloCaMP1$b_{635}$ are both calcium indicators, while HaloCaMP1$a_{669}$ and HaloCaMP1$b_{669}$ are not.

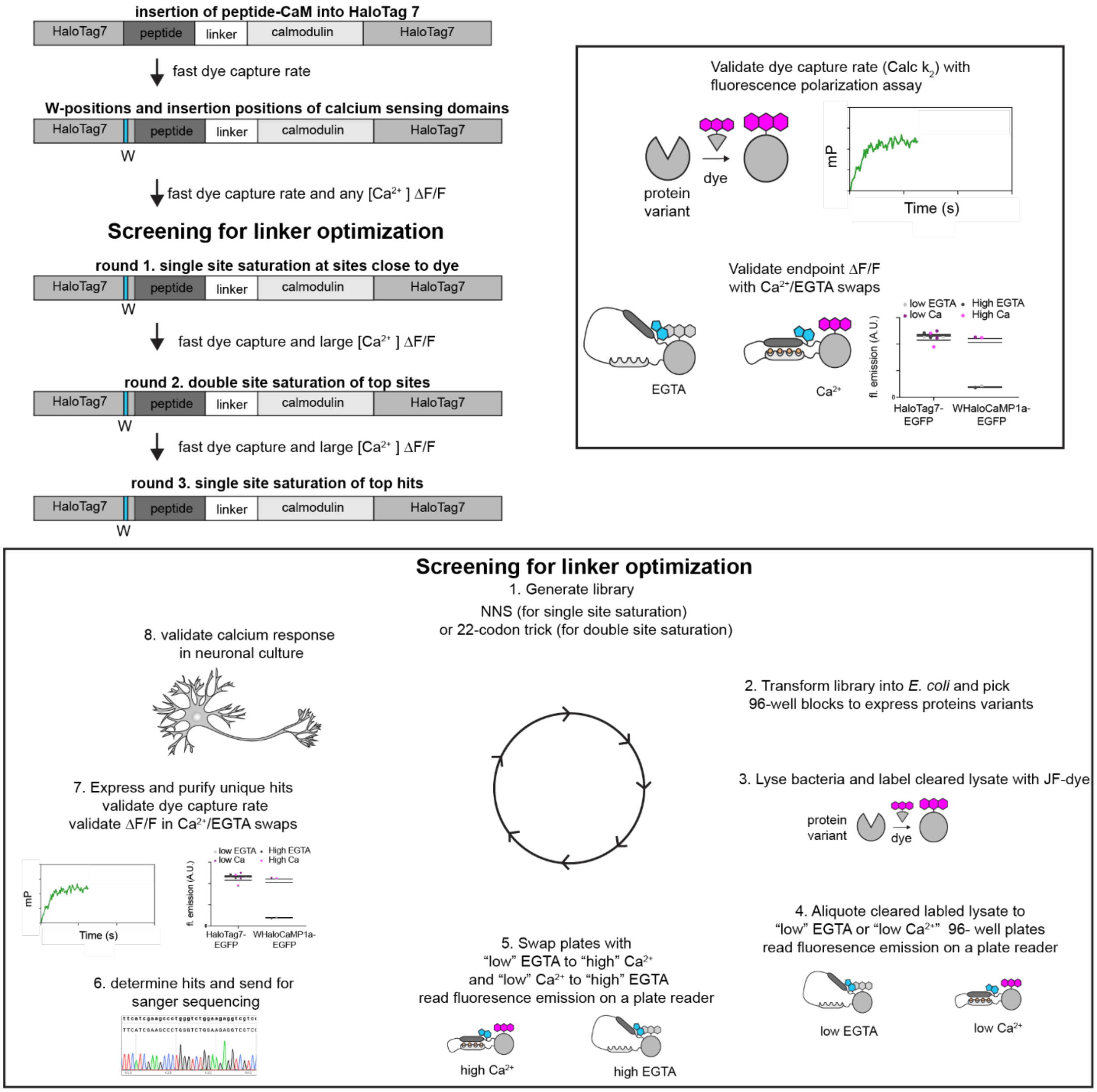

### Supplementary Figure 2. Engineering strategy, screening and validation approach for WHaloCaMP.

Engineering of WHaloCaMPs were performed in several consecutive steps. First, insertion sites of Calmodulin (CaM) and a CaM-binding peptide were explored. Insertion sites with high dye capture rate were moved forward to explore rational tryptophan mutations together with insertions. The variants with fastest dye capture rate and any fluorescence modulation upon calcium addition were moved forward to targeted directed evolution through single or double site saturation mutagenesis. After each round, $Ca^{2+}$ indicator performance was validated in cultured neurons in a field stimulation assay.

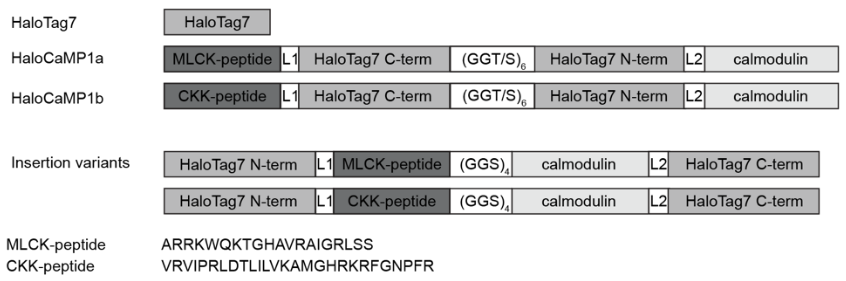

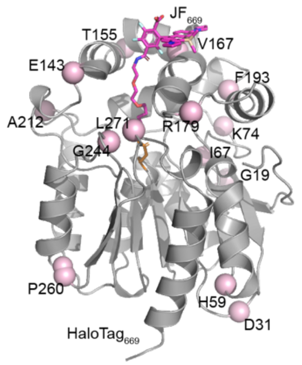

| | EGTA Calc $k_2$ ($M^{-1}$ $s^{-1}$) | $Ca^{2+}$ Calc $k_2$ ($M^{-1}$ $s^{-1}$) |
|---|---|---|
| **HaloTag7** | Over $10^6$ | Over $10^6$ |
| **HaloCaMP1a** | $1 \times 10^5 \pm 1 \times 10^4$ | Over $10^6$ |
| **HaloCaMP1b** | $1 \times 10^5 \pm 7 \times 10^3$ | $8 \times 10^4 \pm 9 \times 10^3$ |

| Insertion at | L1 | Peptide-$(GGS)_4$-CaM /CaM-$(GGS)_4$-peptide | L2 | Calc $k_2$ ($M^{-1}$ $s^{-1}$) EGTA | Calc $k_2$ ($M^{-1}$ $s^{-1}$) $Ca^{2+}$ |
|---|---|---|---|---|---|
| **G19** | - | MLCK-$(GGS)_4$-CaM | - | Over $10^6$ | Over $10^6$ |
| **D31** | - | MLCK-$(GGS)_4$-CaM | - | Over $10^6$ | Over $10^6$ |
| **H59** | - | MLCK-$(GGS)_4$-CaM | - | Over $10^6$ | Not measured |
| **I67** | - | MLCK-$(GGS)_4$-CaM | - | $6 \times 10^3 \pm 3 \times 10^3$ | Below $10^3$ |
| **K74** | - | MLCK-$(GGS)_4$-CaM | - | Over $10^6$ | Over $10^6$ |
| **E143** | GGS | MLCK-$(GGS)_4$-CaM | GGS | $1 \times 10^5 \pm 1 \times 10^4$ | $1 \times 10^5 \pm 1 \times 10^4$ |
| | - | MLCK-$(GGS)_4$-CaM | - | $1 \times 10^5 \pm 2 \times 10^4$ | $1 \times 10^5 \pm 2 \times 10^4$ |
| **T155** | Delta T155 | MLCK-$(GGS)_4$-CaM | Delta D156 | Over $10^6$ | Over $10^6$ |
| | Delta T154 Delta T155 | MLCK-$(GGS)_4$-CaM | Delta D156 Delta V157 | Over $10^6$ | $9 \times 10^4 \pm 9 \times 10^3$ |
| | Delta R153 Delta T154 Delta T155 | MLCK-$(GGS)_4$-CaM | Delta D156 Delta V157 Delta G158 | Below $10^3$ | Below $10^3$ |
| | GGS | CKK-$(GGS)_4$-CaM | GGS | Over $10^6$ | Over $10^6$ |
| | - | CKK-$(GGS)_4$-CaM | - | Over $10^6$ | $4 \times 10^5 \pm 1 \times 10^5$ |
| | Delta T155 | CKK-$(GGS)_4$-CaM | Delta D156 | Over $10^6$ | $3 \times 10^5 \pm 1 \times 10^4$ |
| | Delta T154 Delta T155 | CKK-$(GGS)_4$-CaM | Delta D156 Delta V157 | $1 \times 10^4 \pm 8 \times 10^3$ | $7 \times 10^4 \pm 4 \times 10^3$ |
| **V167** | - | MLCK-$(GGS)_4$-CaM | - | $5 \times 10^3$ | Below $10^3$ |
| **R179** | - | MLCK-$(GGS)_4$-CaM | - | Over $10^6$ | $8 \times 10^4 \pm 1 \times 10^4$ |
| **F193** | - | MLCK-$(GGS)_4$-CaM | - | Over $10^6$ | Over $10^6$ |
| **A212** | - | MLCK-$(GGS)_4$-CaM | - | $3 \times 10^5$ | $5 \times 10^5$ |
| **G244** | - | MLCK-$(GGS)_4$-CaM | - | Below $10^3$ | Below $10^3$ |
| **P260** | - | MLCK-$(GGS)_4$-CaM | - | $3 \times 10^5 \pm 1 \times 10^5$ | $3 \times 10^5 \pm 3 \times 10^4$ |
| **L271** | - | MLCK-$(GGS)_4$-CaM | - | $6 \times 10^3$ | Below $10^3$ |

Supplementary Figure 3. Insertion of Calmodulin (CaM) and CaM-binding peptides into HaloTag7 and dye capture rate.

1D schematic of the topology of the insertion variants of CaM and CaM-binding peptide into HaloTag7 in comparison to HaloCaMP. The crystal structure of HaloTag7 bound to $JF_{669}$-HaloTag ligand is shown, highlighting the insertion positions. Bottom two tables report the measured dye capture rate with $JF_{549}$-HaloTag ligand and the indicated protein. The insertion position (R179) which led to WHaloCaMP1a is highlighted in magenta.

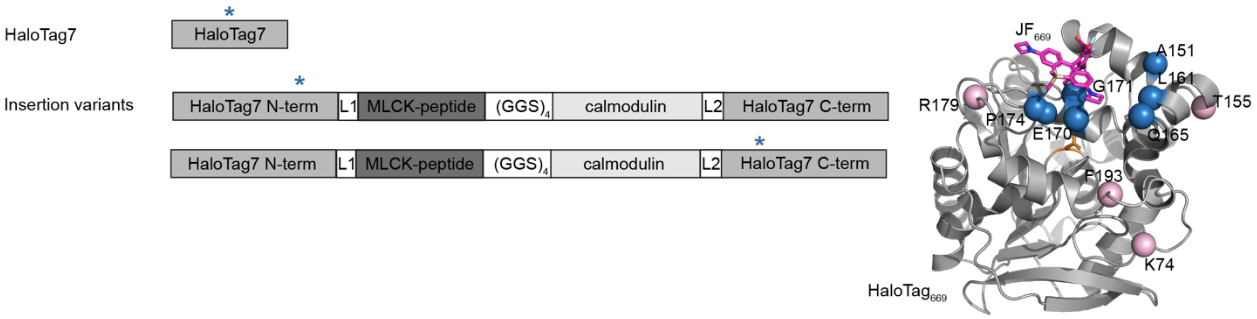

| Protein | W mutation | Calc $k_2$ ($M^{-1}$ $s^{-1}$) EGTA | ΔF/F |
|---|---|---|---|
| **HaloTag7** | G171W | Over $10^6$ | $0.0 \pm 0.1$ |
| **HaloTag7** | A151W | Over $10^6$ | $0.0 \pm 0.1$ |

| Insertion at | Insertion of | W mutation | Calc $k_2$ ($M^{-1}$ $s^{-1}$) EGTA | ΔF/F |
|---|---|---|---|---|
| **K74** | MLCK -(GGS)$_4$- CaM | G171W | Over $10^6$ | $0.0 \pm 0.1$ |
| **T155** | MLCK -(GGS)$_4$- CaM | G171W | Over $10^6$ | $0.0 \pm 0.0$ |
| | Delta T155 MLCK -(GGS)$_4$-CaM Delta D156 | A151W | $4 \times 10^5$ | $0.2 \pm 0.1$ |
| | Delta T154 Delta T155 MLCK -(GGS)$_4$-CaM Delta D156 Delta V157 | A151W | $2 \times 10^5$ | $-0.3 \pm 0.1$ |
| | Delta T155 MLCK -(GGS)$_4$- CaM Delta D156 | L161W | $6 \times 10^4 \pm 7 \times 10^3$ | $0.2 \pm 0.1$ |
| | Delta T155 MLCK -(GGS)$_4$- CaM Delta D156 | Q165W | $3 \times 10^5 \pm 1 \times 10^4$ | $0.2 \pm 0.1$ |
| | Delta T155 MLCK -(GGS)$_4$- CaM Delta D156 | E170W | $4 \times 10^3$ | $-0.1 \pm 0.2$ |
| **R179** | MLCK - (GGS)$_4$-CaM | P174W | $2 \times 10^4 \pm 5 \times 10^3$ | $0.2 \pm 0.1$ |
| | MLCK - (GGS)$_4$-CaM | G171W | Over $10^6$ | $0.2 \pm 0.1$ |
| **F193** | MLCK - (GGS)$_4$-CaM | G171W | Over $10^6$ | $0.0 \pm 0.2$ |

Supplementary Figure 4. Insertion of Calmodulin (CaM) and CaM-binding peptides into HaloTag7 together with rational placement of tryptophan close to the dye binding site.

1D schematic of the topology of insertion variants of CaM and CaM-binding peptide into HaloTag7 (top left). Tryptophan positions are annotated with a blue star. The crystal structure of HaloTag7 bound to $JF_{669}$-HaloTag ligand is shown, highlighting the insertion positions and tryptophan placements (top right). Bottom tables show the measured dye capture rate with $JF_{549}$-HaloTag ligand and the indicated protein, and any change in fluorescence emission ΔF/F with $JF_{669}$-HaloTag ligand. The insertion position (R179), and the tryptophan placement (G171W) which led to WHaloCaMP1a is highlighted in magenta.

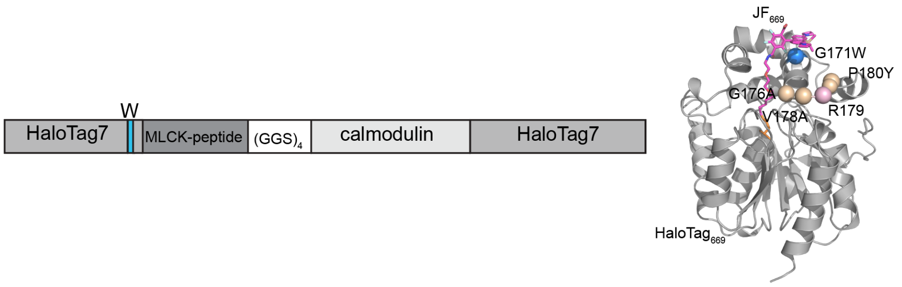

Round 1: single site saturations on top of insertion at site R179 with G171W

**V167X, I169X, E170X, L173X, P174X, M175X, G176X, V178X, R179X, P180X, L181X, T182X**

| Mutation | Calc $k_2$ ($M^{-1}$ $s^{-1}$) EGTA | Calc $k_2$ ($M^{-1}$ $s^{-1}$) $Ca^{2+}$ | ΔF/F |
|---|---|---|---|
| **G176V** | | | $1.2 \pm 0.1$ |
| **G176C** | | | $0.8 \pm 0.2$ |
| **V178R** | $4 \times 10^5 \pm 4 \times 10^4$ | $3 \times 10^4 \pm 1 \times 10^4$ | $2.1 \pm 0.2$ |
| **P180Y** | | | $1.4 \pm 0.3$ |
| **P180V** | | | $1.1 \pm 0.2$ |
| **P180I** | | | $1.3 \pm 0.2$ |
| **P180W** | Over $10^6$ | $3 \times 10^4 \pm 9 \times 10^3$ | $1.8 \pm 0.2$ |
| **P180R** | | | $1.5 \pm 0.2$ |

Round 2: double site saturation of top sites of round 1

**V178X, P180X**

| Mutation | Mutation | Calc $k_2$ ($M^{-1}$ $s^{-1}$) EGTA | Calc $k_2$ ($M^{-1}$ $s^{-1}$) $Ca^{2+}$ | ΔF/F |
|---|---|---|---|---|
| **V178A** | **P180Y** | Over $10^6$ | $8 \times 10^4 \pm 3 \times 10^4$ | $6.3 \pm 0.0$ |
| **V178A** | **P180V** | Over $10^6$ | $9 \times 10^4 \pm 3 \times 10^4$ | $5.0 \pm 0.1$ |
| **V178C** | **P180Y** | Over $10^6$ | $1 \times 10^5 \pm 3 \times 10^4$ | $5.1 \pm 0.0$ |
| **V178C** | **P180G** | Over $10^6$ | $1 \times 10^5 \pm 4 \times 10^4$ | $4.5 \pm 0.5$ |
| **V178C** | **P180R** | Over $10^6$ | $3 \times 10^5 \pm 9 \times 10^4$ | $4.6 \pm 0.5$ |

Round 3: single site saturation on top of hit from round 2

**T148X, L161X, Q165X, I169X, E170, T172X, P174X, G176X,**

| Mutation | Mutation | Mutation | Calc $k_2$ ($M^{-1}$ $s^{-1}$) EGTA | Calc $k_2$ ($M^{-1}$ $s^{-1}$) $Ca^{2+}$ | ΔF/F |
|---|---|---|---|---|---|
| **V178A** | **P180Y** | **G176A** | Over $10^6$ | $2 \times 10^5 \pm 7 \times 10^4$ | $5.6 \pm 0.8$ |
| **V178A** | **P180Y** | **Q165R** | $2 \times 10^5 \pm 1 \times 10^5$ | $8 \times 10^3 \pm 3 \times 10^3$ | $5.9 \pm 0.6$ |

Supplementary Figure 5. Design of WHaloCaMP1a through three rounds of targeted directed evolution.

1D schematic of the topology (top left) of the insertion variants of CaM and CaM-binding peptide into HaloTag7 with tryptophan position and highlighted residues on the crystal structure of $HaloTag_{669}$ (top right). Tables report top hits from sequential mutagenic library screens. First, sites close to the dye binding site were chosen for single site saturation. Double site saturation was then performed on the top two sites from the first round of screening. A third round of single site saturation on top of the be best performing variant was then performed with sites close to the dye binding site. After each round, dye capture rates of top hits were measured by fluorescence polarization using $JF_{549}$-HaloTag ligand and a calcium endpoint assay measured to validate fluorescence modulation by calcium using $JF_{669}$-HaloTag ligand. WHaloCaMP1a is highlighted in magenta.

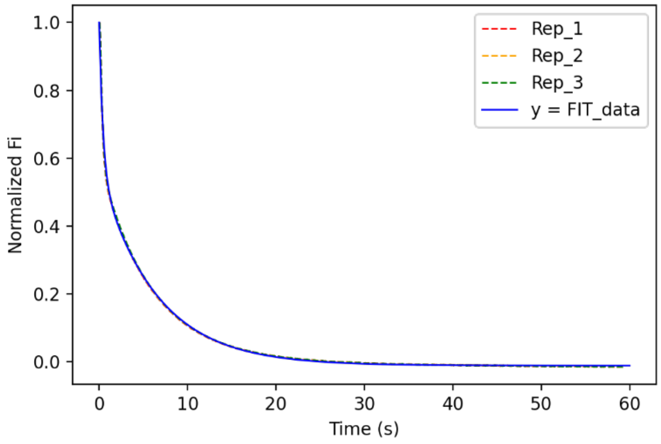

| | $WHaloCaMP1_{a669}$ |
|---|---|
| Model | $y = a_0 + a_1(1 - e^{-b_1 \times t}) + a_2(1 - e^{-b_2 \times t})$ |
| fit | $y = 1 + (-0.57)(1 - e^{-0.16 \times t}) + (-0.43)(1 - e^{-2.3 \times t})$<br>$a_0$ was constrained to 1.0. |
| $k_{off}$ ($s^{-1}$) | $Dissociative_1$: 0.16<br>$Dissociative_2$: 2.3 |

Supplementary Figure 6. Kinetics of $Ca^{2+}$ unbinding from WHaloCaMP1a bound to $JF_{669}$-HaloTag ligand.

A stopped flow instrument was used to follow the decrease in fluorescence emission from recombinant calcium saturated $WHaloCaMP1a_{669}$ following rapid mixing with excess calcium chelator (EGTA, 10 mM). Fluorescence decay was fit to a two-phase exponential model. Three technical replicates (Rep1, Rep 2, Rep 3) was performed, normalized to the initial fluorescence intensity at time 0.

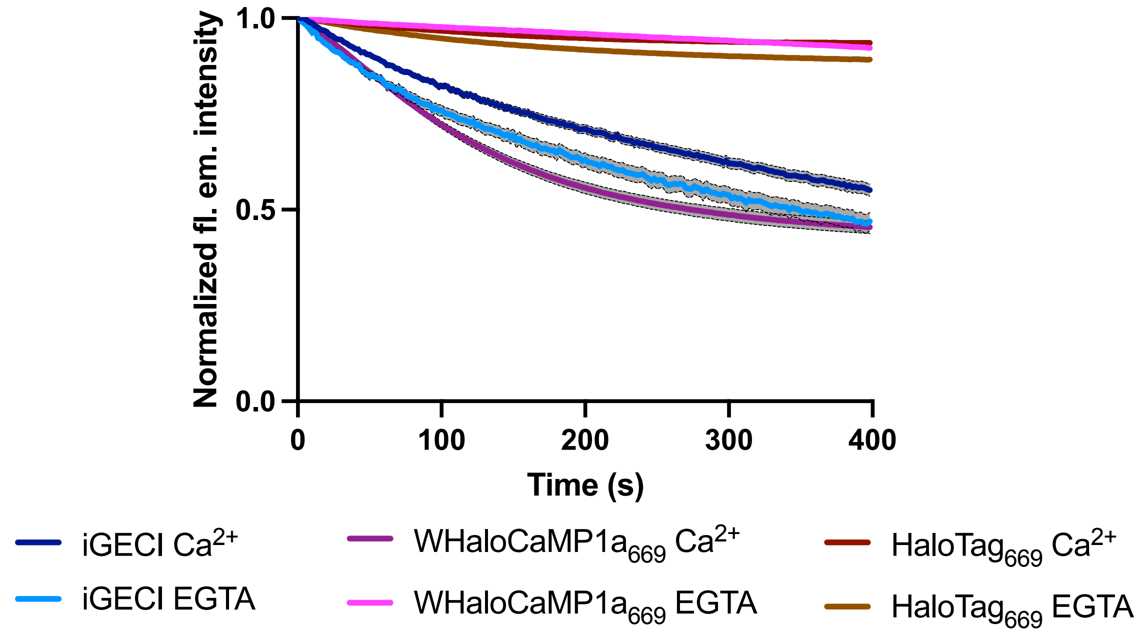

| Protein* | Halftime (s) | 95% confidence interval halftime (s) |
|---|---|---|
| **iGECI, $Ca^{2+}$** | 204 | 190 to221 |
| **iGECI, EGTA** | 160 | 148 to 176 |
| **WhaloCaMP1a$_{669}$, $Ca^{2+}$** | 98 | 95 to 102 |

*one-photon bleaching halftimes of HaloTag$_{669}$ in $Ca^{2+}$, HaloTag$_{669}$ in EGTA, and WHaloCaMP1a$_{669}$ in EGTA could not be fit under these experimental conditions. One-photon bleaching halftimes of these proteins are longer than 400 s under these illumination conditions.

Supplementary Figure 7. One-photon bleaching during widefield microscopy of WHaloCaMP1a$_{669}$.

Aqueous droplets of purified protein in octanol were used to determine bleaching rates on an inverted widefield microscope. The droplets were continuously illuminated at 23 mW/mm$^2$ for 400 seconds, imaging at 0.5 Hz. Approximately 20 droplets were imaged in each bleaching trial and three bleaching trials were pooled to make the bleaching curve. Each fluorescence trace was normalized to the fluorescence emission in the first frame. The bleaching curves were fit with a one phase decay in GraphPad Prism software.

| | Ex/Em$_{apo}$ (nm) | Ex/Em$_{sat}$ (nm) | Dynamic range ($F_{max}/F_{min}$) | $K_d$ (nM) | Hill coeff. | $\varepsilon_{apo}$ (x1000) ($M^{-1}$ $cm^{-1}$) | $\varepsilon_{sat}$ (x1000) ($M^{-1}$ $cm^{-1}$) | $\Phi_{apo}$ | $\Phi_{sat}$ | Brightness$_{apo}$ ($mM^{-1}$ $cm^{-1}$) | Brightness$_{sat}$ ($mM^{-1}$ $cm^{-1}$) |
|---|---|---|---|---|---|---|---|---|---|---|---|
| WHaloCaMP1a$_{669}$ | 677/688 | 679/690 | 7 | 37 ± 2 | 2.5 ± 0.3 | 127 | 174 | 0.08 | 0.50 | 10.1 | 87.0 |
| NIR-GECO1* | 678/704 | 678/704 | ($\Delta F/F_{min}$) -9 | 885 | 1.0 | 69 | 20 | 0.06 | 0.02 | 4.1 | 0.4 |
| NIR-GECO2* | 678/704 | 678/704 | ($\Delta F/F_{min}$) -15 | 331 | 0.9 | 67 | 18 | 0.06 | 0.01 | 4.0 | 0.3 |
| NIR-GECO2G* | 678/704 | 678/704 | ($\Delta F/F_{min}$) -9 | 480 | 0.8 | 74 | 21 | 0.06 | 0.02 | 4.5 | 0.4 |
| iBB-GECO1* | 648/668 | 646/653 | ($\Delta F/F_{min}$) -13 | 105 | 1.9 | 52 | 16 | 0.11 | 0.03 | 6.2 | 0.5 |
| GAF-CaMP2-sfGFP[o] | 630/676 | 642/674 | ($\Delta F/F_{min}$) 0.9 | 289 | 1.53[oo] | 16 | 28 | 0.03 | 0.06 | 0.5 | 1.6 |
| GAF-CaMP3-sfGFP[oo] | 636/674 | 648/676 | ($\Delta F/F_{min}$) 2 | 433 | 1.36 | 14 | 27 | 0.05 | 0.08 | 0.7 | 2.1 |
| iGECI[#] | donor 640/670 acceptor 702/720 | donor 640/670 acceptor 702/720 | 6 | 15 and 890 | 2.5 and 0.9 | - | - | - | - | 12.3 | 2.0 |
| iGECInano[##] | donor 645/670 acceptor 702/720 | donor 640/670 acceptor 702/720 | 4 | 530 | 1.53 | - | - | - | - | - | - |

Supplementary Fig/Table 8. Comparison of the photophysical properties of WHaloCaMP1a$_{669}$ and biliverdin-binding protein-based calcium indicators.

*from (ref)[3]

[o]from (ref)[4]

[oo]from (ref)[5]

[#]from (ref)[6]

[##]from (ref)[7]

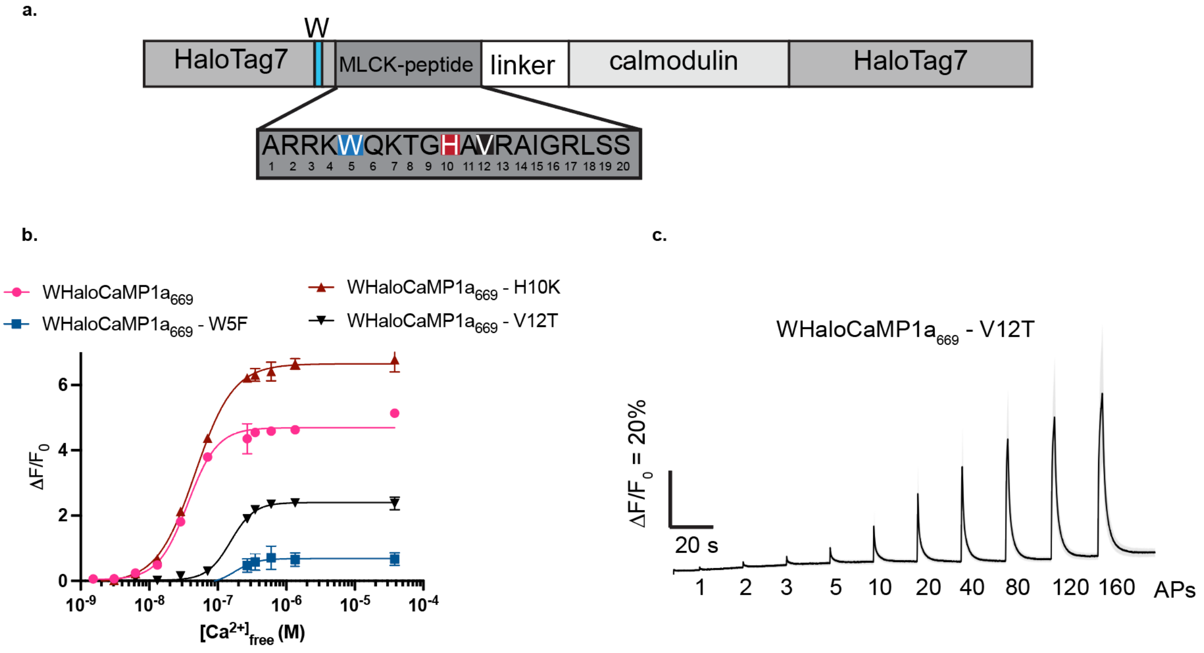

| Peptide mutation | $EC_{50}$ (nM) | Hill | Max $\Delta F/F_0$ |
|---|---|---|---|
| **MLCK peptide** | 37 ± 2 | 2.5 ± 0.3 | 4.7 ± 0.1 |
| **W5F** | 179 ± 55 | 2.7 ± 1.5 | 0.8 ± 0.08 |
| **H10K** | 46 ± 2 | 1.5 ± 0.1 | 6.7 ± 0.1 |
| **V12T** | 154 ± 88 | 2.5 ± 0.2 | 2.3 ± 0.5 |

Supplementary Figure 9. Reducing the $Ca^{2+}$ affinity of WHaloCaMP by rational mutations in the CaM-binding peptide.

**a**, Previously described mutations in the CaM-binding peptide[8] were mapped onto WHaloCaMP1a. **b**, $Ca^{2+}$ titrations of MLCK peptide variants (top) and values extracted from curve fits (bottom table). **c**, Fluorescence trace (black) from WHaloCaMP1a$_{669}$-V12T in primary neuron culture with field stimulation. Grey trace is s.e.m. $F_0$ was calculated 1 s before the first field stimulus. WHaloCaMP1a$_{669}$-V12T showed faster fluorescence decay in neurons compared to WHaloCaMP1a, but lower $\Delta F/F_0$ response at single AP, therefore WHaloCaMP1a$_{669}$ was used for *in vivo* demonstrations.

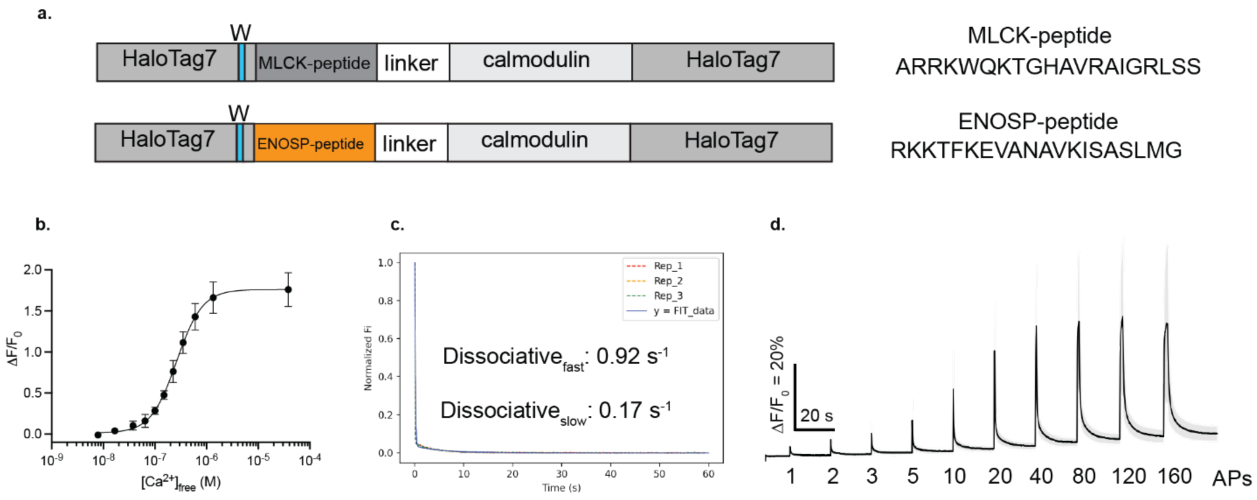

**Round 1:**Diversify 178X and P180X →hit V178I and P180T

**Round 2:**Diversify T148X, L161X, Q165X, I169X, E170X, T172X, P174X, G176X

| variant | Calc $k_2$ ($M^{-1}$ $s^{-1}$) EGTA | Calc $k_2$ ($M^{-1}$ $s^{-1}$) $Ca^{2+}$ | Max $\Delta F/F_0$ (endpoint) | $EC_{50}$ (nM) | Hill |
|---|---|---|---|---|---|
| ENOSP<br>G171W - W quench | Over $10^6$ | $1 \times 10^5 \pm 3 \times 10^4$ | $1.5 \pm 0.1$ | - | - |
| ENSOP - peptide<br>G171W - W quench<br>V178I, P180T, G176A | Over $10^6$ | $4 \times 10^5 \pm 2 \times 10^5$ | $4.6 \pm 0.4$ | 263 ± 17 | 1.8 ± 0.2 |

Supplementary Figure 10. Peptide swap in WHaloCaMP1a to ENOSP peptide used in the jGCaMP8-series.

**a**, The MLCK peptide used in WHaloCaMP1a (top) was replaced with the ENOSP-peptide used in jGCaMP8 series[9] (bottom). Two rounds of directed evolution were performed on the new scaffold: the first round was a double site saturation of sites important in the evolution of WHaloCaMP1a; the second was single site saturation on top of the top hit from the first round, which gave a variant (V178I, P180T, G176A) with fast dye capture rate, lower $Ca^{2+}$ affinity (**b.**), and fast $Ca^{2+}$ kinetics measured by stopped flow (**c.**). **d**, A representative fluorescence trace from a field stimulation (black) from one run of the field stimulation assay with ~40 neuron ROIs. Grey trace is standard error of the mean (s.e.m.) of the fluorescence trace. $F_0$ was calculated 1 s before the first field stimulus. Table reports fits from experiments in panels **b**. Even though the ENOSP variant had faster $Ca^{2+}$ kinetics in neurons, it had lower $\Delta F/F_0$ response at single AP, therefore WHaloCaMP1a$_{669}$ was used for *in vivo* demonstrations.

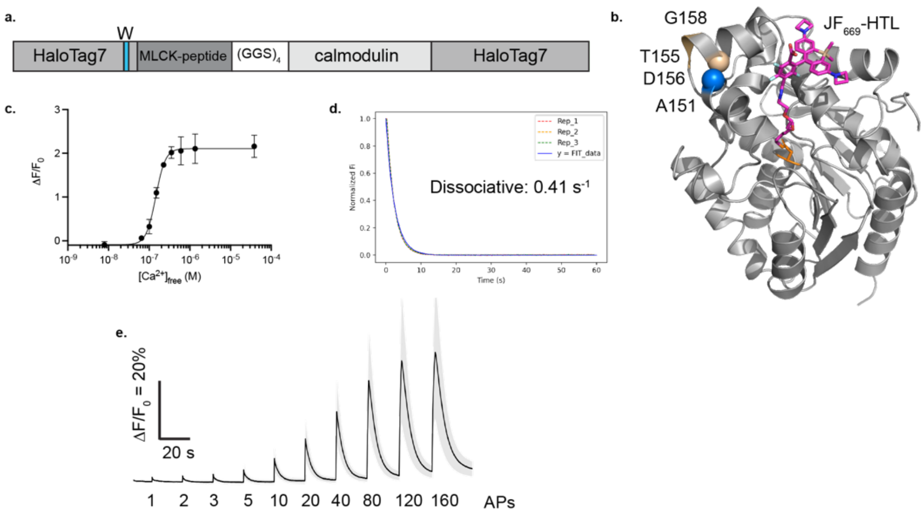

**Round 1**: single site saturations on top of insertion at site Delta T155 MLCK -$(GGS)_4$-CaM Delta D156 with A151W: G158X, K160X, L161X, R153X, T154X, V157X, Q165X.

| Mutation | Calc $k_2$ ($M^{-1}$ $s^{-1}$) EGTA | Calc $k_2$ ($M^{-1}$ $s^{-1}$) $Ca^{2+}$ | ΔF/F | $EC_{50}$ (nM) | Hill |
|---|---|---|---|---|---|
| **V157K** | Over $10^6$ | $1 \times 10^5 \pm 1 \times 10^4$ | $0.5 \pm 0.2$ | | |
| **G158D (WHaloCaMP1b)** | $9 \times 10^4 \pm 7 \times 10^3$ | $1 \times 10^4 \pm 3 \times 10^3$ | $2.5 \pm 0.4$ | 145 ± 6 | 3.7 ± 0.5 |

### Supplementary Figure 11. A second insertion position and tryptophan placement in HaloTag (WHaloCaMP1b).

**a**, The MLCK CaM-binding peptide and CaM were inserted at HaloTag T154, deleting T155 and D156. One round of single site saturation mutagenesis produced hits with improved calcium response (G158D). Further rounds of directed evolution did not give better variants. **b**, Crystal structure of $HaloTag7_{669}$ highlighting relevant positions. **c**, $\Delta F/F_0$ vs. $[Ca^{2+}]$ of $WHaloCaMP1b_{669}$. **d**, Stopped-flow $Ca^{2+}$ kinetics of unbinding for $WHaloCaMP1b_{669}$. **e**, Representative trace (black) from one run of the field stimulation assay with ~40 neuron ROIs. Grey trace is standard error of the mean (s.e.m.) of the fluorescence trace. $F_0$ was calculated 1 s before the first field stimulus. The table reports fits from experiments in panels **c**.

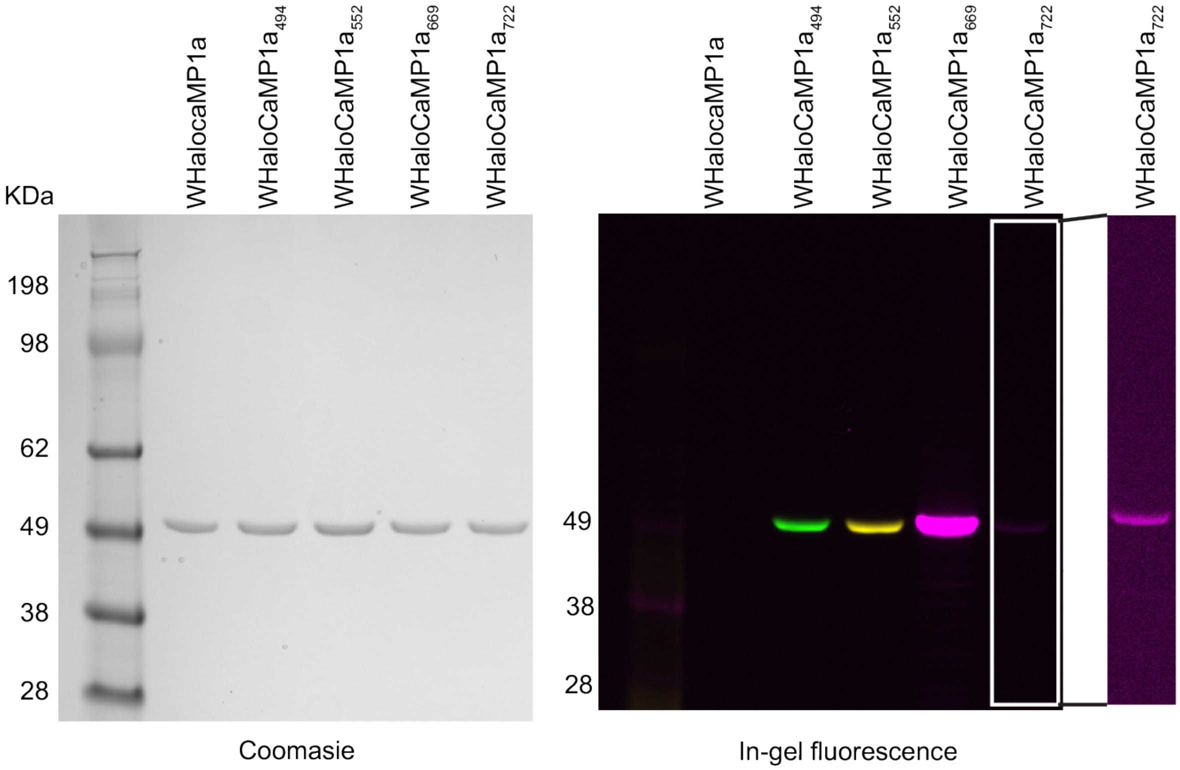

Supplementary Figure 12. In-gel fluorescence and total protein stain of WHaloCaMP1a labeled with different dyes.

Denaturing sodium dodecyl-sulfate polyacrylamide gel electrophoresis (SDS-PAGE) was used to separate purified WHaloCaMP1a labeled with different dye-ligands and LEDs with corresponding filters on a gel imager were used to image the in-gel fluorescence. $WHaloCaMP1a_{722}$ could be imaged with far-red filters if the brightness and contrast was increased in comparison to $WHaloCaMP1a_{669}$, as seen in the white box to the far right.

**a.**

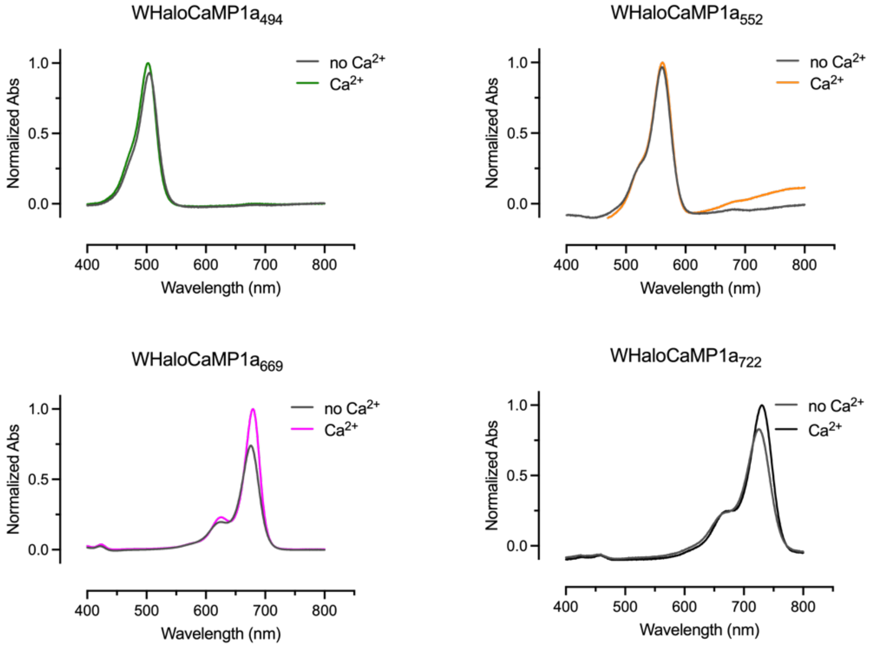

**b.**

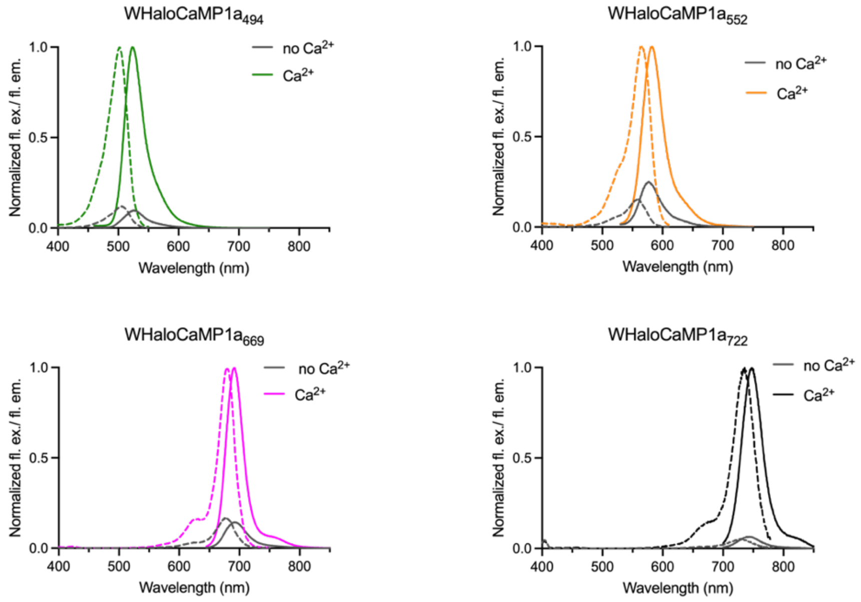

Supplementary Figure 13. Absorption and fluorescence excitation and emission spectra for WHaloCaMP1a .

WHaloCaMP1a bound to $JF_{494}$-HaloTag ligand, $JF_{552}$-HaloTag ligand, $JF_{669}$-HaloTag ligand and $JF_{722}$-HaloTag ligand. Absorption (a). and fluorescence excitation and emission spectra (b.). Spectra were normalized to max absorbance or flourescence exitation or emission in the $Ca^{2+}$ state.

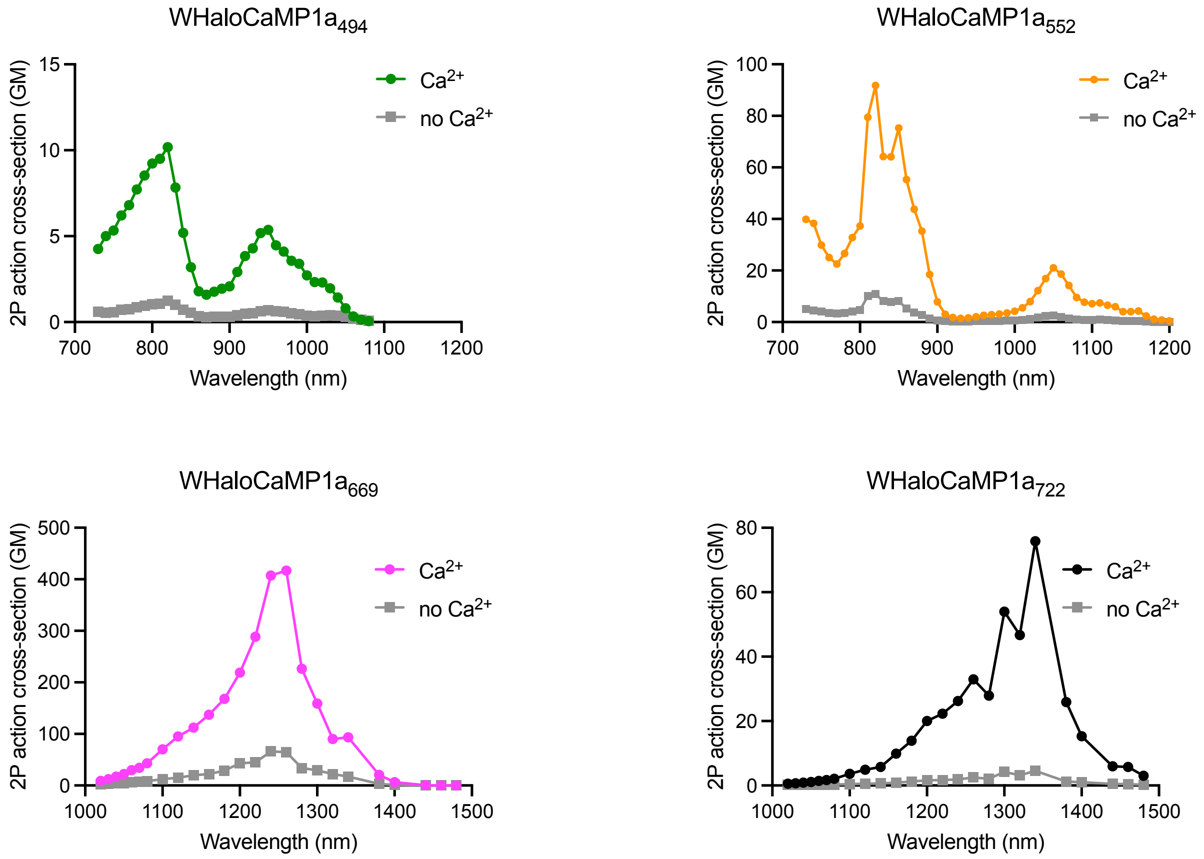

Supplementary Figure 14. Two-photon cross section of WHaloCaMP1a.

WHaloCaMP1a bound to JF$_{494}$-HaloTag ligand, JF$_{552}$-HaloTag ligand, JF$_{669}$-HaloTag ligand and JF$_{722}$-HaloTag ligand. 2P action cross section spectra were collected in purified protein prelabeled with dyes. $Ca^{2+}$ is 39 µM free $Ca^{2+}$ and no $Ca^{2+}$ is 10 mM EGTA.

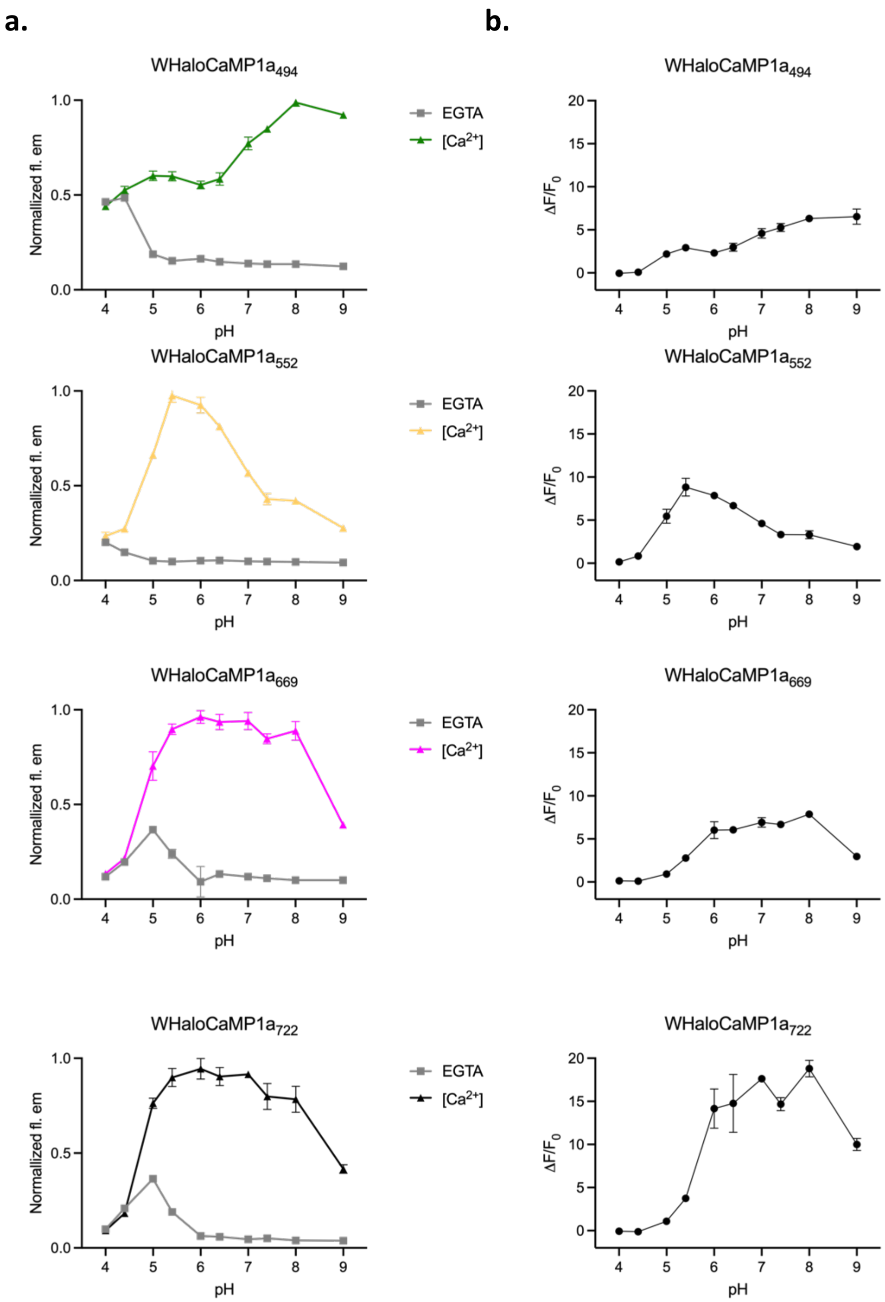

Supplementary Figure 15. pH stability of WHaloCaMP1a bound to different dyes.

**a**, Flourescence emission of WHaloCaMP1a bound to $JF_{494}$-HaloTag ligand, $JF_{552}$-HaloTag ligand, $JF_{669}$-HaloTag ligand and $JF_{722}$-HaloTag ligand in different pH values. Flourescence emission is normalized to the maximum flourescence emission. **b**, Calculated $\Delta F/F_0$ of WHaloCaMP1a bound to dyes at different pH.

| Protein scaffold | Dye/ supplement | Number of cells | ΔF/F (%) ± s.e.m | SNR | Time to peak (ms) | Decay $t_{1/2}$ (s) 1st comp. (Percentage 1st comp.) 2nd comp. |
|---|---|---|---|---|---|---|
| **WHaloCaMP1a** | | | | | | |
| | $JF_{494}$-HTL | 154 | 20 ± 4 | 18 ± 1 | 163 ± 5 | 0.4 ± 0.0 (59 ± 1%) 3.3 ± 0.1 |
| | $JF_{552}$-HTL | 165 | 4.4 ± 0.2 | 20 ± 1 | 178 ± 27 | 0.4 ± 0.0 (60 ± 1%) 5.7 ± 0.2 |
| | $JF_{669}$-HTL | 130 | 7.0 ± 0.4 | 20 ± 1 | 176 ±29 | 0.5 ± 0.1 (68 ± 1%) 5.1 ± 0.3 |
| | $JF_{722}$-HTL* | 21 | 13 ± 6 | 29 ± 3 | * | 1.0 ± 0.2 (55 ± 1%) 11 ± 1 |
| **iGECI** | with biliverdin (25 µM) | Reported from literature** | -12.9 | - | 700 | 14 (100%) |
| **iGECInano** | with biliverdin (25 µM) | Reported from literature*** | -21.7 | - | 700 | 2.4 (100%) |

*imaged at 10 Hz which is too slow to accurately capture time to peak.
**from (ref)[6]
***from (ref)[7]

Supplementary Fig/Table 16. Fluorescence response of WHaloCaMP1a or biliverdin-binding protein-based calcium indicators in a primary neuron culture field stimulation assay with one action potential stimulus.

Comparison of WHaloCaMP bound to different dyes-HaloTag ligands (HTL) and the brightest biliverdin-binding calcium indicators. Mean and standard error of the mean (s.e.m.) is reported for the number of neurons stated.

| Protein scaffold | Dye/ supplement | Number of cells | ΔF/F (%) ± s.e.m |
|---|---|---|---|
| **WHaloCaMP1a** | | | |
| | $JF_{494}$-HTL | 153 | 288 ± 0.1 |
| | $JF_{552}$-HTL | 168 | 60 ± 0.03 |
| | $JF_{669}$-HTL | 141 | 77 ± 0.03 |
| | $JF_{722}$-HTL* | 34 | 80 ± 0.03 |
| **iGECI**[#] | with biliverdin (25 µM) | Reported from literature[#] | -30.4 |

[#]from (ref)[6]

Supplementary Fig/Table 17. Fluorescence response of WHaloCaMP1a or biliverdin-binding protein-based calcium indicators in a primary neuron culture field stimulation assay with 160 action potentials.

Mean and standard error of the mean (s.e.m.) is reported for the number of neurons stated. HTL stands for HaloTag ligand.

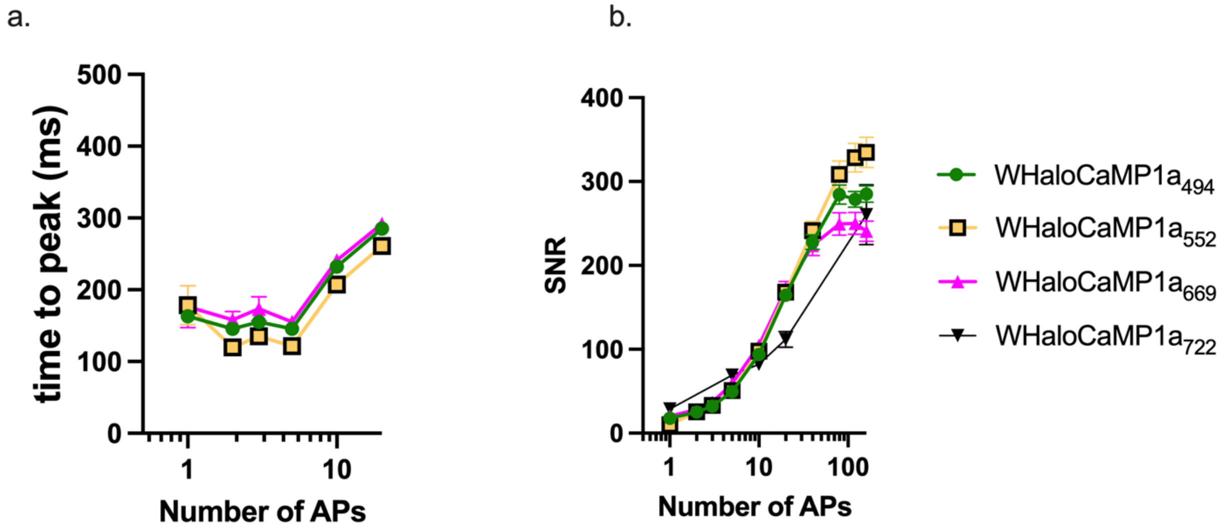

Supplementary Figure 18. Summary of properties of WHaloCaMP1a bound to different dye-ligands in a field stimulation assay in cultured neurons.

**a**, Time to peak (ms) from stimulation onset to max fluorescence value, imaged at 33 Hz. **b**, Signal-to-noise ratio (SNR) calculated as from the amplitude in the fluorescent change on the field stimulation divided by the standard deviation of the baseline fluorescence 1 s before each stimulus. Mean and standard error of the mean (s.e.m.) is shown.

WHaloCaMP1a$_{494}$

| Number AP | Traces fit | Percentage fast % ± s.e.m | $t_{1/2}$ fast (s) ± s.e.m. (s) | $t_{1/2}$ slow (s) ± s.e.m. (s) | Weighted average $t_{1/2}$ (s) |
|---|---|---|---|---|---|
| **1** | 147 | 59 ± 1 | 0.4 ± 0.0 | 3.3 ± 0.1 | 1.6 |
| **2** | 152 | 62 ± 1 | 0.4 ± 0.0 | 4.1 ± 0.1 | 1.8 |
| **3** | 153 | 63 ± 1 | 0.4 ± 0.0 | 4.5 ± 0.1 | 1.9 |
| **5** | 153 | 65 ± 1 | 0.4 ± 0.0 | 4.8 ± 0.1 | 1.9 |
| **10** | 152 | 67 ± 1 | 0.4 ± 0.0 | 4.9 ± 0.1 | 1.9 |
| **20** | 146 | 67 ± 1 | 0.4 ± 0.0 | 4.9 ± 0.1 | 1.9 |
| **40** | 120 | 65 ± 1 | 0.5 ± 0.0 | 4.9 ± 0.1 | 2.0 |

WHaloCaMP1a$_{552}$

| Number AP | Traces fit | Percentage fast % ± s.e.m | $t_{1/2}$ fast (s) ± s.e.m. (s) | $t_{1/2}$ slow (s) ± s.e.m. (s) | Weighted average $t_{1/2}$ (s) |
|---|---|---|---|---|---|
| **1** | 138 | 60 ± 1 | 0.4 ± 0.0 | 5.9 ± 0.3 | 2.6 |
| **2** | 155 | 60 ± 1 | 0.4 ± 0.0 | 5.7 ± 0.2 | 2.5 |
| **3** | 158 | 58 ± 1 | 0.4 ± 0.0 | 5.1 ± 0.2 | 2.4 |
| **5** | 164 | 56 ± 1 | 0.4 ± 0.0 | 4.9 ± 0.1 | 2.4 |
| **10** | 167 | 57 ± 1 | 0.4 ± 0.0 | 5.1 ± 0.1 | 2.4 |
| **20** | 167 | 56 ± 1 | 0.5 ± 0.0 | 5.0 ± 0.1 | 2.5 |
| **40** | 106 | 51 ± 1 | 0.5 ± 0.0 | 5.5 ± 0.1 | 3.0 |

WHaloCaMP1a$_{669}$

| Number AP | Traces fit | Percentage fast % ± s.e.m | $t_{1/2}$ fast (s) ± s.e.m. (s) | $t_{1/2}$ slow (s) ± s.e.m. (s) | Weighted average $t_{1/2}$ (s) |
|---|---|---|---|---|---|
| **1** | 118 | 68 ± 1 | 0.5 ± 0.1 | 5.1 ± 0.3 | 2.0 |
| **2** | 128 | 68 ± 1 | 0.3 ± 0.0 | 4.0 ± 0.1 | 1.5 |
| **3** | 129 | 66 ± 1 | 0.4 ± 0.0 | 4.0 ± 0.1 | 1.6 |
| **5** | 132 | 64 ± 1 | 0.5 ± 0.0 | 4.3 ± 0.1 | 1.8 |
| **10** | 141 | 62 ± 1 | 0.4 ± 0.0 | 4.3 ± 0.1 | 1.8 |
| **20** | 141 | 58 ± 1 | 0.4 ± 0.0 | 4.3 ± 0.1 | 2.0 |
| **40** | 141 | 54 ± 1 | 0.5 ± 0.0 | 4.6 ± 0.4 | 2.4 |

WHaloCaMP1a$_{722}$

| Number AP | Traces fit | Percentage fast % ± s.e.m | $t_{1/2}$ fast (s) ± s.e.m. (s) | $t_{1/2}$ slow (s) ± s.e.m. (s) | Weighted average $t_{1/2}$ (s) |
|---|---|---|---|---|---|
| **1** | 23 | 55 ± 1 | 1.0 ± 0.2 | 11 ± 1 | 5.5 |
| **5** | 21 | 46 ± 1 | 0.8 ± 0.0 | 10 ± 1 | 5.7 |
| **10** | 34 | 41 ± 2 | 0.7 ± 0.0 | 8.3 ± 0.5 | 5.1 |
| **20** | 31 | 42 ± 1 | 1.0 ± 0.1 | 10 ± 1 | 6.2 |

Supplementary Fig/Table 19. Decay properties calculated for WHaloCaMP1a bound to different dyes in a field stimulation assay in cultured neurons.

Decay was calculated from the end of the stimulus given. Mean and standard error of the mean (s.e.m.) is reported for the number of neurons stated.

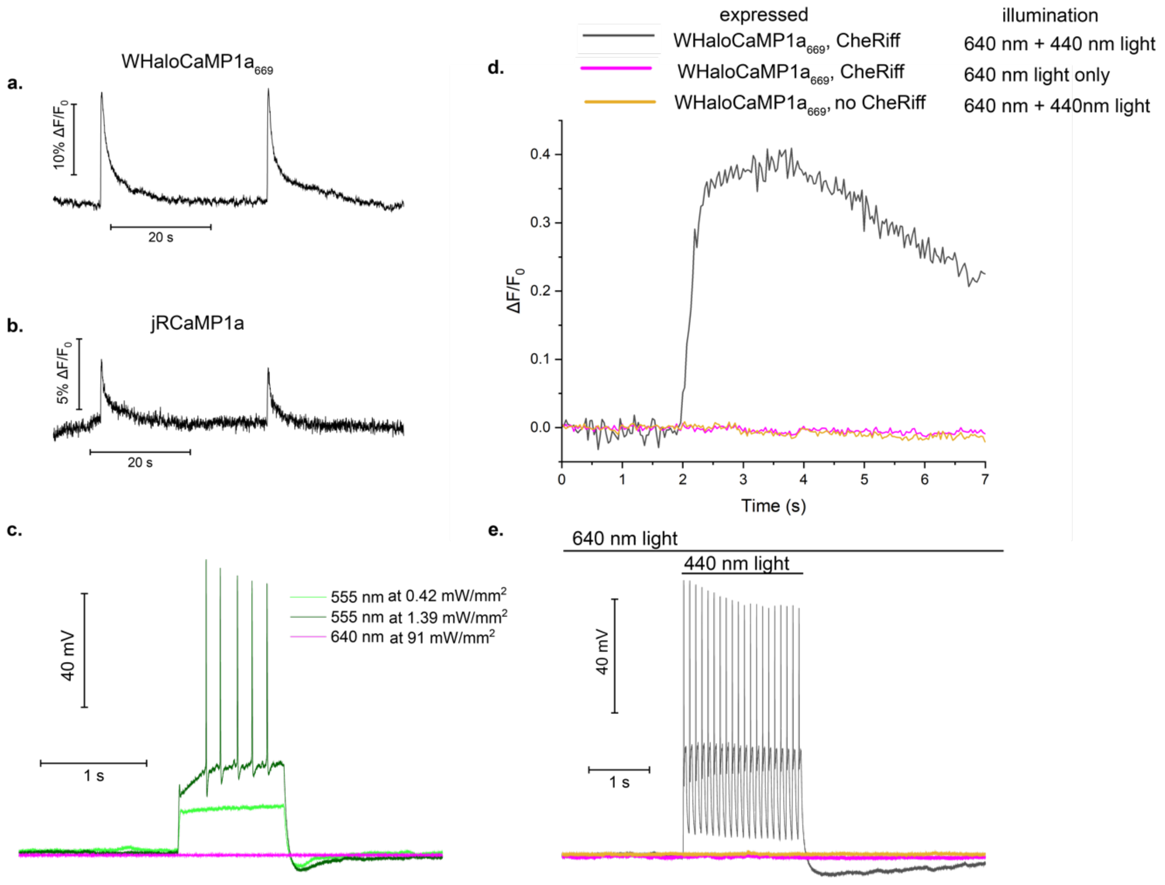

Supplementary Figure 20. WHaloCaMP1a$_{669}$ is spectrally compatible with the channelrhodopsin variant CheRiff.

Primary neuronal cultures were infected with AAV particles for WHaloCaMP1a and labeled with JF$_{669}$-HaloTag ligand, or infected with AAV particles for with jRCaMP1a, to measure spectral compatibility with the blue light-activated channel rhodopsin CheRiff via fluorescence imaging and simultaneous electrophysiology. **a**, Fluorescence trace of WHaloCaMP1a$_{669}$ after current injection, imaged with low light power (0.76 mW/mm$^2$, 640 nm LED). **b**, Fluorescence trace of jRCaMP1a after current injection, imaged with low light power (1.39 mW/mm$^2$, 555 nm LED). **c**, neuronal cultures expressing CheRiff, with illumination at 555 nm or 640 nm. 555 nm light depolarizes the cell at low light levels, even if it does not lead to spiking action potentials. No depolarization was observed with 640 nm illumination, even with over 10× the light power needed to image WHaloCaMP1a$_{669}$ with good SNR. **d** and **e**, WHaloCaMP1a$_{669}$ is compatible with CheRiff activation and does not show any 440 nm light dependent-photoswitching. Fluorescence trace (**d.**) and electrophysiological recoding (**e.**) are shown for the same experiment. Action potentials and changes in fluorescence are only observed when WHaloCaMP1a$_{669}$ is co-expressed with CheRiff and illuminated with 440 nm and 640 nm light (black trace). No action potentials or change in fluorescence are seen when only illuminated with 640 nm light, or when WHaloCaMP1a$_{669}$ is expressed alone.

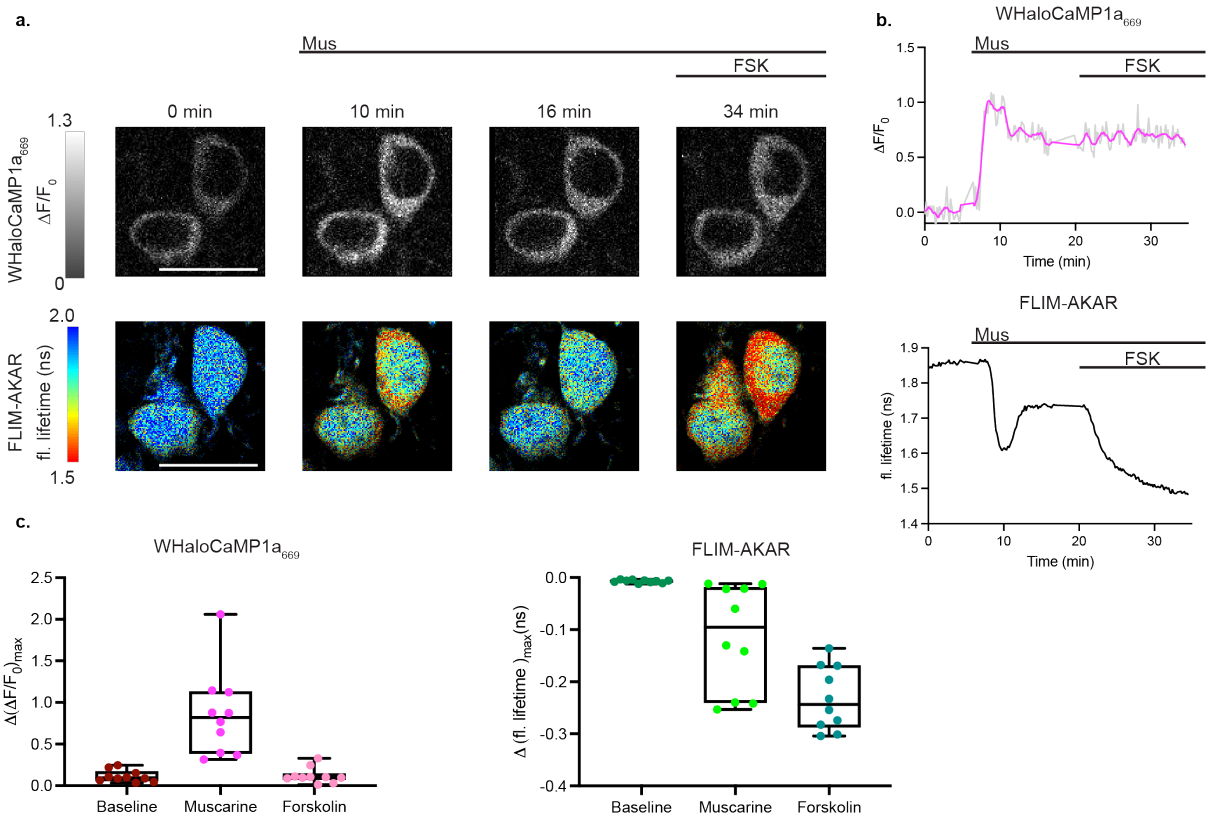

Supplementary Figure 21. Multiplexed imaging in acute brain slices with WHaloCaMP1a$_{669}$ and FLIM-AKAR.

**a,** Heatmap showing multiplexed imaging of WHaloCaMP1a$_{669}$ and FLIM-AKAR. A decrease in fluorescence lifetime in FLIM-AKAR indicates an increase in PKA activity. Top row: intensity heatmap of hippocampal CA1 neurons in acute brain slices expressing WHaloCaMP1a$_{669}$, showing calcium responses to muscarine (mus, 10 µM) and subsequent forskolin (FSK, 50 µM) application. Bottom row: lifetime heatmap of FLIM-AKAR of the cells at the same time points showing PKA phosphorylation responses. Scale bar, 20 µm. **b**, Example traces of WHaloCaMP1a$_{669}$ and cytoplasmic FLIM-AKAR plotted through time. The intensity and lifetime traces correspond to the top right cell in panel a. The gray intensity trace corresponds to raw $\Delta F/F_0$ values, and the trace in magenta corresponds to $\Delta F/F_0$ values smoothed through time using 6 neighboring points and a second order smoothening polynomial in GraphPad Prism. **c**, Summary of maximum responses of cells co-expressing WHaloCaMP1a$_{669}$ and cytoplasmic FLIM-AKAR. Each circle represents one hippocampal CA1 or cortical neuron. Each box represents median and interquartile intervals. The whiskers represent the range of data.

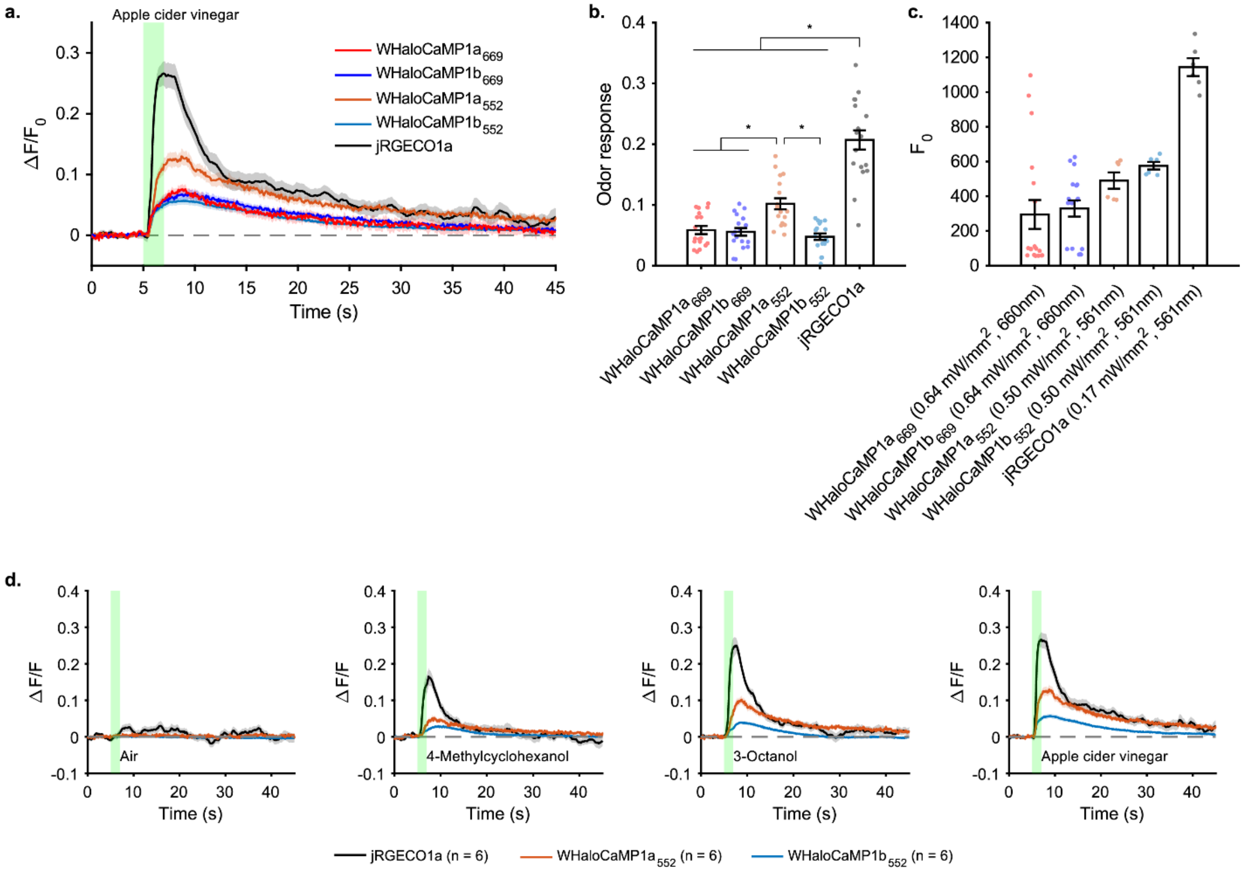

Supplementary Figure 22. Odor responses in fruit flies recorded with different WHaloCaMP variants and JF dye-ligands.

**a**, Fluorescence time courses to apple cider vinegar (ACV) odor. WHaloCaMP-expressing flies **a**, Fluorescence time courses to apple cider vinegar odor. WHaloCaMP-expressing flies were loaded with JF-dye HaloTag ligands (5 µM JF$_{669}$-HTL or 1 µM JF$_{552}$-HTL) via a 1hr incubation/1hr wash-out protocol. n = 6 flies for each group and each fly received three odor presentation trials. **b**, Mean odor responses integrated during the time window from 0.5 and 5.5 seconds after odor onset. Data points are odor trials, while bars are means ± standard error of the mean (s.e.m.). *, $P < 0.01$. t-test with Bonferroni correction of multiple comparisons. Same data as in (**a.**). **c**, Resting florescence ($F_0$). Bars are means ± s.e.m. from odor trials. LED wavelength and intensity at the sample plane are noted. n = 2-6 flies for each group. **d**, Responses of WHaloCaMP1a$_{552}$ and WHaloCaMP1b$_{552}$ in comparison to jRGECO1a for different odor presentations for n = 6 flies. Green shading indicates odor presentation.

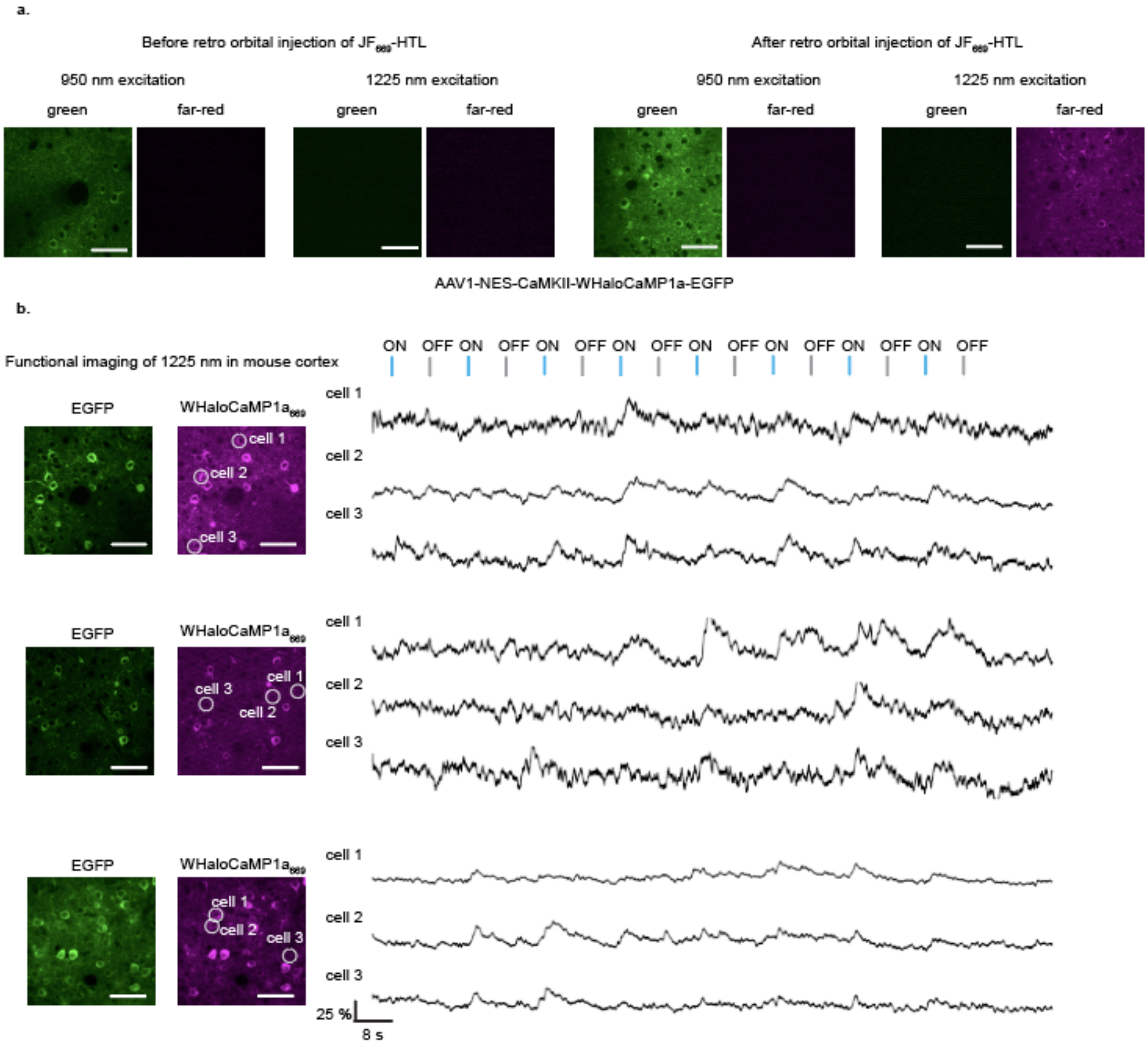

Supplementary Figure 23. Two-photon imaging of WHaloCaMP1a labeled with $JF_{669}$-HaloTag ligand in mouse visual cortex.

**a**, Two photon imaging of mouse cortex expressing WHaloCaMP1a-EGFP with excitation at 950 nm and 1225 nm before and after retro orbital injection with $JF_{669}$-HaloTag ligand. **b**, functional imaging of $WHaloCaMP1a_{669}$ in mouse visual cortex. Left, images of EGFP and $WHaloCaMP1a_{669}$ imaging channels and right, fluorescence $\Delta F/F_0$ traces of highlighted neurons during visual presentation stimulus in front of the mouse eye as indicated by the “ON” and “OFF” indications above the traces. All scale bars, 50 µm.

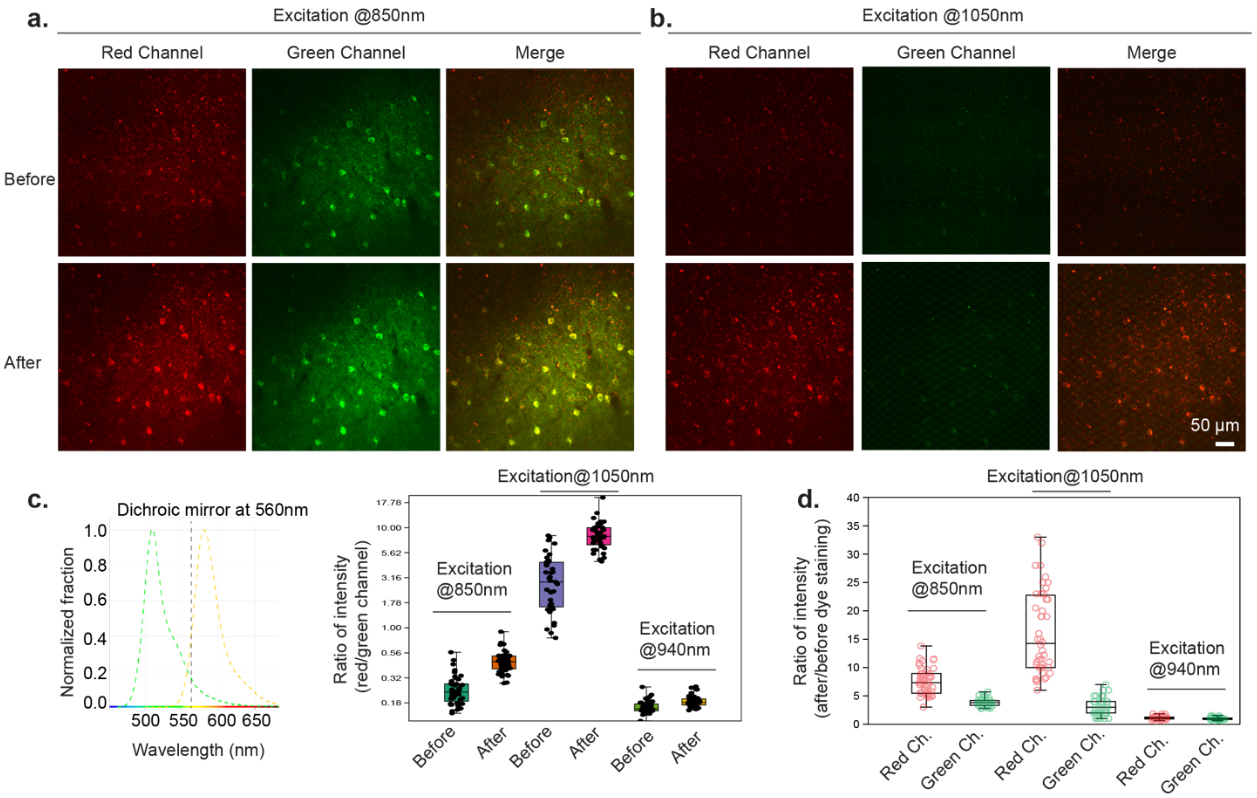

Supplementary Figure 24. Two-photon imaging of WHaloCaMP1a$_{552}$ in mouse primary visual cortex (V1).

**a** and **b**, Images acquired under excitation at 850 nm (**a.**) or 1050 nm (**b.**) Images of the same field-of-view were acquired before (top) and after (bottom) the injection of JF$_{552}$-HaloTag ligand. Scale bar, 50 µm. **c**, Left: Emission spectrum of EGFP and JF$_{552}$ -HaloTag ligand (https://public.brain.mpg.de/shiny/apps/SpectraViewer/). Dashed line indicates the dichroic mirror cutoff wavelength. Right: Quantification of changes in the ratio of intensity (red over green channels) before and after dye injection at 850 nm, 1050 nm and 940 nm 2-photon excitation wavelengths. Y axis is displayed in $\log_{10}$ for better visualization of data trend. Median values/cell numbers are (from left to right): 0.23/45; 0.46/45; 2.87/39; 8.2/41; 0.16/46; 0.18/46. Each dot represents a cell. **d**, Quantification of the ratio of intensity (after dye labeling over before dye labeling) at the three different excitation wavelengths (850 nm, 1050 nm and 940 nm). Median values/cell numbers are (from left to right): 7.57/48; 3.81/48; 14.25/44; 3.0/44; 1.10/49; 0.94/49. Each dot represents a cell.

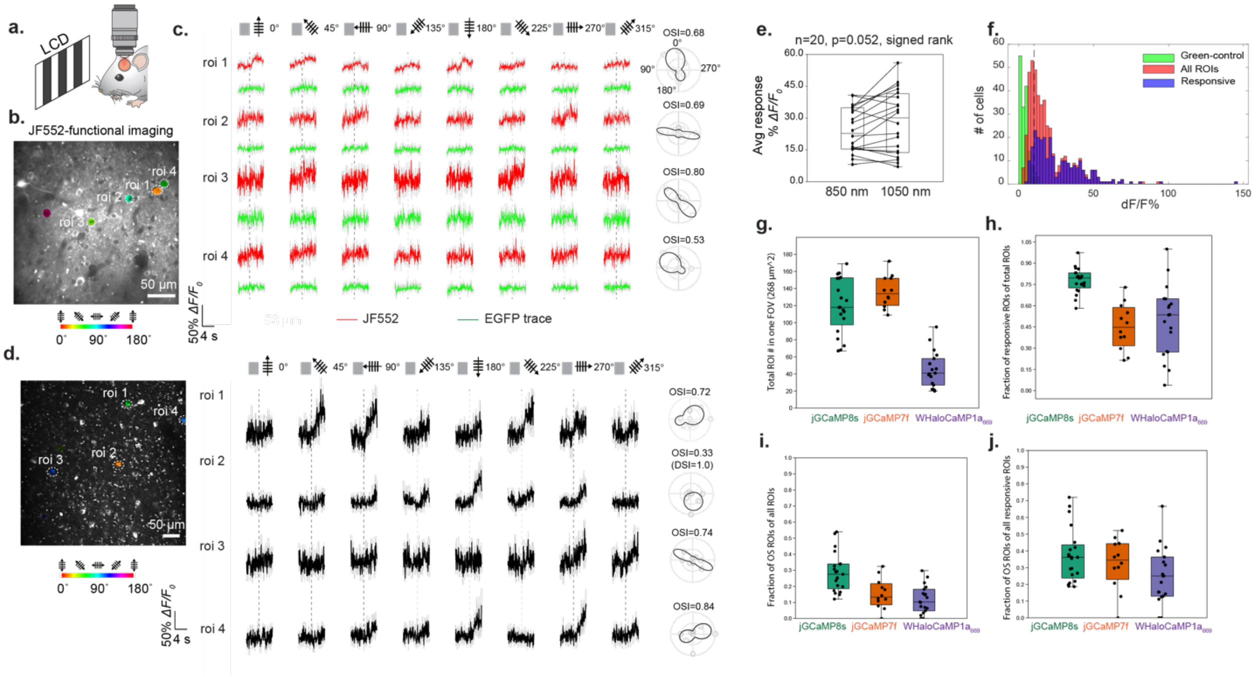

Supplementary Figure 25. Functional imaging in the mouse V1 of WHaloCaMP1a$_{552}$ and quantitative analysis.

**a**, Schematic of the experimental setup: the animal is anesthetized and exposed to moving grafting of different orientations. **b** and **c**, Orientations selectivity (OS) map (**b.**) and calcium (JF$_{552}$ channel) or control (EGFP channel) traces (ROIs 1-4) under 805 nm excitation (**c.**). **d**, OS map (left) and calcium (JF$_{552}$ channel) traces (ROIs 1-4) under 1050 nm excitation. Imaging rate is 15 Hz. Orientation selectivity index (OSI) of cells were shown in the right panel of (**c, d**). **e**, Side-by-side comparison of response magnitude under 850 nm and 1050 nm excitation for the same cells. N = 20 cells. $p$ = 0.052. Signed rank test. **f**, Distribution of response magnitude in green-control channel (ΔF/F % as 10.2% at 95 percentile shown as dashed line, n = 160 cells from 2 mice), all ROIs in the JF$_{552}$ channel (ΔF/F % as 25.5% at 75 percentile, n = 470 cells from 4 mice) and responsive ROIs in JF$_{552}$ channel (ΔF/F % as 35.4% at 75 percentile, n = 260 cells from 4 mice). **g-j,** Comparison of three calcium indicators in terms of **g**, total number of recognizable ROI; **h**, fraction of responsive ROIs; **i**, fraction of OS cells of all ROIs; **j**, fraction of OS cells of all responsive cells. Fractions of responsive or orientation-selective cells from 19 field-of-views in 4 mice for WHaloCaMP1a$_{669}$. For jGCaMP7f and jGCaMP8s, data were from 12 FOVs in 3 mice and from 21 FOVs in 5 mice, respectively, which is the same dataset as used in one of our recent publications[9].

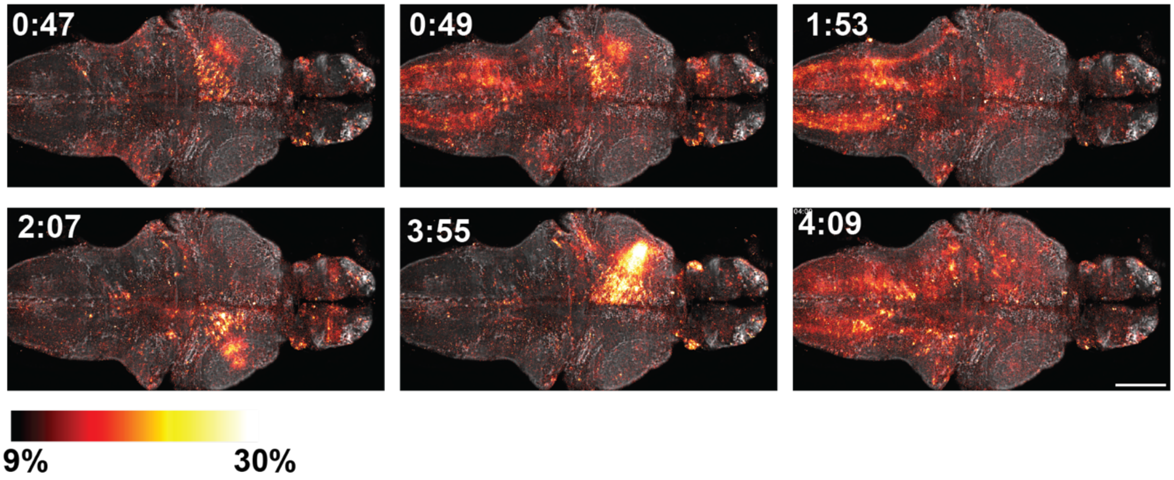

Supplementary Figure 26. Volumetric whole brain imaging of neurons in zebrafish larvae using SiMView.

Tg(*elavl3:*NES-WHaloCaMP1a-EGFP) zebrafish were labeled with $JF_{669}$-HaloTag ligand and mounted in a SiMView light sheet microscope. Three-dimensional volumes of the brain with a 160 µm length along the z-axis were acquired with a z-step size of 4 µm and a rate of 4 volumes per second. Visualization of $\Delta F/F_0$ (shown using a red-hot color look-up table) displayed over a projection of the concatenated projected ΔF/F time series that serves as an anatomical reference (shown in grey). Time stamps are in min:s and scale bar is 100 µm.

A movie of this whole brain time-lapse image data set can be found in the supplement.

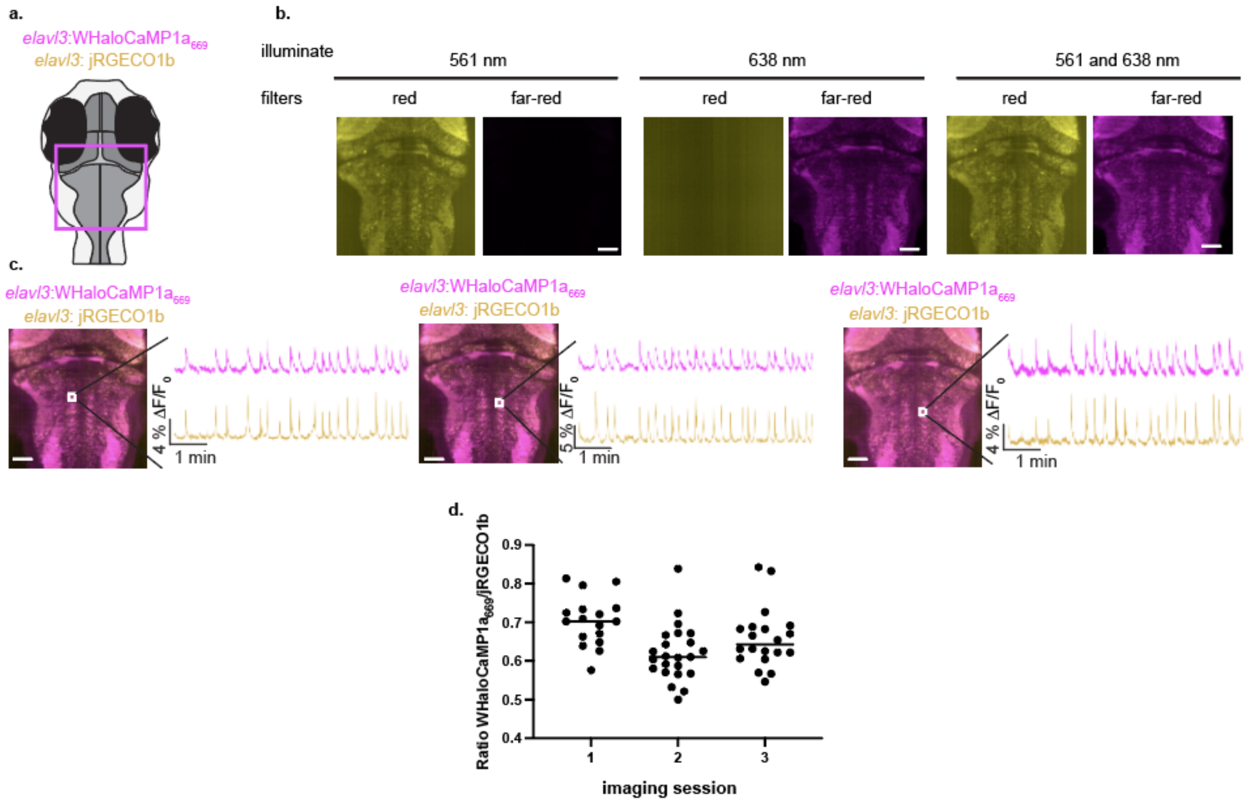

Supplementary Figure 27. Comparison of WHaloCaMP1a669 and jRGECO1b during light sheet imaging of neurons in zebrafish larvae.

**a**, Schematic of a zebrafish larva to indicate with an magenta box where imaging was performed. **b**, Fluorescence images of Tg(*elavl3*: NES-WHaloCaMP1a-EGFP), Tg(*elavl3*: jRGECO1b), with single laser illumination of 561 nm or 638 nm to excite jRGECO1b or WHaloCaMP1a669, respectively, or simultaneous illumination for dual color imaging. **c**, Functional imaging of WHaloCaMP1a669 and jRGECO1b in neurons of zebrafish larvae with comparison of the $\Delta F/F_0$ fluorescence traces from the two channels. Three imaging sessions from two fish are shown. **d**, Quantitative comparison between the WHaloCaMP1a669 channel and jRGECO1b channel for each $Ca^{2+}$ transient in the fluorescence traces. Each dot is a $Ca^{2+}$ transient in the fluorescence trace. Overall, WHaloCaMP1a669 showed 66% of the jRGECO1b signal during the $Ca^{2+}$ transients. All scale bars, 50 µm.

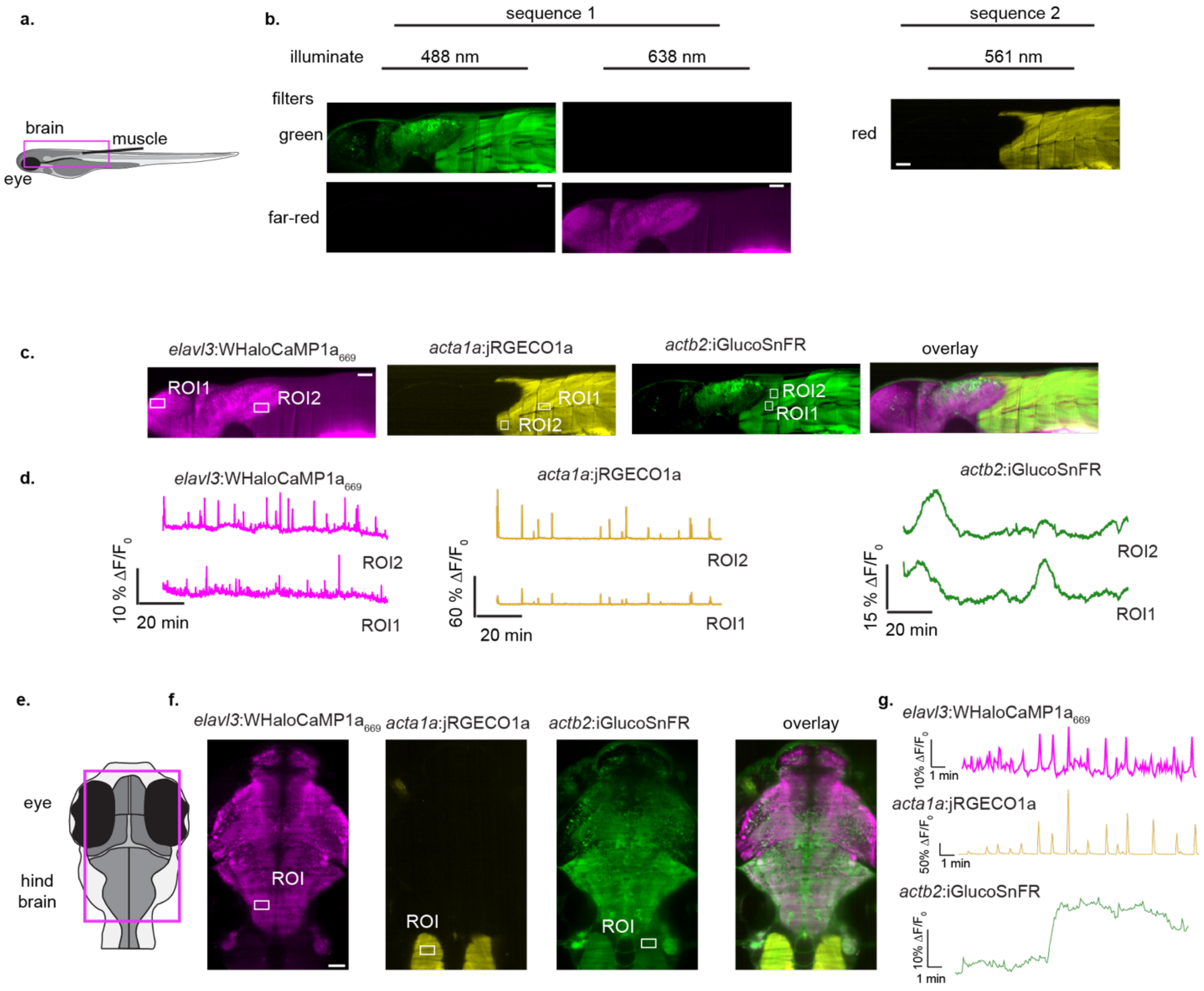

Supplementary Figure 28. Three-color functional multiplexed imaging in zebrafish larvae with WHaloCaMP1a$_{669}$, jRGECO1a and iGlucoSnFR.

**a**, Schematic of a zebrafish larva from the side view, highlighting the eye, brain, and muscle. **b**, Fluorescence images from sequential illumination sequences of sequence 1: 488 nm and 638 nm excitation; sequence 2: 561 nm excitation with representative images showing clear separation of the fluorescence channels. **c**, Fluorescence images of zebrafish larvae used in three color multiplexed functional imaging, with ROI for further analysis highlighted. **d**, Fluorescence $\Delta F/F_0$ traces from ROIs shown in (**c.**) ROI2 in the neuronal WHaloCaMP1a$_{669}$ channel correlates well with the muscle jRGECO1a channel, while ROI1 does not. iGlucoSnFR fluorescence changes were slower than the $Ca^{2+}$ transients in neurons and muscle. **e**, Schematic of a zebrafish larva for three color multiplexed imaging. **f**, Fluorescence images of zebrafish larvae used in three color multiplexed functional imaging, with an ROI for further analysis highlighted. **g**, Fluorescence $\Delta F/F_0$ traces from ROI shown in (**f.**). All scale bars, 50 µm.

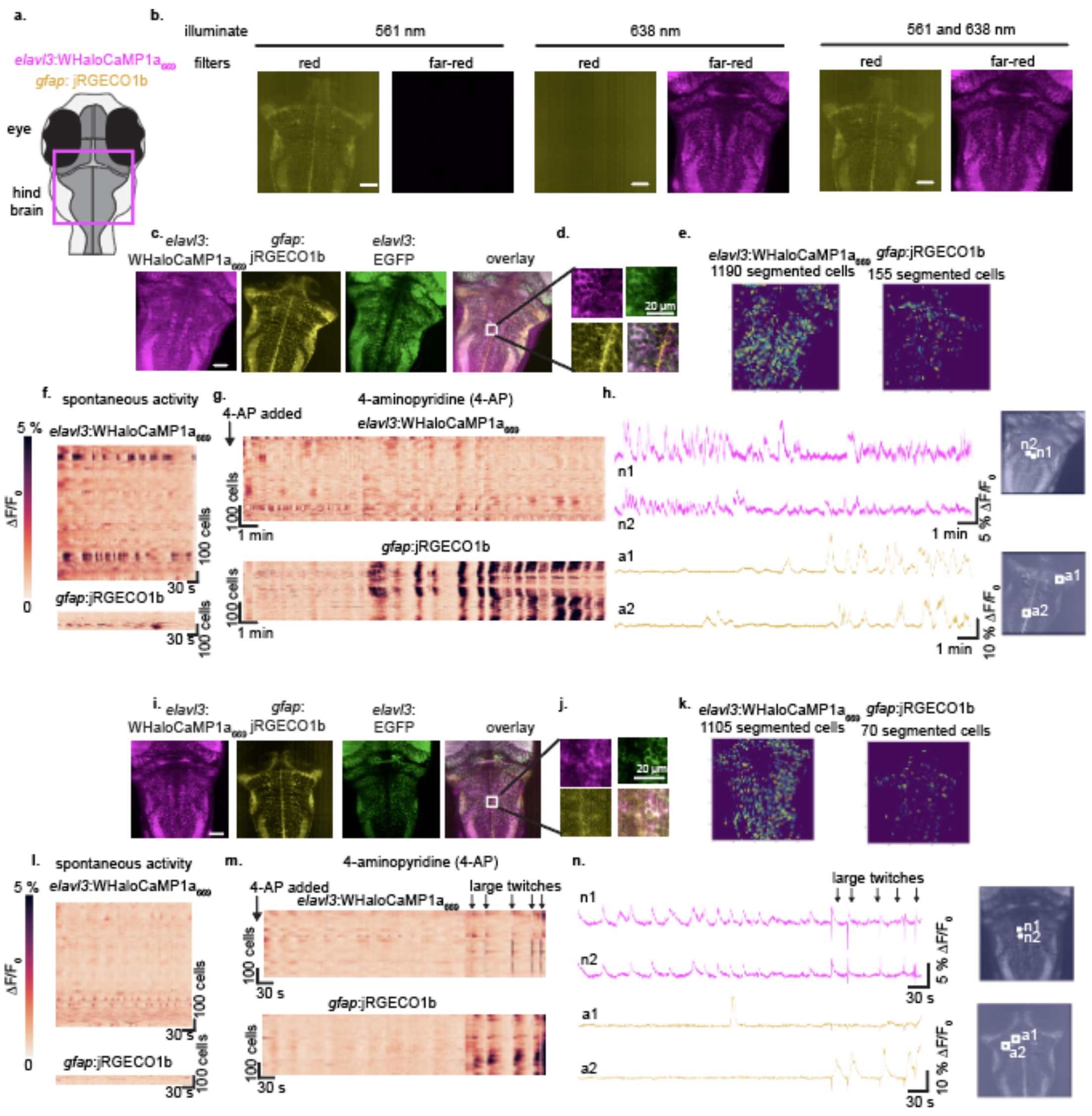

Supplementary Figure 29 .Dual-color functional imaging of astrocyte and neuronal $Ca^{2+}$ in zebrafish larvae during spontaneous activity or in the presence of 4-aminopyridine.

**a**, Schematic showing hind brain region where functional imaging was conducted. **b**, fluorescence images of single illumination of 561 nm or 638 nm, and dual illumination of 561 nm and 638 nm. Clear separation of channels for dual color imaging is possible. **c** and **i**, Fluorescence images of zebrafish larvae expressing NES-WHaloCaMP1a-EGFP in neurons and labeled with $JF_{669}$-HaloTag ligand and jRGECO1b in astrocytes. **d** and **j**, Zoom in of (**c.**) and (**i.**) showing single cell resolution. **e** and **k**, Suite2P and CellPose segmentation of dual color functional imaging. **f**, **g**, and **l**, **m**, Rasterplot of fluorescence changes in neuronal $WHaloCaMP1a_{669}$ (top) and astrocyte jRGECO1b (bottom), during spontaneous activity (**f. and l.)** or after addition of 4-AP (**g. and m.**). **h** and **n**, Two neurons and two astrocytes highlighted fluorescence $\Delta F/F_0$ trace. Functional imaging in (m.) and (**n.**) was cut short by large twitches during imaging. Unless otherwise stated, scale bars, 50 µm.

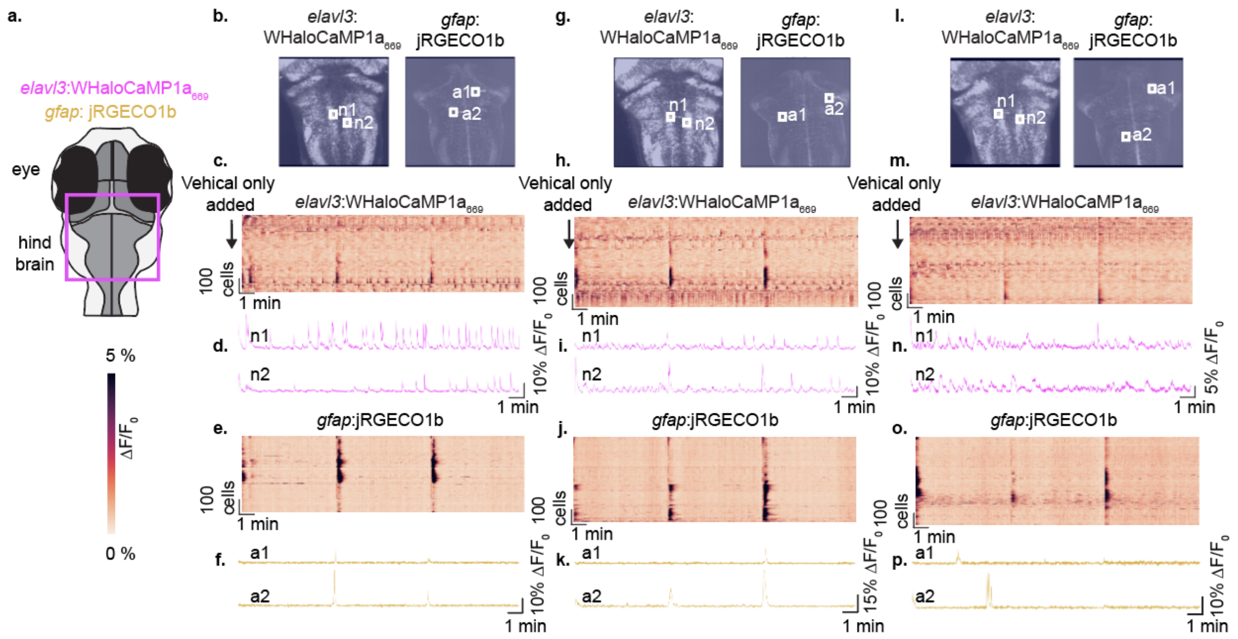

Supplementary Figure 30. Dual-color functional imaging of astrocyte and neuronal $Ca^{2+}$ in zebrafish larvae over longer time in the absence of 4-aminopyridine.

To validate that we could follow $Ca^{2+}$ in neurons reported by $WHaloCaMP1a_{669}$ for up to 18 minutes, we performed light sheet imaging as described in the main text on Tg(*elavl3*:NES-WHaloCaMP1a) crossed with Tg(*gfap*:jRGECO1b), but without the addition of 4-AP. **a**, Schematic of zebrafish larvae showing hind brain. **b**, **g** and **l**, images of $WHaloCaMP1a_{669}$ and jRGECO1b in three individual fish, indicating neurons and astrocyte that are later followed. **c**, **h** and **m**, Rastermap heatmap of neuronal activity as reported by $WHaloCaMP1a_{669}$. Features can be distinguished during the full 18-minute imaging time. **d**, **l** and **n**, Fluorescence $\Delta F/F_0$ traces from highlighted neurons, part of the hind brain oscillator, which we could distinguish during the full 18-minute imaging time. **e**, **j** and **o**, Rastermap heatmap of astrocyte activity as reported by jRGECO1b. No large waves of activity were seen. **f**, **k** and **p**, Fluorescence $\Delta F/F_0$ traces from highlighted astrocytes. Images were acquired as three time series of 6 min 14 s and concatenated for analysis.

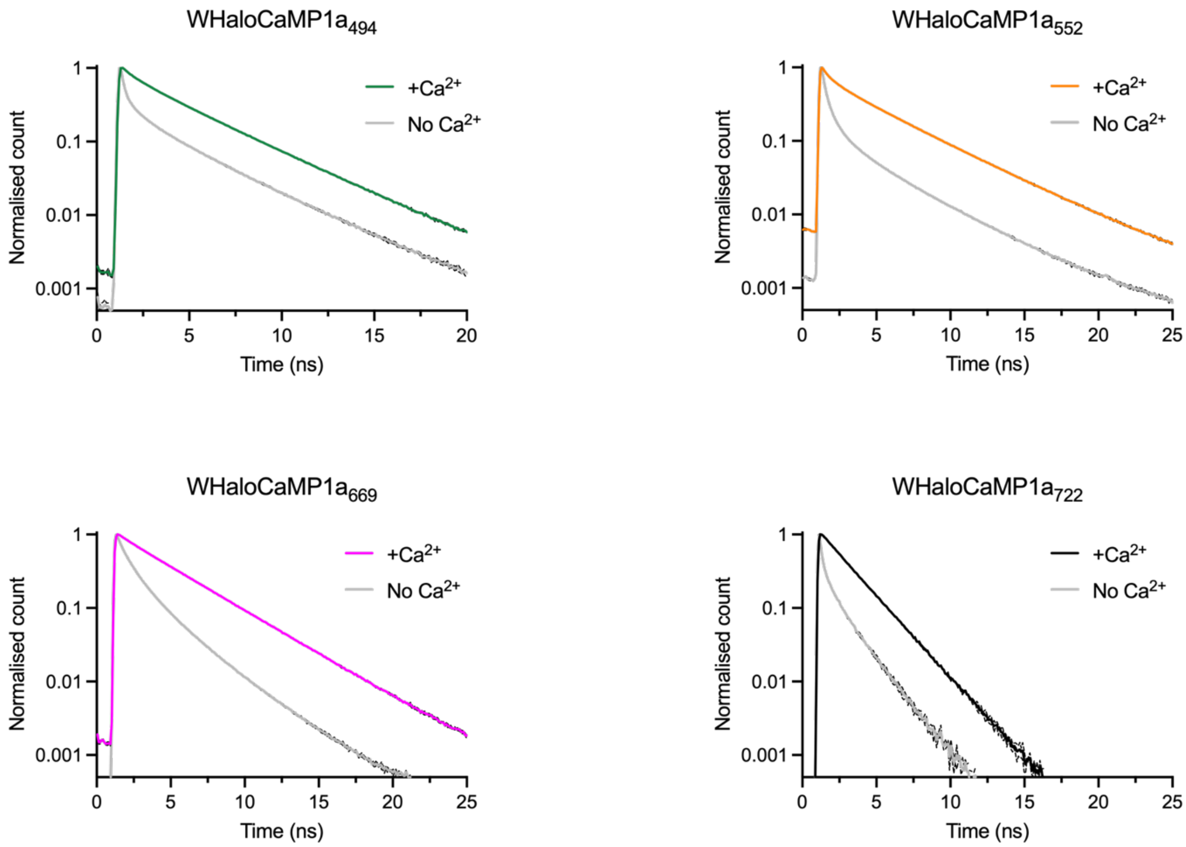

| | $Ca^{2+}$ lifetime (ns) | EGTA lifetime (ns) | Delta lifetime (ns) |
|---|---|---|---|
| **WHaloCaMP1a$_{494}$** | 2.89 ± 0.03 | 0.73 ± 0.01 | 2.02 |
| **WHaloCaMP1a$_{552}$** | 2.87 ± 0.03 | 0.86 ± 0.01 | 2.15 |
| **WHaloCaMP1a$_{669}$** | 3.43 ± 0.04 | 1.27 ± 0.01 | 2.15 |
| **WHaloCaMP1a$_{722}$** | 1.87 ± 0.01 | 0.49 ± 0.01 | 1.34 |

Supplementary Figure 31. Fluorescence lifetime image microscopy (FLIM) with WHaloCaMP1a.
Fluorescence lifetime image microscopy (FLIM) with WHaloCaMP1a bound to JF$_{494}$-HaloTag ligand, JF$_{552}$-HaloTag ligand, JF$_{669}$-HaloTag ligand and JF$_{722}$-HaloTag ligand. A time domain FLIM set-up was used at 40 MHz. The lifetime decay was fit to a three-component fit and the amplitude weighted lifetime is reported. Mean and standard deviation from three replicates from at least two different samples.

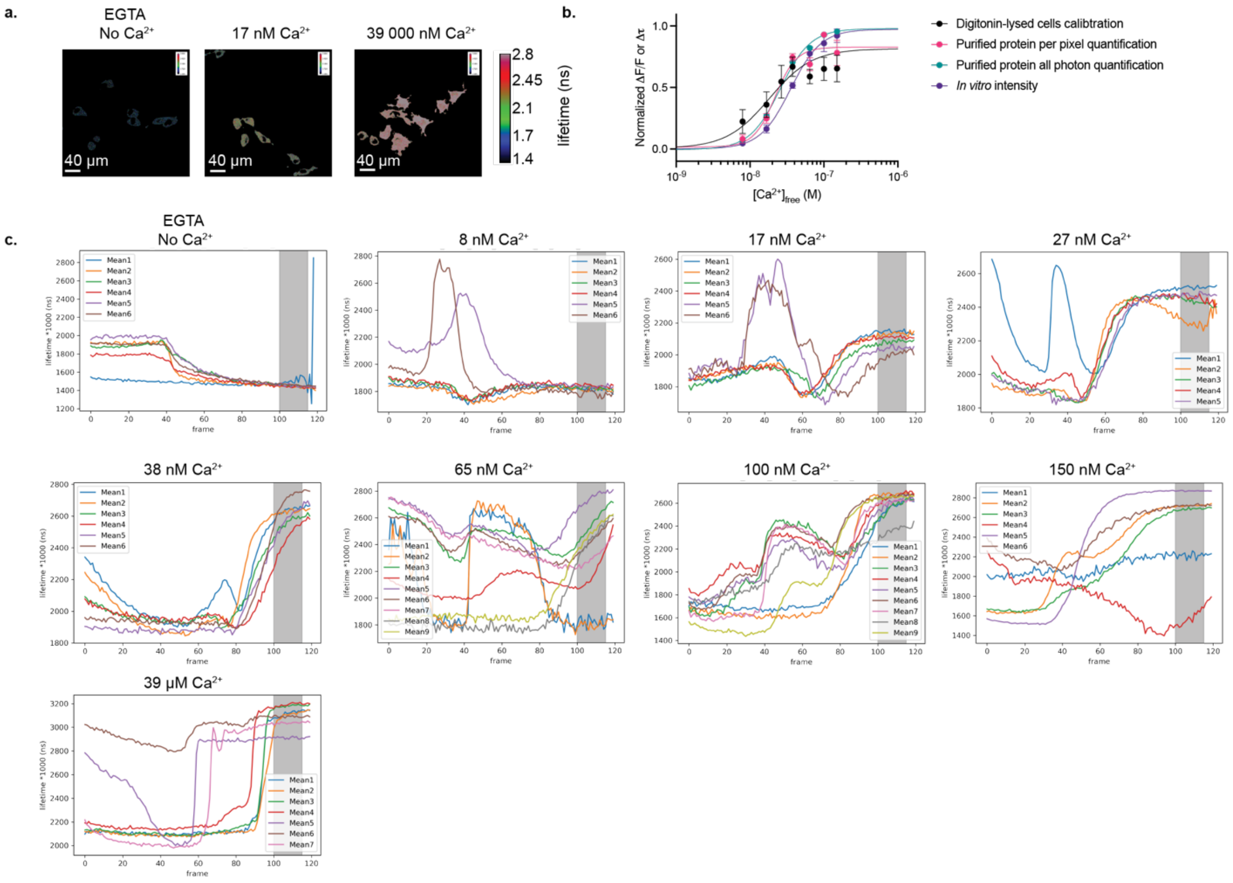

fit-parameters from calibration curves of normalized $\Delta F/F_0$ or $\Delta\tau$ vs. $[Ca^{2+}]$in (**b**.).

| Fit | $EC_{50}$ (nM) | Hill coeff. |
|---|---|---|
| **Digitonin-lysed cells calibration (lifetime) (black)** | 18 ± 5 | 1.5 ± 0 |
| **Purified protein per pixel quantification (lifetime) (pink)** | 22 ± 3 | 3 ± 0.8 |
| **Purified protein all photon quantification(lifetime) (green)** | 26 ± 1 | 2.1 ± 0.1 |
| ***In vitro* intensity (purple)** | 35 ± 2 | 2.0 ± 0.2 |

*the hill coefficient was constrained to greater than 1.5.

### Supplementary Figure 32. Calibration of WHaloCaMP1a$_{669}$ for quantitative $[Ca^{2+}]$ determination by FLIM.

**a**, FLIM images from HeLa cells transfected with WHaloCaMP1a and labeled with $JF_{669}$-HaloTag ligand. Cells were bathed in known concentration of free calcium solution and permeabilized by digitonin. Three representative images are shown for No $Ca^{2+}$, 17 nM $Ca^{2+}$ and 39 µM $Ca^{2+}$. Scalebar, 40 µm. **b**, Fluorescence lifetime vs. $[Ca^{2+}]$ titration calibration curves from digitonin lysed cells (black), purified protein per pixel lifetime fit (pink), purified protein all-photon lifetime fit (green), or purified protein plate reader intensity experiment. All titrations are normalized to max value within the titration curve. Fit values from the titrations curve in the table below the images. **c**, Quantified lifetime from cells after permeabilization with digitonin and bathed in indicated concentration of free calcium. Shaded area indicates region which was used to generate the calibration curve. Representative example from three independent calibration experiments.

To calculate the [$Ca^{2+}$] (x, M) from a measured lifetime (y, ns, scaled by 1000) for images acquired under imaging parameters used for the HeLa cell calibration experiment:

$$x = EC_{50}\left(\frac{a-y}{y-b}\right)^{\left(\frac{1}{h}\right)}$$

| a (*1000) (ns) | b (*1000) (ns) | $EC_{50}$ (M) | h |
|---|---|---|---|
| 1300 | 2900 | $1.6 \times 10^{-8}$ | 1.5 |

Where (a) is the value of fluorescence at the bottom of the calibration curve, (b) is the value of fluorescence at the top of the curve, ($EC_{50}$) is the concentration of agonist that gives a response halfway between bottom and the top, and (h) is the hill or cooperative coefficient.

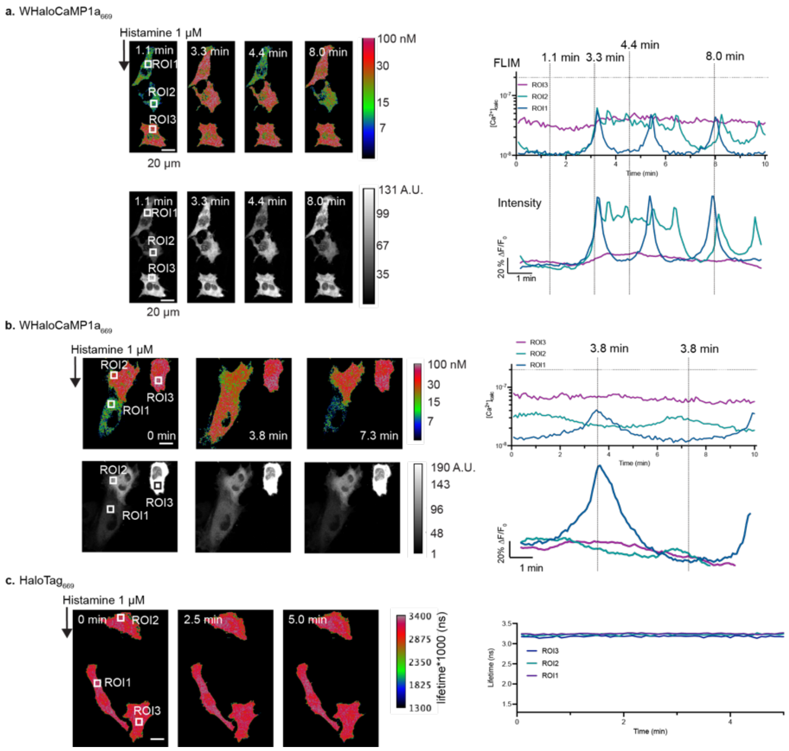

Supplementary Figure 33. WHaloCaMP1a$_{669}$ as a FLIM probe in HeLa cells.

Equation used to calculate [$Ca^{2+}$] from a given lifetime (scaled by 1000 ns), as well as the fit constant values used, derived from in-cell titration with permeabilized HeLa cells and known free $Ca^{2+}$ solutions. **a** and **b**, Two further examples of WHaloCaMP1a$_{669}$ in HeLa cells used to determine $Ca^{2+}$ concentrations during histamine-induced oscillations. Example images of the lifetime images converted to [$Ca^{2+}$] (top) and intensity images (bottom) to the left, as well as traces of calculated [$Ca^{2+}$] from the fluorescence lifetime and fluorescence $\Delta F/F_0$ traces for three ROIs highlighted in the images. The maximum concentration of [$Ca^{2+}$] that could be quantified is indicated by a horizontal line on the y-axis (200 nM), and vertical lines indicate time points of images in the left representative images. **c**, HaloTag$_{669}$ expressed in HeLa cells and stimulated with histamine shows no changes in fluorescence lifetime. Scale bar, 20 µm.

To calculate the $[Ca^{2+}]$ (x, M) from a measured lifetime (y, ns, scaled by 1000 ns) for the images acquired under images parameters *for in vivo* zebrafish larvae imaging:

$$x = EC_{50}\left(\frac{a-y}{y-b}\right)^{\left(\frac{1}{h}\right)}$$

| a (*1000) (ns) | b (*1000) (ns) | $EC_{50}$ (M) | h |
|---|---|---|---|
| 1400 | 2900 | $3.8 \times 10^{-8}$ | 2.1 |

Where (a) is the value of fluorescence at the bottom of the calibration curve, (b) is the value of fluorescence at the top of the curve, ($EC_{50}$) is the concentration of agonist that gives a response halfway between bottom and the top, and (h) is the hill or cooperative coefficient.

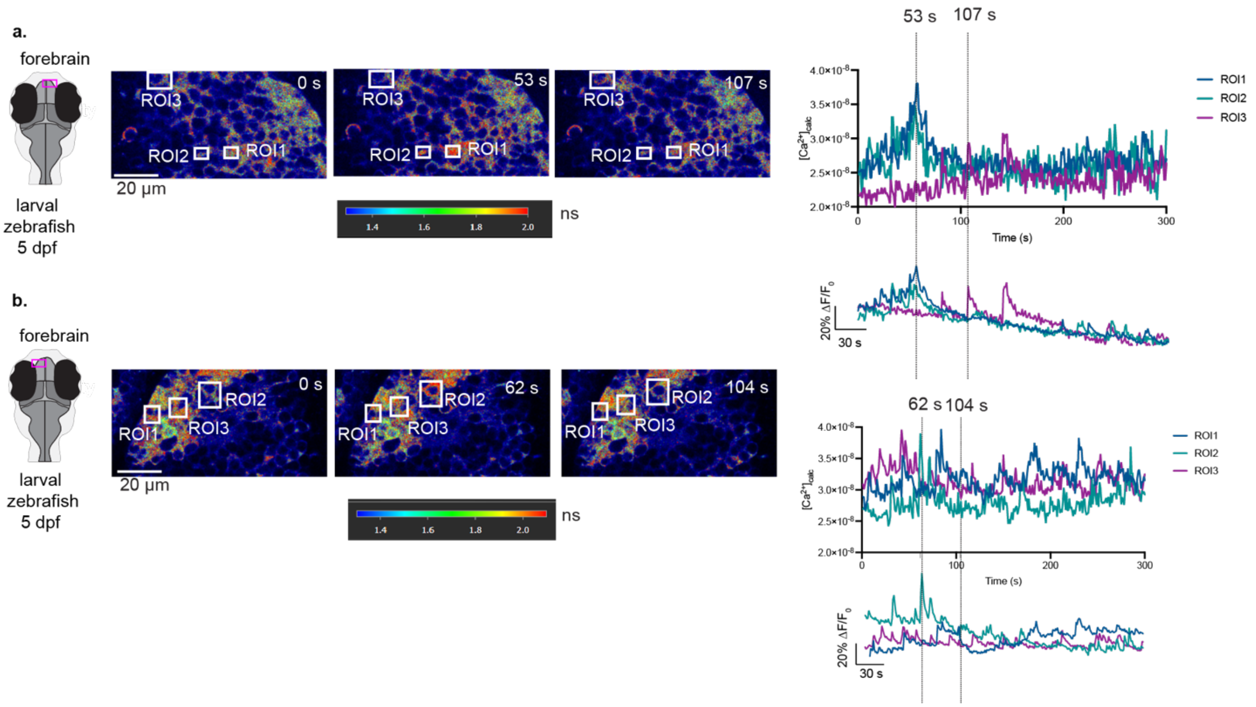

Supplementary Figure 34. *In vivo* quantitative FLIM for $[Ca^{2+}]$ in zebrafish larvae.
Equation used to calculate $[Ca^{2+}]$ from a given lifetime (scaled by 1000 ns), as well as the fit constant values used, derived from purified protein calibration at these image settings. **a** and **b**, Two further examples of in vivo FLIM to determine $[Ca^{2+}]$ in zebrafish larvae on the single cell resolution. Tg(*elavl3*:NES-WHaloCaMP1a-EGFP) was labeled with $JF_{669}$-HaloTag ligand and zebrafish larvae mounted on an inverted FLIM set-up, with zoom in on the forebrain as indicated in the schematic. FLIM images and intensity images overlaid in the Leica LASX software are shown, with the color bar for lifetime. Analysis was done on raw images of lifetime, which were used to calculate the $[Ca^{2+}]$ from a calibrated $Ca^{2+}$ titration in purified protein. $[Ca^{2+}]$ calculated from fluorescence lifetimes (top) in ROIs highlighted in the images, as well as fluorescence $\Delta F/F_0$ traces (bottom) are shown. Dashed lines indicated time points at which representative images are shown. Scale bar, 20 µm.

### Supplementary Figure/Table 35

| | $HaloTag7_{669}$ (PDB 8SW8) |
|---|---|
| **Data collection** | |
| Space group | $P4_32_12$ |
| Cell dimensions | |
| *a*, *b*, *c* (Å) | 62.74, 62.74, 163.55 |
| α, β, γ (°) | 90, 90, 90 |
| Resolution (Å) | 58.65-1.90 (1.94-1.90) |
| *R*merge | 0.121(1.341) |
| *I* / σ*I* | 23.0 (4.0) |
| Completeness (%) | 99.6 (98.9) |
| Redundancy | 26.3 (27.1) |
| **Refinement** | |
| Resolution (Å) | 58.65-1.90 |
| No. reflections | 26365 |
| $R_{work}$ / $R_{free}$ | 0.193, 0.246 |
| No. atoms | |
| Protein | 4635 |
| Dye-HaloTag ligand | 96 |
| Chloride ion | 1 |
| Water | 173 |
| *B*-factors | |
| Protein | 23.4 |
| Dye-HaloTag ligand | 44.3 |
| Chloride ion | 19.2 |
| Water | 29.1 |
| R.m.s. deviations | |
| Bond lengths (Å) | 0.010 |
| Bond angles (°) | 1.662 |

*Structure was determined for one crystal.
**Values in parentheses are for highest-resolution shell.

### Sequences of WHaloCaMPs

**WHaloCaMP1a**

MAEIGTGFPFDPHYVEVLGERMHYVDVGPRDGTPVLFLHGNPTSSYVWRNIIPHVAPTHRCIAPDLIGMGK
SDKPDLGYFFDDHVRFMDAFIEALGLEEVVLVIHDWGSALGFHWAKRNPERVKGIAFMEFIRPIPTWDEWP
EFARETFQAFRTTDVGRKLIIDQNVFIEWTLPMAVARARRKWQKTGHAVRAIGRLSSGGSGGSGGSGGSDQ
LTEEQIAEFKEAFSLFDKDGDGTITTKELGTVMRSLGQNPTEAELQDMINEVDADGNGTIDFPEFLTMMARK
MKDTDSEEEIREAFRVFDKDGNGYISAAELRHVMTNLGEKLTDEEVDEMIREADIDGDGQVNYEEFVQMM
TAKYLTEVEMDHYREPFLNPVDREPLWRFPNELPIAGEPANIVALVEEYMDWLHQSPVPKLLFWGTPGVLIP
PAEAARLAKSLPNCKAVDIGPGLNLLQEDNPDLIGSEIARWLSTLEISG

HaloTag, MLCK-peptide, Linker, Calmodulin, HaloTag
Rationally placed tryptophan: G171W
Hits from directed evolution screen: G176A, V178A, P180Y

**WHaloCaMP with ENOSP peptide**

MAEIGTGFPFDPHYVEVLGERMHYVDVGPRDGTPVLFLHGNPTSSYVWRNIIPHVAPTHRCIAPDLIGMGK
SDKPDLGYFFDDHVRFMDAFIEALGLEEVVLVIHDWGSALGFHWAKRNPERVKGIAFMEFIRPIPTWDEWP
EFARETFQAFRTTDVGRKLIIDQNVFIEWTLPMAVIRRKKTFKEVANAVKISASLMGGGSGGSGGSGGSDQL
TEEQIAEFKEAFSLFDKDGDGTITTKELGTVMRSLGQNPTEAELQDMINEVDADGNGTIDFPEFLTMMARK
MKDTDSEEEIREAFRVFDKDGNGYISAAELRHVMTNLGEKLTDEEVDEMIREADIDGDGQVNYEEFVQMM
TAKTLTEVEMDHYREPFLNPVDREPLWRFPNELPIAGEPANIVALVEEYMDWLHQSPVPKLLFWGTPGVLIP
PAEAARLAKSLPNCKAVDIGPGLNLLQEDNPDLIGSEIARWLSTLEISG

HaloTag, ENOSP-peptide, Linker, Calmodulin, HaloTag
Rationally placed tryptophan: G171W
Hits from directed evolution screen: G176A, V178I, P180T

**WHaloCaMP1b**

MAEIGTGFPFDPHYVEVLGERMHYVDVGPRDGTPVLFLHGNPTSSYVWRNIIPHVAPTHRCIAPDLIGMGK
SDKPDLGYFFDDHVRFMDAFIEALGLEEVVLVIHDWGSALGFHWAKRNPERVKGIAFMEFIRPIPTWDEWP
EFARETFQWFRTARRKWQKTGHAVRAIGRLSSGGSGGSGGSGGSDQLTEEQIAEFKEAFSLFDKDGDGTITT
KELGTVMRSLGQNPTEAELQDMINEVDADGNGTIDFPEFLTMMARKMKDTDSEEEIREAFRVFDKDGNGYI
SAAELRHVMTNLGEKLTDEEVDEMIREADIDGDGQVNYEEFVQMMTAKVDRKLIIDQNVFIEGTLPMGVVR
PLTEVEMDHYREPFLNPVDREPLWRFPNELPIAGEPANIVALVEEYMDWLHQSPVPKLLFWGTPGVLIPPAE
AARLAKSLPNCKAVDIGPGLNLLQEDNPDLIGSEIARWLSTLEISG

HaloTag, MLCK-peptide, Linker, Calmodulin, HaloTag
Rationally placed tryptophan: A151W
Hits from directed evolution screen: G158D
