## Supplementary material for "A modular chemigenetic calcium indicator enables in vivo functional imaging with near-infrared light": Methods

**Molecular biology**

Synthetic DNA oligonucleotides were purchased from Twist Bioscience and Integrated DNA technologies. Q5 high fidelity DNA polymerase (New England Biolabs) was used for all PCR amplifications. Isothermal assembly reactions were performed with a NEBuilder HiFi kit (New England Biolabs). Small scale DNA isolation was performed with QIAprep Spin Miniprep Kit (Qiagen). Vector backbones were acquired from the following sources: pRSET was acquired from Life Technologies; pGP-AAV-syn-jGCaMP7f-WPRE was a gift from Douglas Kim & GENIE Project (Addgene plasmid # 104488); pAAV-CaMKIIa-EGFP was a gift from Bryan Roth (Addgene plasmid # 50469); p10xUAS-IVS-Syn21-Voltron-p10 was from a previously deposited Addgene plasmid (Addgene plasmid # 119041); pTol2-HuC(*elavl3*)-CaMPARI2 was from a previously deposited Addgene plasmid (Addgene plasmid # 137185); piGECI-N1 was a gift from Vladislav Verkhusha (Addgene plasmid # 160421) ;pBAD-iGECInano was a gift from Vladislav Verkhusha (Addgene plasmid # 186189). pCAGGS was used from an inhouse source. Cloning was conducted by PCR amplification (pDUET, pRSET) or restriction enzyme digestion of vector backbones. For use in mammalian cell culture, inserts were cloned into pCAGGS at the AflII and AgeI restriction sites. For use in rodent neurons, the inserts were cloned into pAAV-hsyn or pAAV-CaMKII at the BamHI and HindIII restriction sites. For expression in *Danio rerio* pTol2 with HuC *elavl3* promoter, the inserts were cloned into the AgeI restriction site. For expression in *Drosophila melanogaster*, the insert was cloned into 10XUAS-IVS-Syn21 at the XhoI and XbaI restriction sites.

Inserts were amplified by PCR amplification. WHaloCaMP constructs were cloned with a nuclear exclusion signal (NES) to mimic the cellular localization of jGCaMPs in neurons. Vector backbones and inserts were assembled by isothermal assembly with 10-30 base pair overlap, and sequence verified by Sanger sequencing (Azenta Life Sciences) or by nanopore full-plasmid sequencing (Plasmidsaurus). For large scale plasmid preparation, DNA was isolated by the Janelia Molecular Biology Facility. Adeno-associated viruses (AAVs) were prepared by Janelia Virus Services. Plasmids generated in this work have been deposited with Addgene: pRSET-WHaloCaMP1a-EGFP (205303), pRSET-WHaloCaMP1a (205304), pRSET-WHaloCaMP1b-EGFP (205305), pRSET-WHaloCaMP-eNOSpep-EGFP (205306), pAAV-synapsin-WHaloCaMP1a-EGFP (205307), pAAV-synapsin-WHaloCaMP1a (205308), pAAV-CaMKII-WHaloCaMP1a-EGFP (205309), pAAV-CaMKII-WHaloCaMP1a (205310), pTol2-elavl3-WHaloCaMP1a-EGFP (205311), pTol2-elavl3-WHaloCaMP1a (205312).

**Protein expression and purification**

For expression and purification of proteins, T7 express (New England Biolabs) were transformed with pRSET plasmids encoding for the protein of interest. The bacteria were grown in auto-induction media using the Studier method[1] with antibiotics at 30 °C for 48 h shaking at 200 rotations per minute (r.p.m.). Cell pellets were collected by centrifugation, lysed in TRIS Buffered Saline (TBS) (19.98 mM Tris, 136 mM NaCl, pH 8.0), with n-Octyl-β-D-thioglucopyranoside (5 g $L^{-1}$). Aggregations were disrupted by sonication and the lysate cleared by centrifugation. Protein purification was performed on a N-terminal poly-histidine ($His_6$) tag using HisPur Ni-NTA resin (Thermofisher Scientific), according to manufacturer's recommendations. Purified proteins were buffer exchanged into TBS using Amicon concentration filters (Merck). Protein aliquots were stored at 4 °C, or flash frozen and stored at -80 °C until assayed.

**Polyacrylamide gel electrophoresis**

To verify protein expression and covalent bond formation with dyes-HaloTag Ligand, proteins were run sodium dodecyl sulfate–polyacrylamide gel electrophoresis (SDS-PAGE). Proteins were pre-labeled with 1.2 molar equivalence of dye-ligands, before being separated on an NuPAGE 4 to 12%, Bis-Tris protein gel (ThermoFisher Scientific). Imaging of in gel fluorescence was performed using a ChemiDoc MP (BioRad) with epi-blue (460-490 nm), epi-green (520-545 nm) or epi-red (625-650 nm) excitation, and emission filters set to 518-546 nm, 577-613 nm, or 675-725 nm. PageBlue protein staining solution (ThermoFisher Scientific) was used to visualize the total protein amount in the gel.

**Crystallography**

For crystallization, HaloTag7 was further purified by size exclusion chromatography Superdex 200 10/300 GL column (GE Healthcare) at a flow rate of 0.5 ml $min^{-1}$ in 50 mM Tris-HCl, 75 mM NaCl, pH 7.4. The purified protein was incubated with 2 molar equivalants of $JF_{669}$-HaloTag ligand for 5 hours at 22 °C (room temperature), and excess dye-ligand was removed by PD MiniTrap desalting columns with Sephadex G-25 resin (Cytiva Life Sciences), pre-equilibrated 50 mM TRIS-HCl, with 75 mM NaCl, pH 7.4. Crystallization was performed at 22 °C (room temperature) and the crystals were grown using the sitting drop vapor diffusion method in 96-well plates using precipitant solutions from Crystal Screen Cryo HT (Hampton Research). $HaloTag7_{669}$ (10 mg $mL^{-1}$) was crystallized by mixing 1:1 vol:vol with a precipitant solution of 0.2 M ammonium phosphate monobasic, 0.1 M TRIS pH 8.5, 50% v/v (+/-)-2-methyl-2,4-pentanediol. Blue $HaloTag7_{669}$ crystals were plunged into liquid nitrogen for storage and transport. X-ray diffraction data were collected at beamline 8.2.2 at the Advanced Light Source, and a wavelength of 1 Å and a temperature of 100 K under a cold nitrogen stream. Diffraction data were integrated using Xia2[2]/DIALS[3] and scaled using Scala from within the CCP4 suite[4]. The structure was solved using PHASER[5], using a prior structure of HaloTag7 bound to trimethyl rhodamine HaloTag7 Ligand as a search module (PDBID 6U32). Refinement was performed iteratively using REFMAC5[6], and model rebuilding/adjustments in Coot[7]. The refined structure of $HaloTag7_{669}$ (PDBID 8SW8) exhibited good model geometry with 94.9% of residues in the favored regions of the Ramchandran plot and 1% in disallowed regions.

**Library construction**

For directed evolution of WHaloCaMPs, the gene of interest fused to EGFP as an expression marker were cloned in the desired geometry into the pRSET plasmid. NNS degenerate codons were used for single site mutagenesis and primers designed using the 22-codon trick[8] were used for two site mutagenesis. PCR was used to amplify the gene fragment including the degenerate codon site (s) and NEBuilder HiFi kit (New England Biolabs) according to the manufacturer's instructions. The reactions were diluted 1:3 before electroporated into T7 express *E. coli* and plated on LB + ampicillin plates to isolate single colonies. Around 10-20 colonies were sequenced from each reaction to verify library generation.

**Library bacterial lysate screen**

For library expression, single colonies were picked and transferred to liquid growth media in 96-well blocks containing 800 µL autoinduction media and ampicillin (100 µg $mL^{-1}$). The blocks were shaken at 400 r.p.m. for 48 hours. The bacteria were then pelleted, and lysed by 3 freeze/thaw cycles at -20 °C or -80 °C. The pellet was resuspended in 30 mM 3-morpholinopropane-1-sulfonic acid (MOPS), 100 mM KCl at pH 7.2, with shaking for ~1h at 37 °C, and the lysate cleared by centrifugation. After clearing, 250 µl of the lysate was transferred into 96-well blocks of 10 µL pre-aliquoted solutions of 10 µM $JF_{669}$-HaloTag ligand in 30 mM MOPS, 100 mM KCl at pH 7.2 (final labeling concentration of $JF_{669}$-HaloTag ligand was ~380 nM). Two 96-well black plates were prepared with "low" concentration of $Ca^{2+}$ (5 µL, 10 mM $CaCl_2$ in 30 mM MOPS, 100 mM KCl at pH 7.2) or "low" concentration of ethylene glycol-bis(β-aminoethyl ether)-N,N,N′,N′-tetraacetic acid (EGTA ) (5 µL, 20 mM EGTA in 30 mM MOPS, 100 mM KCl at pH 7.2). 95 µL of the $JF_{669}$-dye labeled lysate was transferred into the prepared 96-well plates (final concentration low $Ca^{2+}$ is 0.5 mM, and low EGTA 1 mM) and fluorescence was read on a plate reader (TECAN Spark 20M) with a stacking module. Fluorescence was read at 485 nm excitation and 510 nm emission for EGFP and 670 nm excitation and 690 nm emission for proteins labeled with $JF_{669}$ with 5 nm bandgap. After initial reading the $Ca^{2+}$ and EGTA conditions were swapped: 20 µL of a solution with 100 mM EGTA was added to the 96-well plate containing low $Ca^{2+}$, and 20 µL of a solution with 50 mM $CaCl_2$ was added to the 96-well plate containing low EGTA. The fluorescence emission was again read on a plate reader (TECAN Spark 20M) with a stacking module. To identify top performing variants, variants above a threshold value of EGFP fluorescence were ranked from calculating $\Delta F/F_0$ (where $\Delta F$ = fluorescence intensity in $Ca^{2+}$- fluorescence intensity in EGTA, and $F_0$ is fluorescence intensity in EGTA) of the $JF_{669}$ signal both in the "low" $Ca^{2+}$/EGTA and "high"

$Ca^{2+}$/EGTA regime. Depending on the library, 0-38 top hits were chosen, expanded from the pelleted lysate before labeling and sequenced through sanger sequencing.

**Library $Ca^{2+}$-response validation**

Unique sequences of top performing variants were expressed and purified using their N-terminal $His_6$-tag as described above. The concentration of the purified proteins was quantified on the absorbance of EGFP fused to the C-terminal ($\varepsilon_{488}$ = ~55 000 $cm^{-1}$ $M^{-1}$). The variants were labeled with sub stoichiometric amounts of dye-ligand (15 µM purified protein with 10 µM $JF_{669}$-HaloTag ligand) for at least 1 hour at 22 °C (room temperature). The labeled variants (2 µL) were aliquoted into 98 µL of 0.5 mM "low" $Ca^{2+}$ and 1 mM "low" EGTA and the fluorescence emission read on a plate reader, recording both the EGFP and $JF_{669}$ fluorescence emission, as previously done. The "low" conditions were then changed to "high" by adding 20 µL of 100 mM EGTA to the "low" $Ca^{2+}$ wells and 20 µL of 50 mM $CaCl_2$ to the "low" EGTA wells and fluorescence read once again on the plate reader. The $\Delta F/F_0$ was calculated for each variant, both in the "low" $Ca^{2+}$/EGTA and "high" $Ca^{2+}$/EGTA regime. The overall fluorescent intensity was compared to HaloTag-EGFP labeled and read under the same conditions in the plate reader. The variants with the largest $\Delta F/F_0$ in the validation were further validated in a dye-ligand capture assay by fluorescent polarization.

**Fluorescence polarization dye-ligand capture rate assay**

To determine the dye-ligand capture rate of the protein variants with dye-HaloTag ligand a fluorescence polarization assay was used. Purified protein was diluted into buffer containing 30mM MOPS, 100 mM KCl, pH 7.2, 0.5 mg $mL^{-1}$ bovine serum albumin (BSA), with either 10 mM EGTA or 20 mM $CaCl_2$. Proteins were prepared to 625 nM, 312.5 nM, 156 nM and 78 nM. 80 µL of the diluted solutions were added to 96-well black well plates. A solution of $JF_{549}$-HaloTag ligand was prepared in 30mM MOPS, 100 mM KCl, pH 7.2, 0.5 mg $mL^{-1}$ BSA, 10 mM EGTA to a final concentration of 62.5 nM. Fluorescence polarization was read over 5-20 minutes on a plate reader (Tecan Spark 20 M) at 37 °C, after mixing 80 µl of the protein solutions with 20 µl of the dye-ligand solution with a final concentration of protein at 500 nM, 250 nM, 125 nM and 62.5 nM, and the final concentration of dye-ligand at 12.5 nM. Excitation was set to 535 nm and emission at 580 nm, with 20 nm bandwidth. The G-factor was manually set to 1.79 after measuring a dilute concentration of dye-ligand in buffer with the same excitation and emission settings. The binding curve over time was fit with a single exponential in a custom written python script with curve_fit in scipy.optimize were (y) is the fluorescence polarization in mP, $Y_0$ is the value of Y when intercepting the y-axis, Pl is the value of Y at plateau, k is the pseudo-first order rate constant, and x is the time in s.

$$y(x) = Y_0 + (Pl - Y_0) \times (1 - e^{-kx})$$

The pseudo first order rate constant was calculated for each protein concentration and an apparent or calculated second order rate constant for dye-ligand capture was calculated, where $k_{2,calc}$ is the calculated second order rate constant in $M^{-1}$ $s^{-1}$, k is the pseudo-first order rate constant and [protein] is the protein concentration during the run.

$$k_{2,calc} = \frac{k}{[protein]}$$

This plate reader assay was too slow to capture rates faster than $10^6$ $M^{-1}$ $s^{-1}$, however, we used it to verify that we had not lost the fast dye-ligand capture rate for the variants.

**Ultraviolet-visible spectroscopy of protein-dye conjugates**

To record absorbance spectra of and molar extinction coefficients of WHaloCaMP bound to dyes, proteins were first pre-labeled with dye-HaloTag ligand. The dye-HaloTag ligand was reconstituted from lyophilized powder to 2 mM in DMSO. For excess dye-ligand addition and subsequent removal, HaloTag, or variants of WHaloCaMP, were diluted to 50 µM in TBS and dye-HaloTag ligand was added to a final concentration of 100 µM. After incubation of at least 1 hour at 22 °C (room temperature), excess dye-ligand was removed by a PD SpinTrap G-25 (Cytiva Life Sciences), pre-equilibrated with 30 mM MOPS, 100 mM KCl at pH 7.2. After elution the protein-dye concentration was quantified by NanoDrop Onec UV-VIS spectrophotometer (Thermo) by reading the absorbance at 280 nm and using a calculated molar extinction coefficient at 280 nm

(Expasy, ProtParam tool). Protein concentrations were around 40 µM. For sub stoichimetric labeling of the protein variants, 20 µM protein was diluted in TBS and labeled with 10 µM JFdye-HaloTag ligand. DMSO concentrations did not exceed 1% vol/vol. The protein-dye conjugates were diluted to 1.5 µM the two solutions from Calcium Calibration Buffer Kit #1 (Invitrogen) zero and 10 mM CaEGTA, containing either 10 mM EGTA or 39 µM free [$Ca^{2+}$] in 30 mM MOPS, 100 mM KCl at pH 7.2. Absorption spectra were recorded on a Cary Model 100 spectrometer (Agilent) using 1-cm path length semi-micro quartz cuvettes with self-masking sides (1 mL volume). Spectra were normalized to the max absorption in the presence of saturating $Ca^{2+}$. For molar extinction measurements, proteins-dye conjugates were further diluted in with the zero or 10 mM CaEGTA buffers to 750 nM, 375 nM and 187 nM and the absorbance spectra recorded. The molar extinction coefficient at a particular wavelength was determined by performing a background correction and plotting the absorbance (y) against the concentration (x). The molar excitation coefficient (b) was taken to be the line of the line fit through the points.

$$y\ (x) = a + bx$$

**Fluorescence spectroscopy of protein-dye conjugates**

To record fluorescence excitation and emission spectra proteins were first pre-labeled with dye-HaloTag ligand in a sub stoichiometric molar ratio. 20 µM protein was diluted in TBS and labeled with 10 µM JFdye-HaloTag ligand. DMSO concentrations did not exceed 1% vol/vol. After incubation of at least 1 hour at 22 °C (room temperature), the protein-dye conjugate was diluted in either zero or 10 mM CaEGTA buffers. The final concentration of the protein-dye conjugate was 666 nM. Fluorescence spectra were recorded on a Cary Eclipse fluorometer (Agilent) using 1 cm path length quartz spectrophotometer 3.5-mL cuvettes. Spectra were normalized to the max fluorescence excitation or emission in the presence of saturating [$Ca^{2+}$].

**Two-photon spectroscopy of protein-dye conjugates**

WHaloCaMP1a was prelabeled with a sub stoichiometric amount of dye (10 µM dye-ligand to 20 µm WHaloCaMP1a) for at least 1h at room temperature. The protein-JF-dye ligand conjugates were then diluted to 1 μM final concentration of dye-ligand equivalence in either zero or 10 mM CaEGTA buffers from Calcium Calibration Buffer Kit #1 (Invitrogen). Two-photon spectroscopy of WHaloCaMP-JF dye ligand conjugates were performed as previously described[9, 10]. Briefly, an inverted microscope (IX81, Olympus) equipped with a 60x1.2 numerical aperture (NA) water objective (Olympus) was used. To excite the samples, a pulsed 80 MHz Ti-Saphire laser (Chameleon Ultra II, Coherent) was used for 710-1080 nm and OPO (Chameleon Compact OPO, Coherent) for 1000-1500 nm. For excitation of $WHaloCaMP1a_{494}$ and $WHaloCaMP1a_{552}$, a dichroic filter (675DCSXR, Omega) and a short pass filter (720SP, Semrock) were used, and emission was collected using a band pass filter (539BP278, Semrock). Whereas for $WHaloCaMP1a_{669}$ and $WHaloCaMP1a_{722}$, laser excitation was achieved through a dichroic filter (FF825-SDio1, Semrock) and emission was collected using a band pass filter (709BP167, Semrock). Fluorescence detection was achieved by a fiber-coupled Avalanche Photodiode (SPCM_AQRH-14, PerkinElmer). All the fluorescence excitation spectra are corrected for the wavelength-dependent transmission of the dichroic and bandpass filters, and quantum efficiency of the detector. The corrected spectra were used to calculate the action cross section using fluorescein[11], Rhodamine B[12], and styryl 9M[12] as a reference. Spectra are averages (n = 2).

**Quantum yield determination of protein-dye conjugates**

To record quantum yields proteins were first pre-labeled with JFdye-HaloTag ligand in a sub stoichiometric molar ratio. The protein-dye ligand conjugates were diluted in either zero or 10 mM CaEGTA buffers. A Quantaurus-QY spectrometer (model C11374, Hamamatsu) with an integrating sphere was used and measurements were performed on dilute samples (A < 0.1) and self-absorption corrections were performed[13] using the instrument software.

**Calcium titrations**

To determine the calcium affinity and cooperativity of the WHaloCaMP sensors, calcium titrations were performed with a commercial Calcium Calibration Buffer Kit #1 (Invitrogen) with zero and 10 mM CaEGTA. Proteins were pre-labeled with excess dye-HaloTag ligand and any excess dye-ligand removed by PD SpinTrap G-25 (Cytiva Life Sciences), pre-equilibrated with 30 mM MOPS, 100 mM KCl at pH 7.2. 2 µL of the pre-labeled protein-dye conjugate was diluted into 98 µL of a pre-mixed solutions of Ca-ETA in black 96-well plates. Fluorescence intensities were read on a plate reader (Tecan Spark 20M). For WHaloCaMP1a$_{494}$ excitation was 490 nm and emission 530 nm. For WHaloCaMP1a$_{552}$ excitation was 560 nm and emission 580 nm. For WHaloCaMP1a$_{669}$ excitation was 670 nm and emission 690 nm. For WHaloCaMP1a$_{722}$ excitation was 730 nm and emission 750 nm. All bandwidths were set to 5 nm. The free calcium concentration was calculated taking the dissociation constant of EGTA for $Ca^{2+}$ to be 150 nM at 22 °C and pH 7.2[14]. The fluorescence (y) was plotted against the free calcium concentration (x) and a four-parameter dose-response curve (variable slope) using GraphPad Prism software was fit where (a) is the value of fluorescence at the bottom of the curve, (b) is the value of fluorescence at the top of the curve, ($EC_{50}$) is the concentration of agonist that gives a response halfway between bottom and the top, and (h) is the hill or cooperative coefficient.

$$y = a + \frac{x^h \times (b - a)}{x^h + {EC_{50}}^h}$$

**pH sensitivity of probes in vitro**

Proteins were pre-labeled with dye-HaloTag ligand in a sub stoichiometric molar ratio: 20 µM protein was diluted in TBS and labeled with 10 µM dye-HaloTag ligand. Labeled protein was then diluted into the following buffer systems containing 1 mM $CaCl_2$ or 50 mM EGTA of the following buffer systems: citrate (pH 4.0–6.2); phosphate (pH 5.8–8.0); tris (pH 7.8–9.0), all with 10 mM buffer and 150 mM NaCl. 2 µL of the protein was diluted into the buffer in 96-well black plates and fluorescence was read on a plate reader (Tecan Spark 20M).

**Stopped flow kinetic measurements of $Ca^{2+}$ kinetics**

$Ca^{2+}$ unbinding kinetics of WHaloCaMPs were determined using a Photophysics SX-20 Stopped-flow device. The pre-labeled protein (~1.3 µM) was rapidly mixed with a solution of 30 mM MOPS, 100 mM KCl, 10 mM EGTA, pH 7.2 in a 1:1 ratio. The sensors were excited using a lightemitting diode (625 nm), and fluorescence emission was collected through a 665-nm longpass filter. Fluorescence decay was fit with decay curves described in the supp figs.

**One-photon widefield bleaching of WHaloCaMP1a$_{669}$**

One-photon bleaching experiments were performed on inverted Nikon Eclipse Ti2 microscope as previously described[15]. Briefly, bleaching experiments were carried out in aqueous droplets of purified protein labeled with dye-ligand isolated in 1-octanol[16]. Labeled purified protein were diluted to 500 nM in the commercial Calcium Calibration Buffer Kit #1 (Invitrogen) with zero and 10 mM CaEGTA and supplemented to 0.3 mg mL$^{-1}$ BSA. This solution was diluted 10-fold in a 1-octanol/PBS suspension and agitated by tapping. The solution was placed on a pre-silanized glass slide and sandwiched with a coverslip. The microdroplets were illuminated using an 40× (NA = 1.3, PLAN Flour, Nikon (FN = 25.2)) oil immersion objective. Illumination was performed with an LED (SpectraX Light engine, Lumencor), and imaged through the following filter cube 89000 cube (Chroma), ET645/30x, 89100bs, ET705/72m. Images were captured by a scientific CMOS camera (ORCA-Flash 4.0, Hamamatsu), and images were acquired with 1 s exposure every 2 s. Power at the objective was measured with a microscope slide power sensor (S170C, Thorlabs) to 7 mW, giving an irradiance of 23 mW mm$^{-2}$. Each sample was bleached continuously for 6.6 minutes. Background subtraction was performed in FIJI[17], the fluorescence emission was normalized to the first frame. One phase decay was fit to the data in the GraphPad Prism software.

**Primary neuronal culture and labeling with dyes-HaloTag ligand**

Procedures involving rodent animals were conducted in accordance with protocols approved by the Howard Hughes Medical Institute (HHMI) Janelia Research Campus Institutional Animal Care and Use Committee and Institutional Biosafety Committee. Hippocampal neurons extracted from P0 to 1 Sprague-Dawley rat pups and plated into 24-well glass bottom plates or 35 mm dishes (Mattek, #1.5 coverslip). A coating of poly-D-lysine was applied prior to plating the cells. Neurons were cultured at 37 °C with 5% $CO_2$ in a humidified atmosphere in NbActiv4 medium (BrainBits). 3-4 days after plating, neurons were transduced with AAV1 virus to express the gene of interest under a *hsyn1* promoter. 17-21 days after transduction, the neurons were labeled with dye-HaloTag ligand. Dye-HaloTag ligands were first diluted to 2 mM stock solutions in DMSO from lyophilized powder. The neurons were labeled with 50-100 nM dye-HaloTag ligand at 37 °C for 30 min-45 min. The neurons were then washed with imaging buffer containing 145 mM NaCl, 2.5 mM KCl, 10 mM glucose, 10 mM HEPES, pH 7.4, 2 mM $CaCl_2$ and 1 mM $MgCl_2$. After 3 washes, imaging buffer containing synaptic blockers[18] (10 µM 6-cyano-7-nitroquinoxaline-2,3-dione (CNQX), 10 µM 3-(2-carboxypiperazin-4-yl)propyl-1-phosphonic acid (CPP), 10 µM Gabazine and 1 mM (S)-α-methyl-4-carboxyphenylglycine (MCPG)) were added to block ionotropic glutamate, gamma aminobutyric acid (GABA), and metabotropic glutamate receptors.

**Field stimulation of WHaloCaMP1a in primary neuronal culture.**

Field stimulation of neurons were performed using platinum wire electrode inserted in the medium, controlled by a high-current isolator (A385, World Precision Instruments) set at 90 mA. Trains of action potentials were delivered to the platinum wire electrode from 1 to 160 APs, at intervals described in each imaging modality. Unless otherwise stated, the fluorescent trace from hand segmented neurons from at least 3 fields of view from at least two separate neuron batches were extracted in FIJI[17]. $\Delta F/F_0$ was calculated with $F_0$ being the average fluorescence recorded 1 s before each stimulus. Signal to noise ratio (SNR) was calculated from the amplitude in the fluorescent change on the field stimulation divided by the standard deviation of the baseline fluorescence 1 s before each stimulus. Calculated time to peak was calculated from the onset of stimulus to the maximal fluorescence value. The $t_{1/2}$ of decay was calculated by fitting a two-phase decay in Graphpad Prism software to the $\Delta F/F_0$ trace after the field stimulus has stopped.

**Widefield imaging in the visible spectrum**

Widefield imaging in the visible spectrum (up to 700 nm) was performed on an inverted Nikon Eclipse Ti2 microscope with a 20× air objective (NA = 0.75, Nikon) equipped with a Spectra X light engine (Lumencore) and imaged onto a ORCA-Flash 4.0 sCMOS camera (Hamamatsu). The SPECTRA X light engine was equipped with 485/25 nm, 550/15 nm, and 640/30 nm excitation filters for exciting $WHaloCaMP_{494}$, $WHaloCaMP_{552}$, and $WHaloCaMP_{669}$. A quad bandpass (set number: 89000, Chroma) with 490/20 nm, 555/25 nm, 645/30 excitation filter, a dichroic mirror (89100bs; Chroma) and 525/36 nm, 605/52 nm, and 705/72 nm emission filters (Chroma) were also used. Imaging was performed at 33 Hz, and the field stimulation was controlled by an Arduino Uno board synchronized with light sources in the Nikon Elements software. A train of 1, 2, 3, 5, 10, 20, 40, 80, 120 and 160 APs with 1 ms pulse duration at 80 Hz were elicited with 20 s time intervals between each stimulation.

**Widefield imaging in the near infrared**

For imaging WHaloCaMP1a$_{722}$ in the near infrared, we used a custom-built upright microscope as previously described[19] with modifications. Neurons expressing WHaloCaMP1a and labeled with JF$_{722}$-HaloTag ligand were imaged using a 40× objective (Nikon N40X-NIR, 0.8 NA, and 3.5 mm working distance). To excite WHaloCaMP1a$_{722}$ we used a 671 nm laser (gem 671, Laser Quantum). To image the EGFP expression marker, we used a 4 wavelength LED light source (Thorlabs LED4D067), using the LED at 470 nm. We used an imaging dichroic beam splitter with edge at 699 nm (FF699-FDI01-T1-25X36, Semrock) and 715 nm emission filter (FF01-715/LP-25, Semrock) to clean up fluorescence signal before detection by camera. Both visible and near-infrared fluorescence signals were imaged using an InGaAs camera with optimized sensitivity in the NIR/SWIR range (Ninox 640 II, Raptor Photonics). Images were acquired using µManager, an open source microscopy software[20]. Images were acquired at 10 frames per s (fps) for 400 to 500 frames for each run. We performed field stimulation with the stimulation controlled using the MATLAB-based waveform generator Wavesurfer to elicit 1, 5, 10, 20 or 160 APs with 1 ms pulse duration with a square pulse train at 80 Hz and analyzed the data as described under field stimulation.

**Simultaneous electrophysiology and fluorescence imaging in primary neuron culture**

All imaging and electrophysiology measurements were performed in imaging buffer. Internal solution for current clamp recordings contained the following: 130 mM potassium methanesulfonate, 10 mM HEPES, 5 mM NaCl, 1 mM $MgCl_2$, 1 mM Mg-ATP, 0.4 mM Na-GTP, 14 mM Tris-phosphocreatine, adjusted to pH 7.3 with KOH, and adjusted to 300 mOsm with sucrose. Glass capillary with filament (Sutter Instruments) were pulled to a tip resistance of 4 – 6 MΩ.

Pipettes were positioned with a MPC200 manipulator (Sutter Instruments). EPC800 amplifier (HEKA) were used for acquiring electrophysiology recordings, filtered at 10 kHz with the internal Bessel filter, and digitized using a National Instruments PCIe-6353 acquisition board at 20 kHz. Data were acquired from cells with access resistance < 25 MΩ. WaveSurfer software was used to generate the various analog and digital waveforms to control the amplifier, camera, light source, and record voltage and current traces. Cells were initially held at -70 mV. 440 nm light pulse (5 ms, 10 Hz at 0.45 mW $mm^{-2}$, when CheRiff is co-expressed with WHaloCaMP1a) or current injection (when only WHaloCaMP1a is expressed) were applied to evoke action potentials and initiate calcium entry into the cell.

640 nm light at 0.76 mW $mm^{-2}$ was used to excite WHaloCaMP1a with an emission filter at 705/72 nm. 555 nm light at 1.39 mW $mm^{-2}$ was used for excitation of jRCaMP1a and emitted fluorescence was filtered with a 607/70 nm emission filter. Fluorescence images were collected using a scientific CMOS camera (ORCA-Flash 4.0, Hamamatsu) and image acquisition was performed using HCImage Live (Hamamtsu) with frame rate at 30 Hz.

**Current clamp recording of CheRiff with different wavelength light**

For testing CheRiff's response to longer wavelength stimulation, 555 nm and 640 nm were applied for 1s to illuminate CheRiff-expressing neurons and voltage signals were monitored. Light powers were measured using a power meter (Thorlabs PM100A) with a Si photodiode (Thorlabs S120C or S170C).

**Acute mouse slices 2P imaging with WHaloCaMP1a**

All procedures for rodent husbandry and surgery were performed following protocols approved by the Washington University Institutional Animal Care and Use Committee and in accordance with National Institutes of Health guidelines. FLIM-AKAR, together with a transcriptional stop codon flanked by loxP sites, was knocked in at the ROSA26 locus. These mice were subsequently crossed with Emx1$^{IRES\ cre}$ mice (JAX #005628) to create mice homozygous for both Emx1-Cre and FLIM-AKAR at their respective loci.

WHaloCaMP1a was delivered via stereotaxic injections. AAV1-CAMKII-NES-WHaloCaMP1a ($5x10^{12}$ virus molecules $mL^{-1}$) was delivered using a UMP3 micro-syringe pump (World Precision Instruments) via glass pipette to P0/P1 pups. A total of 200 nL $pup^{-1}$ was delivered at 10 different coordinates (20 nL $coordinate^{-1}$), at a rate of 20 nL $s^{-1}$. The coordinates are as follows, relative to lambda in millimeters:

1. Anterior: 1.2; Lateral: 1.2; Depth from the skin: 1.2, 0.8, 0.4
2. Anterior: 0.8; Lateral: 1.5; Depth from the skin: 1.5, 1.0, 0.5
3. Anterior: 0.5; Lateral: 2.0; Depth from the skin: 2.0, 1.5, 1.0, 0.5

The $JF_{669}$-HaloTag ligand fluorescent dye was delivered either by retro-orbital (R.O.) injection into mice prior to acute slicing, or by incubation of acute brain slices (from dye-naive mice) in dye-infused ACSF. For R.O. injection, a total of 200 nmoles of $JF_{669}$-HaloTag ligand was first dissolved in 20 µL of DMSO, and subsequently diluted with 20 µL of 20% Pluronic in DMSO (Fisher) and 90 µL of PBS (Corning). Dye was injected within 24 hours of imaging. For acute slice incubation, 100 nmol of $JF_{669}$-HaloTag ligand were reconstituted in 20 uL of DMSO and mixed with ACSF to create a solution of 0.0626-0.25 µM. Acute brain slices were incubated in this dye-ACSF solution for 30 minutes, and were subsequently washed in dye-free ACSF for at least 30 minutes prior to imaging.

Mice were anesthetized with isoflurane and subsequently euthanized. All solutions were bubbled with carbogen (5% $CO_2$ and 95% $O_2$) prior to use and ACSF was continuously bubbled while in use. Mice aged P25 or younger were sectioned in cold sucrose cutting solution (containing in mM: 87 NaCl, 25 $NaHCO_3$, 1.25 $NaH_2PO_4$, 2.5 KCl, 75 sucrose, 25 glucose, 1 $MgCl_2$). Mice aged P30 or older underwent intracardiac perfusion with cold ACSF (containing in mM: 127 NaCl, 2.5 KCl, 25 $NaHCO_3$, 1.25 $NaH_2PO_4$, 2 CaCl2, 1 $MgCl_2$, and 25 glucose). Their brains were rapidly dissected and sectioned in cold choline cutting solution (containing in mM: 25 $NaHCO_3$, 1.25 $NaH_2PO_4$, 2.5 KCl, 7 $MgCl_2$, 25 glucose, 0.5 $CaCl_2$, 110 choline chloride, 11.6 ascorbic acid, 3.1 pyruvic acid). Acute coronal slices were collected at 300 µm thickness using a Leica VT1000S vibratome.

After sectioning, slices from both groups were transferred to ACSF for recovery at 34°C for 5-10 minutes. Following recovery, the slices were kept in ACSF at room temperature. Once ready for imaging, the slices were transferred to a microscope chamber where they were perfused with ACSF at a flow rate of 2- 4 mL $min^{-1}$.

A custom-built microscope was used to perform two-photon imaging in acute brain slices. $WHaloCaMP1a_{669}$ was excited at a wavelength of 1250 nm, and FLIM-AKAR at 920 nm. Two mode-locked laser sources were used in an alternating fashion to excite $WHaloCaMP1a_{669}$ (Spectra-Physics, Insight X3, 80 MHz) and FLIM-AKAR (Spectra-Physics, Mai Tai HP Ti:Sapphire, 80 MHz), respectively. Photons were collected with fast photomultiplier tubes (PMTs, Hamamatsu, H10770PB-40), using a 60X objective (Olympus, NA 1.1). Image acquisition was done with ScanImage, a custom-written software, in MATLAB 2012b[21].

FLIM was performed as described previously with FLIM-AKAR.[22, 23] The FLIM board used was SPC-150 (Becker and Hickl GmbH), and time-domain single photon counting was performed in 256 time channels. Intensity $Ca^{2+}$ imaging was performed with $WHaloCaMP1a_{669}$. Neurons with nuclear-excluded basal $WHaloCaMP1a_{669}$ signals were used for experiments. 128x128 pixel images were collected.

Both sensor's emission light was collected through a 580 dichroic mirror (FF580-FDi01-25X36, Semrock) followed by a 525/50 band-pass filter (FF03-525/50-25, Semrock) for FLIM-AKAR or a 690/50 band-pass filter (ET690/50m, Chroma) for $WHaloCaMP1a_{669}$.

All acute slice experiments were performed in the presence of 1 µM DPCPX to inhibit adenosine receptors, and 1 µM tetrodotoxin to block action potentials.

All chemicals were added to acute slices via addition into the perfusion reservoir. The final concentrations of chemicals are as follows: (+)-muscarine iodide (mus), 10 µM, from Tocris; tetrodotoxin citrate (TTX), 1 µM, from Hello Bio; DPCPX, 1 µM, from Tocris; forskolin (FSK), 50 µM, from Cayman Chemical; potassium chloride (KCl), 55 mM, from Cayman Chemical.

The region of interest (ROI) for each neuronal cell body was manually selected. An ROI for background fluorescence with no cellular processes was similarly selected. The fluorescence signal for all pixels in a neuron's ROI was averaged and the average background fluorescence was subtracted from it. The resulting net fluorescence was plotted against time. The $\Delta F/F_0$ was calculated as $(F-F_0)/F_0$, where $F_0$ is the fluorescence signal averaged over the entire baseline period.

The amplitudes of intensity changes were quantified as follows:

BaselineStart = average $\Delta F/F_0$ of first three intensity acquisitions of baseline;
BaselineEnd = average $\Delta F/F_0$ of last three intensity acquisitions of baseline;
MusEnd = average $\Delta F/F_0$ of last three intensity acquisitions of muscarine only application;
BaselineMax = max $\Delta F/F_0$ recorded during baseline;
MusMax = max $\Delta F/F_0$ recorded during muscarine only application;
FSKMax = max $\Delta F/F_0$ recorded during FSK application;

MaxΔBaseline = MaxBaseline – BaselineStart;
MaxΔMus = MusMax – BaselineEnd;
MaxΔFSK = FSKMax – MusEnd;

For presentation purposes, the WHaloCaMP1a$_{669}$ line was smoothened and presented along the raw data. Smoothing was performed in GraphPad Prism, using 6 neighboring points and a 2$^{nd}$ order polynomial for smoothing.

FLIM curve fitting, calculation of average lifetime over an ROI, and ROI analysis was performed as described previously.[22] The amplitudes of cytoplasmic lifetime changes were quantified as follows:

BaselineStart = average lifetime of the first three acquisitions of baseline;

BaselineEnd = average lifetime of the last three acquisitions of baseline;
BaselineMin = minimum lifetime during baseline;
MusEnd = average lifetime of the last three acquisitions of muscarine only application;
MusMin = minimum lifetime during muscarine only application;
FSKMin = minimum lifetime during forskolin application;

MaxΔBaseline = BaselineMin – BaselineStart;
MaxΔMus = MusMin – BaselineEnd;
MaxΔFSK = FSKMin – MusEnd;

The MATLAB programs for ScanImage for data acquisition and analysis are available at https://github.com/YaoChenLabWashU/2pFLIM_acquisition. All other code is available upon request.

**WHaloCaMP imaging in living adult flies using wide-field epifluorescence microscopy**

Experiments were performed on 2- to 10-day-old heterozygous progeny of a cross between a mushroom body driver (R13F02-Gal4) and the two WHaloCaMP variants, 10XUAS-NES::WHaloCaMP1a::EGFP-P10 in VK00005 and 10XUAS-NES::WHaloCaMP1b::EGFP-P10 in VK00005. jRGECO1a (20XUAS-IVS-NES-jRGECO1a-p10 in VK00005), which shares excitation and emission spectra with $JF_{552}$, was used as a comparison.

Dissections of head-fixed flies and dye-ligand incubation were performed similarly as in Voltron experiments[15]. Briefly, a small window was opened in the head cuticle, and fat tissue and trachea that overlaid the mushroom body calyx region were removed. The exposed brain was bathed in a drop (~200 µL) of dye-containing saline (5 µM for $JF_{669}$-HaloTag ligand and 1 µM for $JF_{552}$-HaloTag ligand for 1 hr. Saline contains (in mM): NaCl, 103; KCl, 3; $CaCl_2$, 1.5; $MgCl_2$, 4; $NaHCO_3$, 26; N-tris(hydroxymethyl) methyl-2-amino- ethane-sulfonic acid, 5; $NaH_2PO_4$, 1; trehalose, 10; glucose, 10 (pH 7.3 when bubbled with 95% $O_2$ and 5% $CO_2$), 275 mOsm. The brain was then washed with fresh saline several times and maintained in the saline for 1 hr. Imaging was performed on a wide-field fluorescence microscope (SOM, Sutter Instruments) equipped with a 60×, NA 1.0, water-immersion objective (LUMPlanFl/IR; Olympus) and a sCMOS camera (Orca Flash 4.0 v3, Hamamatsu). Images of 256 x 256 pixels were acquired at 10 Hz with 4x4 binning through the Hamamatsu imaging software (HCImage Live). Illumination was provided from an optical beam combining system (LB-OBC-LLG, Sutter) equipped with a 660-nm LED (FG-OBC-660, Sutter) with an excitation filter (FF01-660/13-25-STR, Semrock) and a 561-nm LED (FG-OBC-561, Sutter) with an excitation filter (FF01-554/23-25-STR, Semrock). As the incidence angles were arranged at 18° in the four-beam combiner, the bandpass spectra of the excitation filters were shifted toward a shorter wavelength by 5-10 nm (SearchLight spectra viewer modeling). For WHaloCaMP$_{669}$, the 660-nm LED was used and intensity at the sample plane was ~0.64 mW mm$^{-2}$; emission was separated from excitation light using a dichroic mirror (FF677-Di01-25/36, Semrock) and an emission filter (FF01-719/60-25, Semrock). For WHaloCaMP$_{552}$ and jRGECO1a, the 561-nm LED was used and intensities at the sample plane were ~0.50 mW mm$^{-2}$ and ~0.17 mW mm$^{-2}$ respectively; emission was separated from excitation light using a dichroic mirror (FF562-Di03-25x36, Semrock) and an emission filter (FF01-590/36-25, Semrock).

Odors used were 3-octanol (218405-50G, Sigma-Aldrich), 4-methylcyclohexanol (153095-250ML, Sigma-Aldrich) and apple cider vinegar (Heinz). Odors were diluted using a two-step, air-dilution-based olfactometer[24] to a final concentration of 1:16, and delivered to flies at 200 mL min$^{-1}$ for 2 s.

Data were analyzed in MATLAB with a custom script. Images were registered to correct for X-Y movement. Regions of interest (ROIs corresponding to calyx were manually selected, and the mean intensity of the ROI was extracted. $F_0$ was calculated as the mean over the 5 s of the imaging session before odor onset. Mean odor responses were integrated over a time window from 0.5 s and 5.5 s after odor onset.

**Mice**

All experimental protocols Janelia Research Campus were conducted according to the National Institutes of Health guidelines for animal research and were approved by the Institutional Animal Care and Use Committee at the Janelia Research Campus, HHMI. All experimental protocols at University of Maryland Baltimore were conducted according to the National Institutes of Health guidelines for animal research and approved by the Institutional Animal Care and Use Committee at University of Maryland Baltimore.

**Retroorbital injection of JF-dye ligands to mice**

Retro orbital injection of dyes-HaloTag ligands were performed as previously described[25] with minor modifications. 200 nmol of the lyophilized dye-HaloTag ligand was suspended in 20 µL DMSO. After vortexing, 20 µL of 20% Pluronic in DMSO was added and the solution carefully pipetted up and down to not form bubbles. 80 µL phosphate buffered saline (PBS, 137 mM NaCl, 10 mM phosphate, 2.7 mM KCl; pH 7.4) was carefully added to the solution and before letting the solution sit for 10 minutes to let foam and bubbles settle. 100 µL of this solution was injected into one eye of an anesthetized mouse using a 0.5 ml 27 G syringe to the retro-orbital sinus.

**Two-photon imaging of WHaloCaMP1a$_{669}$ in mouse visual cortex**

Viral injections (300 nl, 400 µm below brain surface) were performed followed by implantation of a 4mm cranial window, as described previously[26] and retroorbital injection of dye-ligands were performed 24 hours before imaging.

Two-photon imaging of WHaloCaMP1a$_{669}$ in the mouse visual cortex using long wavelength light was performed on a customized adaptive optics two photon microscope based on the Thorlabs Bergamo II microscope, previously described[19]. Mice were anesthetized with isoflurane using a precision vaporizer, 3% (vol/vol) in oxygen for induction and 0.5–2% (vol/vol) during visual stimulation. The mouse was placed in a custom-built goniometic heated holder to restrict movement and maintain a 37 °C body temperature. Excitation was provided from a Coherent Discovery NX TPC. The objective was an Olympus XLUMPFLN objective, 20x, 1.0 NA with water immersion. Excitation was performed at 950 nm (25% of light at laser source) for EGFP and 1225 nm (100% of light as laser source) for WHaloCaMP1a$_{669}$. Power ranged from 2-30 mW for 950 nm and 5-15 mW for 1225 nm. Emitted light was separated from excitation light via a 735 nm longpass dichroic, then filtered through 525/50 bandpass dichroic (green PMT) and 690/60 nm bandpass dichroic (near-infrared PMT). Light was collected via two ThorLabs PMT2100s, and digitized via a laser-clocked National Instruments DAQ card controlled by ScanImage[21] 2020 (Vidrio Technologies). Imaging parameters were 512 x 512 pixels with 44.8 fps at 2×zoom, for a 200 x 200 µm field of view. Visual stimulation was displayed on a LCD screen placed 10 cm in front of the mouse's eye and moving bar stimuli generated in MATLAB using the Psychtoolbox[20] (with 8 s ON, 8 s OFF) in eight equally spaced directions spaced by equal time periods of mean luminance and were shown to the animal synchronized with the acquisition. Light from the visual stimulation screen was filtered so that it minimally impacted the red and green PMT channels. Functional images were analyzed in suite2p[21] and fluorescent traces calculated with $F_0$ as the lowest 0.5 percentile, calculating a rolling mean over 22 frames (0.5 s).

**Two-photon imaging of $WHaloCaMP1a_{552}$ in mouse visual cortex**

A 3.5-mm diameter craniotomy was first made over left V1 of mice. Then 40 nl of virus-containing solution (AAV2/1.CAMK2.NES-WHaloCaMP1a-EGFP, $9 \times 10^{12}$ infectious units per ml) was injected 0.5 mm below pia into left V1 at four injection sites at the intersection points of the two left–right lines at Bregma −3.4 mm and −4.0 mm, and two anterior–posterior lines at 2.2 mm and 2.6 mm from the midline over 2 minutes. After the pipette was pulled out of the brain, a glass window made of a single coverslip (Fisher Scientific, no. 1.5) was embedded in the craniotomy and sealed in place with dental acrylic. A titanium headpost was then attached to the skull with cyanoacrylate glue and dental acrylic.

Visual stimuli were presented by a tablet monitor (SCEPTRE) displaying only blue light. The screen was positioned 15 cm from the right eye, covering 70° × 70° of visual space and oriented at ∼40° to the long body axis of the animal. Visual stimuli were generated using custom-written codes based on Psychophysics Toolbox[27] in Matlab (Mathworks). During visual stimulation, the luminance level was kept constant. Each stimulus trial consisted of 8 s blank period (uniform gray display at mean luminance) followed by 8 s drifting grating (0.05 cycles per degree, 1 Hz temporal frequency, eight randomized different directions). Each oriented drifting grating was presented for a total of 5 trials. Imaging was synchronized with ScanImage to start 4 s after the onset of gray session for 8 s. Therefore, each imaging session covers 4 s of gray screen and 4 s of drifting grating period.

Mice were kept on a warm blanket (37˚C) and anesthetized using 0.5% isoflurane and sedated with chlorprothixene (20–30 µL at 0.33 mg $mL^{-1}$, intramuscular). Imaging was performed with a custom-built two-photon microscope with a resonant scanner. Imaging was performed 4 weeks after virus injection and 24-hours after R.O. injection of the dye-ligand. Each experimental session lasted 45 min to 2 h. Multiple sections (imaging planes) may be imaged within the same mouse. Fluorophores were excited by different wavelength with a femtosecond laser system (Chameleon Discovery, Coherent) that was focused by an Olympus 25×, 1.05 NA objective. Emitted fluorescence photons reflected off a dichroic long-pass beamsplitter (FF705-Di01–25 × 36; Semrock) and were split with a dichroic mirror (565DCXR, Chroma) detected by photomultiplier tubes (H16201P-40//004, Hamamatsu) after filtered by 510/84 filter (84-097, Edmund) for the green channel and two 750SP filters (64-332, Edmund) for the red channel. Images were acquired using ScanImage[21] (Vidrio Technologies). Functional images (512 × 512 pixels, 268 × 268 $\mu m^2$) of L2/3 cells (50–300 µm under the pia mater) were collected at 15 Hz. For 850 nm, post-objective laser power was 15 mW for imaging 148 to 210 um deep brain. For 1050 nm, post-objective laser between 92 and 140 mW was used depending on the depth (100 to 362 um) of cells.

Imaging data were processed with custom programs written in MATLAB (Mathworks) and Fiji[17]. Images were registered with an iterative cross-correlation-based registration algorithm[28]. All identifiable cell bodies were outlined by hand as regions of interest (ROIs). The averaged fluorescence signal within the ROI was calculated as the raw ROI signal Fraw. Neuropil subtraction was performed as follows. First, A square neuropil region, centered on an ROI, was determined by adding 7 pixels to the width and height of the ROI, while excluding a 2-pixel gap between the ROI and the neuropil region. Then, the average fluorescence signal of the neuropil excluding the ROI, was calculated as the raw neuropil signal $F_{neuropil}$. The neuropil signal was then baseline-subtracted by subtracting the average of the lowest 5% values of the neuropil signal time trace to get $\Delta F_{neuropil}$. Neuropil-subtracted fluorescence signal within the ROI was then calculated[29] as

$F = F_{raw} - 0.7*\Delta F_{neuropil}$. We calculated $\Delta F/F_0 = (F - F_0)/F_0$, where F is the instantaneous fluorescence signal and $F_0$ is the average fluorescence in the interval 4 s before the start of the visual stimulus. Visual responses were measured for each trial as $\Delta F/F_0$, averaged over the stimulus period. Visually responsive neurons were defined as cells with significant stimulus-related fluorescence changes (ANOVA across blank and eight direction periods, $P < 0.01$) with an average $\Delta F/F_0$ at preferred orientations greater than 10%. The fraction of cells detected as responsive was calculated as the number of significantly responsive cells over all the ROIs analyzed. The cumulative distribution of peak $\Delta F/F_0$ responses included the maximal response amplitude from all analyzed cells, calculated as described above for each cell's preferred stimulus. The orientation sensitivity index (OSI) was calculated as before[29, 30] by fitting the fluorescence response from individual cells to the eight drifting grating stimuli with two Gaussians, centered at the preferred response angle ($R_{pref}$) and the opposite angle ($R_{opp}$). The OSI was calculated as

$$OSI = \frac{R_{pref} - R_{orth}}{R_{pref} + R_{orth}}$$

where $R_{orth}$ is the orthogonal angle to the preferred angle.

Standard functions and custom-written scripts in MATLAB were used to perform all analysis. The data were tested for normal distribution. Parametric tests were used for normally distributed data and non-parametric tests were applied to all other data. Bar graphs and mean ± standard error of the mean (SEM) were used to describe the data with normal distribution, while box plots and median ± interquartile range (IQR) were used to describe the non-normally distributed data. Box plots represent median and 25th—75th percentiles and their whiskers shown in Tukey style (plus or minus 1.5 times IQR). A nonparametric test (Wilcoxon signed-rank test) was used to examine paired data. Direct non-paired comparisons between two groups were made using Wilcoxon rank-sum test for non-normally distributed data. The statistical significance was defined as $^*P < 0.05$, $^{**}P < 0.01$, $^{***}P < 0.001$, respectively. Experiments were not performed blind. Sample sizes were not predetermined by statistical methods, but were based on those commonly used in the field. Medians, IQR, means and SEM are reported throughout the text.

**Zebrafish larvae**

All zebrafish experiments were conducted in accordance with the animal research guidelines from the National Institutes of Health and were approved by the Institutional Animal Care and Use Committee and Institutional Biosafety Committee of Janelia Research Campus. Larvae were reared using standard protocols in 14:10 light-dark cycles at 28.5 °C.

To generate WHaloCaMP1a lines, NES-WHaloCaMP1a or NES-WHaloCaMP1a-EGFP were cloned into Tol2 vectors with the *elavl3* pan-neuronal promoter. The plasmid was injected into 1-2 cell embryos from *casper* background zebrafish with mRNA encoding for the Tol2 transposon. Potential founders were selected either by screening with $JF_{522}$-HaloTag ligand or based on the EGFP expression. Potential founders were then screened by outcrossing to *casper* and selecting progeny (F1 generation) that expressed WHaloCaMP1a in the central nervous system. Experiments were performed on 4 to 5 day post fertilization (d.p.f) larvae produced from crosses with *casper*, from in crosses or from crossing with other lines for multicolor experiments. For three color multiplexed functional imaging, we crossed Tg(*elavl3*:NES-WHaloCaMp1a) with Tg(*actb2*:iGlucoSnFR; *acta1a*: jRGECO1a)[31] for neuronal and astrocyte imaging, we crossed Tg(*elavl3*:NES-WHaloCaMp1a-EGFP) with Tg(*gfap*:jRGECO1b)[32]. For comparing $WHaloCaMP1a_{669}$ against jRGECO1b in the same neuronal population, we crossed Tg(*elavl3*:NES-WHaloCaMP1a-EGFP) with Tg(*elavl3*:jRGECO1b)[33] .

**Dye delivery to zebrafish larvae**

Dye-ligands were delivered to the central nervous system of zebrafish larvae by adding the dye - HaloTag ligand (4 µM final concentration, from a 2 mM stock in DMSO) to the system water for 2 h. The larvae were then washed 2 × 1 h in fresh system water before mounting for whole brain imaging.

**Whole brain light sheet imaging with SimView**

Whole brain imaging was performed on a SiMView microscope as previously described[34, 35]. Tg(*elavl3*:NES-WHaloCaMP1a-EGFP) zebrafish larvae 5.p.f. were labeled with $JF_{669}$-HaloTag ligand, as described above. The fish were screened for EGFP fluorescence, and EGFP positive fish were paralyzed in a drop of 1 mg $ml^{-1}$ alpha-bungarotoxin solution (Invitrogen) in system water for 30 s, before recovering in system water for 30 minutes. Zebrafish were placed in 2% agarose in system water and positioned in a custom-designed glass capillary (2 mm outer diameter, 20 mm length; Hilgenberg GmbH), which was mounted in an imaging chamber filled with system water. Fluorescence light sheet imaging of the $WHaloCaMP1a_{669}$ was performed with Nikon 16x/0.8 NA water-dipping objectives and a Hamamatsu Orca Fusion BT camera. A 685 nm laser (1.6 mW $cm^{-2}$ at the sample) was used to excite $WHaloCaMP1a_{669}$ and the light was collected though a 795/188 nm detection filter (Semrock). Volumetric imaging was performed with an exposure time of 5 ms, at a step size of 4 µm across a 160 µm deep volume, resulting in volume imaging rate of 4 image stacks per s. Image analysis was performed as previously described[34], using a custom written pipeline in MatLab (MathWorks, Inc.). After analysis, $WHaloCaMP1a_{669}$ activity was visualized by creating a multi-channel composite image of the ΔF/F time-series images, which were superimposed on a static anatomical reference image. The anatomical reference image was generated by computing a projection of the ΔF/F images along the z- and time-axes and applying a CLAHE filter with block size of 15, bins 255, max gradient 4.

**Multiplexed light sheet imaging of far-red, red, and green fluorescence in zebrafish larvae**

Multiplexed light sheet single plane functional imaging was performed on a Zeiss Lightsheet Z.1 microscope with two a pco.edge 5.5 sCMOS cameras (PCO). For single cells resolution imaging of neuronal and astrocyte activity, fish were paralyzed in a drop of 1 mg $ml^{-1}$ alpha-bungarotoxin solution (Invitrogen) in system water for 30 s, before recovering in system water for 5 minutes. This paralysis was not performed for three color multiplexed imaging of Tg(*elavl3*:NES-WHaloCaMP1a) x Tg(*actb2*:iGlucoSnFR; *acta1a*:jRGECO1a) to retain the muscle calcium transients. Fish were mounted in 2% low melting agarose in system water in a glass capillary, and the solidified agarose extruded into an imaging chamber filled with system water for imaging. For imaging with 4-aminopyridine (4-AP), a solution of 8 mM was prepared in system water, which was diluted into the imaging chamber before imaging. Final concentration of 4-AP was 500-700 nM.

Images were acquired using Objective W Plan-Apochromat 10x/0.5 M27 75mm illumination objective and two 5x/0.1 (with a correction ring) for side illumination. As we saw artifacts on dual-side illumination, only one side of illumination was used during acquisition. Pivot scan was always on during acquisition. Before each imaging session, the cameras were manually aligned, and the light sheet was corrected using the correction rings on the 5x illumination objectives. Single plane light sheet imaging was performed with 488 nm, and 561 nm and 638 nm excitation laser lines and a laser blocking filter (Filter Module LBF 405/488/561/638) was used in the excitation light path. Exposure time was 234.69 ms.

For three color functional imaging of Tg(*elavl3*:NES-WHaloCaMP1a) xTg(*actb2*:iGlucoSnFR; *acta1a*:jRGECO1a), two tracks were used with a filter change between the track acquisitions. This filter change decreased the imaging been to 5s per frame. In track 1, 488 nm and 638 nm excitation were used with a 560 nm beam splitter. Emission filter 505-545 nm was in front of camera 1, and longpass 660 nm filter in front of camera 2. In track 2, 561 nm excitation light was used with 640 nm beam splitter and 575-615 nm emission filter in front of camera 1. Images were acquired at 1×zoom for up to 60 min at 1000 x 1800 pixels. Images where registered using suite2p[36] before $\Delta F/F_0$ of ROIs determined in FIJI[17] were calculated using a rolling average of 250 frames with of lowest 0.1 quantile used for $F_0$.

For dual color imaging with Tg(*elavl3*:NES-WHaloCaMP1a-EGFP) x Tg(*gfap*:jRGECO1b and Tg(*elavl3*:NES-WHaloCaMP1a-EGFP) x Tg(*elavl3*:jRGECO1b) anatomical images were first acquired with single fluorophore excitation of: EGFP, 488 nm excitation, 560 nm beam splitter, 505-545 nm emission filter; jRGECO1b, 561 nm excitation, 640 nm beam splitter, 575-615 nm emission filter; WHaloCaMP1a$_{669}$, 638 nm excitation, 560 nm beam splitter, longpass 660 nm emission filter. Functional imaging was performed with simultaneous 561 nm and 638 nm excitation light, a 560 nm beam splitter and 575-615 nm emission filter in front of camera 1 and longpass 660 nm emission filter in front of camera 2. The 561 nm was reduced to the lowest amount to limit bleedthrough of the jRGECO1b signal to the far-red emission channel. Images were acquired at 2.5×zoom at 1960 x 1960 pixel size, 354.4 µm x 354.4 µm, 0.18 µm per pixel.

Time series of 1500 frames were collected. Time series were concatenated together when longer analysis was desired. Images were downscaled for analysis.

**Image analysis of single plane light sheet neuronal and astrocyte imaging**

For analysis dual color functional imaging of WHaloCaMP1a$_{669}$ in neurons and jRGECO1b in astrocytes a combination of FIJI[17], suite2p[36], Cellpose[37], and Rastermap[38] was used. The two channels were analyzed separately. First suite2p was used for registration of the images, and cell segmentation was performed using Cellpose seeding with a manually determined diameter. Suite2p was then used to extract the traces from the segmented cells. The $\Delta F/F_0$ was calculated using a rolling average of 100 frames (for *elavl3*:jRGECO1b acquisitions) or 200 frames (for *gfap*:jRGECO1b acquisitions) with $F_0$ calculated from the lowest 0.1 quantile. Fluorescent traces were clustered with Rastermap before being presented as heatmaps.

**Fluorescence lifetime image microscopy (FLIM)**

Fluorescence lifetime image microscopy was performed on a STELLARIS 8 FALCON microscope (Leica Microsystems), with a White Light Laser with tunable excitation wavelengths 440–790 nm operating at 80 MHz. For full lifetime decay calculations, the microscope was set to operate at 40 MHz. A 20× water immersion objective (Leica, HC PL IRAPO, NA = 0.75) was used for all imaging. HyD photon-counting detectors were used.

**FLIM in purified protein**

In order to generate a calibration curve of lifetime vs. [$Ca^{2+}$] in purified protein, WHaloCaMP1a was labeled with a dye-ligand and diluted into solutions made by mixing different proportions of concentrations made from Calcium Calibration Buffer Kit #1 (Invitrogen). A small drop was placed on a microscope slide, and a double adhesive divider was placed on the slide, before being covered with a coverslip (#1.5). For WHaloCaMP1a$_{494}$ excitation was 488 nm, and emission collected on a HyDX2 set to 500-575 nm. For WHaloCaMP$_{552}$ excitation was 561 nm, and emission collected on a HyDS3 set to 575-650 nm. For WHaloCaMP$_{669}$ excitation was 671 nm, and emission collected on a HyDX4 set to 680-780 nm. For $WHaloCaMP_{722}$ excitation was 730 nm, and emission collected on a HyDR5 set to 745-850 nm.

**FLIM in tissue cultured cells**

Janelia Cell Culture Facility performed tissue culture and regular mycoplasma testing. HeLa cells (CCL-2) were purchased from ATCC and were cultured in Dulbecco's modified Eagle medium (DMEM, phenol red-free; Life Technologies) supplemented with 10% v/v fetal bovine serum (FBS, Life Technologies), 1 mM GlutaMAX (Life Technologies). Cells were cultured in 37 °C in a humidified 5% (v/v) $CO_2$ environment. Cells were transiently transfected by nucleofection (Lonza) 24 hours before imaging, and plated in 35 mm dishes (Mattek, #1.5 coverslip). Before imaging, cells were labeled with 50 nM $JF_{669}$-HaloTag ligand in growth media, before being washed 2x with imaging buffer containing 145 mM NaCl, 2.5 mM KCl, 10 mM glucose, 10 mM HEPES, pH 7.4, 2 mM $CaCl_2$ and 1 mM $MgCl_2$ for histamine stimulation experiments or Hanks' Balanced Salt solution (HBSS) with no calcium, no magnesium, no phenol red (ThermoFisher Scientific, 14175095) for calcium titration experiments. Images were collected at 1600 x 1600 pixels, pinhole set to 1 Airy Unit (AU), and 2×zoom. Images were collected every 5 s for 10 minutes.

For cellular calcium titration experiments, $[Ca^{2+}]$-free HBSS was replaced by solutions of $[Ca^{2+}]_{free}$ concentrations made from Calcium Calibration Buffer Kit #1 (Invitrogen) with zero and 10 mM CaEGTA, and equilibrated for 10 min at 37 °C. Digitonin (Promega, G944A) was then used to permeabilize the cells. The digitonin stock was diluted to 2 mg $mL^{-1}$ in buffer and 1 µL added to the 2 mL volume in the imaging dish immediately before acquisition, giving a final concentration of 1 µM.

For histamine stimulation experiments, histamine was diluted in imaging buffer to 100 mM stock solution. This was further diluted by adding 20 µL to the 2 mL imaging buffer on the cells, to give a final concentration of 1 µM.

**FLIM of neurons in zebrafish larvae**

Zebrafish larvae, Tg(*elavl3*:NES-WHaloCaMP1a-EGFP) 4-5 d.p.f. were labeled and paralyzed as described above. The fish were then mounted, with the brain closest to the coverslip, in 2% low melting agarose in a 35 mm dish (Mattek, #1.5 coverslip). The fish were imaged in system water with 6×zoom, pinhole set to 1 AU and 512x 512 pixels. The imaging rate was 1 frame per s.

**FLIM image analysis**

Fluorescence lifetime fitting was performed in the LASX FLIM software. A spatial binning of 2 was performed, and each time point was split in the software. A three-component fit was used to fit the fluorescence lifetime. The amplitude weighted fluorescence lifetime for the whole image is reported for WHaloCaMP1a labeled with different dye-ligands and imaged at 40 MHz. For quantifying concentrations of [$Ca^{2+}$] in HeLa cells and neurons of zebrafish larvae each pixel in the image to a decay curve. The fit was performed on images that had a threshold of 20 – 50 counts for each pixel. The image was then exported to a TIFF file with a 0.001 scaling factor on the lifetime and analyzed in FIJI[17]. A calibration curve from the purified protein or digitonin lysed cells was fit using a four-parameter dose-response curve (variable slope) using GraphPad Prism software. (a) is the value of fluorescence at the bottom of the curve, (b) is the value of fluorescence at the top of the curve, ($EC_{50}$) is the concentration of agonist that gives a response halfway between bottom and the top, and (h) is the hill or cooperative coefficient.

$$y = a + \frac{x^h \times (b-a)}{x^h + {EC_{50}}^h} \qquad\qquad x = EC_{50}\left(\frac{a-y}{y-b}\right)^{\left(\frac{1}{h}\right)}$$

To calculate the concentration of $Ca^{2+}$ in the cell, the equation was rearranged, and a value of x = [$Ca^{2+}$] was calculated for each measured value of y (lifetime (ns)) for an ROI in a time series. The intensity channel was also exported from the LASX FLIM software, and the ΔF/F was calculated from the 1st quartile for the time series. For visualization purposes for the lifetime time series, a gaussian filter of 0.8 was performed. All quantitative analysis was performed on the raw exported images before any image processing.
