## Supplementary Note 1 for "A modular chemigenetic calcium indicator enables in vivo functional imaging with near-infrared light"

### The $K_{L-Z}$

Standard rhodamines that have an *ortho*-carboxyl group on the pendant phenyl ring can exist in two forms in equilibrium (**Fig. 1a**). The lactone form (**L**) occurs when the carboxyl group forms a bond with the central carbon atom in an intramolecular reaction, forming a lactone. In this uncharged structure, the two aryl groups are independent, yielding a colorless and nonfluorescent molecule. This is in equilibrium with the zwtterionic form (**Z**), where the lactone bond is broken, yielding a positively charged xanthylium or xanthylium-like species and a carboxylate moiety. The extended conjugated system in the zwitterionic form gives rise to the strong visible absorption and fluorescence properties that are characteristic of rhodamine dyes.

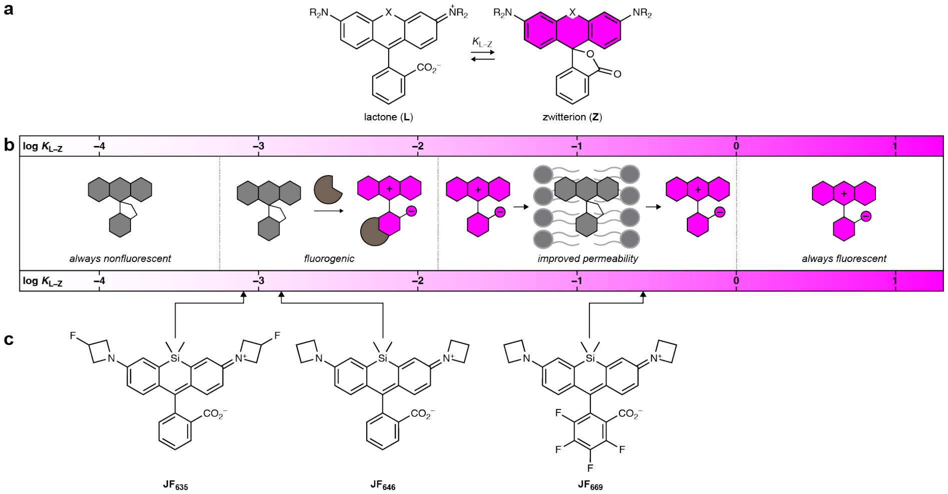


**Figure 1. The lactone–zwitterion equilibrium constant ($K_{L-Z}$) predicts performance of rhodamine dyes in biological systems.** (**a**). General rhodamine structure showing the equilibrium between the lipophilic colorless lactone (**L**) and fluorescent zwitterion (**Z**). (**b**) Framework relating the $K_{L-Z}$ to performance of rhodamine dyes. (**c**) Structures of Si-rhodamine dyes **JF635**, **JF646**, and **JF669**.

The dynamic equilibrium between the lactone and zwitterionic forms of rhodamine dyes is dependent on both the environment around the dye and the chemical structure of the fluorophore. Differences in the lactone–zwitterion equilibrium can be measured in various ways, for example by dioxane-water titrations used in various works,[1-5] or by measuring the $K_{L-Z}$[6-8].

$K_{L-Z}$ is calculated using equation 1:

$$K_{L-Z} = \frac{\varepsilon_{dw}/\varepsilon_{max}}{1 - \varepsilon_{dw}/\varepsilon_{max}}$$

Where is $\varepsilon_{dw}$ is the molar extinction coefficient when the dye is in a 1:1 (v/v) dioxane:water mixture. $\varepsilon_{max}$ is the maximum molar extinction coefficient measured in either 0.1% (v/v) trifluoroacetic acid (TFA) in ethanol or 0.1% (v/v) TFA in 2,2,2-trifluoroethanol (TFE) depending on dye type. These strongly acidic conditions presumably protonate the *ortho*-carboxyl group; the resulting cationic species is a proxy for the zwitterionic form allowing estimation of the absorptivity of pure zwitterion and subsequent calculation of $K_{L\text{–}Z}$.

We previously developed a general framework correlating $K_{L\text{–}Z}$ with the performance of dyes in biological environments[6-8] (**Fig. 1b**). We found that both fluorogenicity—dyes that turn on when they bind their cognate biomolecular target—and bioavailability strongly depend on $K_{L\text{–}Z}$. HaloTag ligands based on dyes with $4\times10^{-4} < K_{L\text{–}Z} < 9\times10^{-3}$ are typically highly fluorogenic. Dyes with $9\times10^{-3} < K_{L\text{–}Z} < 1$ usually show high cell and tissue permeability. An example is the Si-rhodamine series **JF$_{646}$**, **JF$_{635}$**, and **JF$_{669}$** (**Fig. 1c**). **JF$_{646}$** exhibits a $K_{L\text{–}Z}$ = 0.0014, which puts it squarely in the fluorogenic region (**Fig. 1b**); **JF$_{646}$-HaloTag ligand** shows modest fluorogenicity with a ~20-fold increase in absorption and fluorescence intensity upon binding the HaloTag protein.[5] Addition of a single fluorine atom on each azetidine auxochrome to give **JF$_{635}$** substantially decreases $K_{L\text{–}Z}$ = $6\times10^{-4}$; this leads to a higher degree of fluorogenicity for the **JF$_{635}$-HaloTag ligand** (~200-fold).[6] Installation of fluorine atoms on the pendant phenyl ring to give **JF$_{669}$** increases the $K_{L\text{–}Z}$ = 0.262; the **JF$_{669}$-HaloTag ligand** shows only modest fluorogenicity but high bioavailability with the ability to cross the blood–brain barrier after *intra venous* administration.[7] This rubric is also applicable for standard oxygen-containing rhodamines and carborhodamines.[7] The example of **JF$_{646}$**, **JF$_{635}$**, and **JF$_{669}$** demonstrates the challenge of creating a fluorophore that is both fluorogenic and highly bioavailable, since both attributes depend on a single property.

The $K_{L\text{–}Z}$ parameter is not perfect and we note several criticisms. First, the measurement is done in a mixture of dioxane and water. Although this gives a wide range of values that show clear trends with bioavailability and fluorogenicity, these conditions are not biologically relevant; it is difficult to relate the measured $K_{L\text{–}Z}$ to the dye state in a biological environment. Second, we measure the $K_{L\text{–}Z}$ of parent dyes, and not conjugated to functional handles for example the chloroalkane in the HaloTag Ligand. Functional handles can affect this equilibrium, making this parameter useful in comparing compounds within a specific class (*e.g.*, HaloTag ligands) but not across different compound types. Third, other properties affect cell-permeability. For example, the phosphine oxide dye **JF$_{722}$** exhibits a $K_{L\text{–}Z}$ = 0.026, which should indicate high bioavailability for HaloTag ligand derivatives.[7] The **JF$_{722}$-HaloTag ligand** does not show efficient labeling in animals models, however, presumably due to the additional lipophilic phenyl group on the fluorophore structure.

In summary, the $K_{L\text{–}Z}$ is a useful predictor of the performance of dye conjugates in biological environments. The relationship between $K_{L\text{–}Z}$ and bioavailability and fluorogenicity provides a straightforward framework for optimization of dye ligands within a structural class[7]. For chemigenetic sensors, however, the dependence of both fluorogenicity and bioavailability on a single property is problematic. Previous sensors, such as HaloCaMP,[9] relied on fluorogenic dye ligands derived from fluorophores with relatively low $K_{L\text{–}Z}$ values. Although these dyes show robust fluorescence changes in response to changes in the environment, they did not exhibit good bioavailability in the central nervous system in animal models, limiting the ability to allow functional imaging in the central nervous system with single cell resolution. In this work, we focus instead on building chemigenetic $Ca^{2+}$ sensors around dye ligands where the dye has higher $K_{L\text{-}Z}$ values, such as **JF$_{669}$**. As **JF$_{669}$-HaloTag ligand** shows only modest fluorogenicity we needed to explore a different way of modulating the fluorescence emission – here we use a strategically placed tryptophan, which can quench the fluorescence of the dye in a $Ca^{2+}$ dependent manner.
