## Supplementary Video Captions for "A modular chemigenetic calcium indicator enables in vivo functional imaging with near-infrared light"

**Supplementary Movie 1. Volumetric SiMView light sheet imaging of WHaloCaMP1a$_{669}$ in zebrafish larva**
Volumetric imaging in the near infrared of neuronal activity in Tg(*elavl3*:NES-WHaloCaMP1a-EGFP) labeled with JF$_{669}$-HaloTag ligand 5 days post fertilization (d. p. f.) The zebrafish larva was paralyzed with alpha bungarotoxin. Imaged at 4 volumes per second. Displayed as $\Delta F/F_0$ (red-hot color look-up table) superimposed on anatomy (grey). Scale bar, 100 µm. Time stamp in min:s.

**Supplementary Movie 2. Three color functional multiplexed imaging of WHaloCaMP1a$_{669}$, jRGECO1a and iGlucoSnFR in zebrafish larva**
Single plane light sheet imaging of Tg(*elavl3*:NES-WHaloCaMP1a, *actb2*:iGlucoSnFR; *acta1a*:jRGECO1a) labeled with JF$_{669}$-HaloTag ligand at 5 days post fertilization (d. p. f.). Imaged at 0.2 Hz. Three colors are displayed in raw fluorescence together with an overlay of fluorescence of all three channels.

**Supplementary Movie 3. Dual color imaging of WHaloCaMP1a$_{669}$ in neurons and jRGECO1b in astrocytes in zebrafish larva**
Single plane light sheet imaging of Tg(*elavl3*:NES-WHaloCaMP1a-EGFP, *gfap*:jRGECO1b) labeled with JF$_{669}$-HaloTag at 5 days post fertilization (d. p. f.). The zebrafish larva was paralyzed with alpha bungarotoxin, imaged at 4 Hz. Each channel is displayed as raw fluorescence as greyscale with the insert showing the overlay of WHaloCaMP1a$_{669}$ in magenta and jRGECO1b in yellow. Time stamp in min:s.

**Supplementary Movie 4. Dual color imaging of WHaloCaMP1a$_{669}$ in neurons and jRGECO1b in astrocytes during a seizure-like state induced by 4-aminopyridine**
Single plane light sheet imaging of Tg(*elavl3*:NES-WHaloCaMP1a-EGFP, *gfap*:jRGECO1b) labeled with JF$_{669}$-HaloTag at 5 days post fertilization (d. p. f.), and treated with 4-aminopyridine. The zebrafish larva was paralyzed with alpha bungarotoxin. Imaged at 4 Hz. Each channel is displayed as raw fluorescence as greyscale with the insert showing the overlay of WHaloCaMP1a$_{669}$ in magenta and jRGECO1b in yellow. Three time series of 6.2 min are concatenated. Time stamp in min:s.

**Supplementary Movie 5. FLIM of WHaloCaMP1a$_{669}$ expressed in HeLa cells stimulated by histamine**
Raw fluorescence intensity (left) and FLIM (right) of WHaloCaMP1a labeled with JF$_{669}$-HTL in HeLa cells stimulated by histamine (1 µM). Lifetime color bar is scaled x 1000 ns. One frame was collected every 5 s. Time stamp in min:s. Still images and $\Delta F/F_0$ or $[Ca^{2+}]_{calc}$ from a calibration curve of lifetime in Fig. 5d.

**Supplementary Movie 6. FLIM of WHaloCaMP1a$_{669}$ in the forebrain of zebrafish larva**
Overlaid near infrared intensity and lifetime using the LASX Leica FLIM software of Tg(*elavl3*:NES-WHaloCaMP1a-EGFP) labeled with JF$_{669}$-HTL. The zebrafish larva was paralyzed

with alpha bungarotoxin and imaged at 1 Hz. Time stamp in min:s. Still images and $\Delta F/F_0$ or $[Ca^{2+}]_{calc}$ from a calibration curve of lifetime in Fig. 5f.
